# Knowledge-guided data mining on the standardized architecture of NRPS: subtypes, novel motifs, and sequence entanglements

**DOI:** 10.1101/2022.03.14.484258

**Authors:** Ruolin He, Jinyu Zhang, Yuanzhe Shao, Shaohua Gu, Chen Song, Long Qian, Wen-Bing Yin, Zhiyuan Li

**Affiliations:** Center for Quantitative Biology, Academy for Advanced Interdisciplinary Studies, Peking University, Beijing, 100871, China; State Key Laboratory of Mycology, Institute of Microbiology, Chinese Academy of Sciences, Beijing 100101, PR China; Savaid Medical School, University of Chinese Academy of Sciences, Beijing, 100049, PR China; Peking-Tsinghua Center for Life Sciences, Academy for Advanced Interdisciplinary Studies, Peking University, Beijing, 100871, China

## Abstract

Non-ribosomal peptide synthetase (NRPS) is a diverse family of biosynthetic enzymes for the assembly of bioactive peptides. Despite advances in microbial sequencing, the lack of a consistent standard for annotating NRPS domains and modules has made data-driven discoveries challenging. To address this, we introduced a standardized architecture for NRPS, by using known conserved motifs to partition typical domains. This motif-and-intermotif standardization allowed for systematic evaluations of sequence properties from a large number of NRPS pathways, resulting in the most comprehensive cross-kingdom C domain subtype classifications to date, as well as the discovery and experimental validation of novel conserved motifs with functional significance. Furthermore, our coevolution analysis revealed important barriers associated with reengineering NRPSs and uncovered the entanglement between phylogeny and substrate specificity in NRPS sequences. Our findings provide a comprehensive and statistically insightful analysis of NRPS sequences, opening avenues for future data-driven discoveries.

**Author Summary:** NRPS, a gigantic enzyme that produces diverse microbial secondary metabolites, provides a rich source for important medical products including antibiotics. Despite the extensive knowledge gained about its structure and the large amount of sequencing data available, the frequent failure of reengineering NRPS in synthetic biology highlights the fact that much is still unknown. In this work, we applied existing knowledge to data mining of NRPS sequences, using well-known conserved motifs to partition NRPS sequences into motif-intermotif architectures. This standardization allows for integrating large amounts of sequences from different sources, providing a comprehensive overview of NRPSs across different kingdoms. Our findings included new C domain subtypes, novel conserved motifs with implication in structural flexibility, and insights into why NRPSs are so difficult to reengineer. To facilitate researchers in related fields, we constructed an online platform “NRPS Motif Finder” for parsing the motif-and-intermotif architecture and C domain subtype classification (http://www.bdainformatics.org/page?type=NRPSMotifFinder). We believe that this knowledge-guided approach not only advances our understanding of NRPSs but also provides a useful methodology for data mining in large-scale biological sequences.

## Introduction

Non-ribosomal peptide (NRP) synthesized by non-ribosomal peptide synthetase (NRPS) are a diverse family of natural products widespread in fungi and bacteria^1–4^. According to the BiG-FAM database (as of 2022/8), NRPS is the most prevalent class (29.30%) out of the 1,060,938 biosynthetic gene clusters (BGCs) detected from 170,585 bacterial genomes, and 46.81% out of the 123,939 BGCs from 5,588 fungal genomes^5^. Microbes utilize NRP for various niche-construction activities, such as communication^6, 7^, defense^8–12^ and resource-scavenging^13^. Diverse NRP serve as reservoirs for pharmaceutical innovations, with anti-cancer, anti-bacterial, anti-fungal, anti-viral, cytotoxic, and immunosuppressive activities^14–17^. To date, nearly thirty NRP structural medicines have been approved for commercialization^14^, including actinomycin^18^, penicillin^19, 20^ and vancomycin^21^.

Many reengineering efforts have been inspired by the pharmacological potentials of NRPS, as well as its modular architecture^22, 23^. NRPS synthesizes peptides in an assembly-line fashion using its repeating module units. Each module is generally comprised of three key domains, including the adenylation (A) domain, which recognizes the substrate and activates it as its aminoacyl adenylate; the condensation (C) domain, which catalyzes amide bond formation between two substrates; and the thiolation (T) domain, which shuttles the substrates and peptide intermediates between catalytic domains. Other optional domains are also important, such as the thioesterase (TE) domain or terminal condensation-like (C_T_) domain^24^, which cyclize or release peptide from the NRPS and generally occurs at the end of a NRPS; epimerization (E) domain, which catalyzes conversion of L-amino acids substrate to D-amino acids and usually follows the C domain in case it occurs. In order to create novel products, many synthetic efforts have been made to re-arrange these building blocks of NRPSs. For example, due to the central roles of A domains in substrate recognition and activation, early works focused on the A domain, including A domain substitution, reprogramming A domain substrate coding residues, and A subdomain substitution^22^. Later, considering possible roles of C domains in substrate selectivity, researchers attempted to re-engineer catalytic domains together with the A domain, such as substituting a complete module, or the C+A domain and T+C+A domain^22^. Recently, Bozhüyük et al. advanced the exchange unit concept, and recombined NRPS at a fusion point between the C and A domains which they named as “C-A linker”^25^, and they also found another cutting point inside the C domain^26^. Calcott et al. also attempted to recombine at another cutting point between the C and A domains^27^, which was almost located at the start of the C-A linker defined by Bozhüyük et al. However, in most cases, reengineering often substantially reduce the product yield. Furthermore, conversions of these units were typically performed between two substrates with similar chemical properties, based on well-studied NRPS systems with a small number of sequences^28, 29^. A systematic understanding towards the rational design of NRPS has yet to be developed.

The massive expansion of microbial sequencing data has created opportunities towards a comprehensive understanding of NRPSs^30^. As of October 2021, the Integrated Microbial Genomes Atlas of Biosynthetic gene Clusters (IMG-ABC) v5 database contains a total of 411,027 BGCs based on sequence annotation^31^, and Minimum Information about a Biosynthetic Gene Cluster (MiBiG) v2 database contains 1926 BGCs confirmed to be biologically active^32^. Together with accelerated expansion of databases, bioinformatics tools and platforms that specifically predict and annotate the secondary metabolites are also in rapid development^33^, such as antiSMASH^34^, PRISM^35^, SeMPI^36^. Pfam, a widely utilized database of protein families and domains since 1997^37, 38^, has also been commonly used for NRPS domain detection by profile hidden markov models (HMM). They have been used to aid in the discovery of new NRPS antibiotics^15, 39^. However, different bioinformatics tools and platforms employ different annotation standards and algorithms, and their definitions of domains and inter-domains in NRPS are perplexing.

Aside from annotating the module-domain architecture, predicting the substrate specificity of A domains in NRPS has been a focus of research for many years, with various algorithms being developed to address the challenge^36, 40–43^. Some of these methods use supervised learning and rely on a training set of full-length A domain sequences and their known substrates^41^. Other well-known approaches incorporate structural information of the A domain^44^. With advances in protein structures, the substrate-specifying residues of modular mega-enzymes, such as NRPS^28, 45^ and polyketide synthase (PKS)^46^, were identified. Based on the first crystal structure of the A domain (PheA, PDB 1AMU)^28^, Stachelhaus et al. identified ten substrate-specifying residues in the A domain’s binding pocket. These ten residues, known as the “specificity-conferring code”, strongly associate with the substrate specificity, and have been widely exploited by researchers to improve prediction efficiency^44^. However, the diversity of the specificity-conferring code for the same substrate remains a challenge, making the performance of current machine learning algorithms heavily dependent on the training set. Actually, diverse forms of the specificity-conferring code could exist for the same substrate^44^. Such diversity of the specificity-conferring code causes barriers in extending the prediction algorithm: without a one-to-one mapping between the specificity-conferring code and substrate specificity, the performance of current machine learning algorithms heavily replies on the training set. Actually, performances of these algorithms in fungal NRPSs are unsatisfactory, as fungi have a significantly smaller number of A domains with known substrate. Furthermore, reengineering based on modifying the specificity-conferring code does not operate as expected^45^, highlighting the complexities in the A domain’s function.

Coevolution analysis has been used as a general strategy to uncovering protein interactions and functional couplings from a large number of sequences^47^. Existing methods in coevolution analysis include normalized mutual information (MI) of amino acid frequencies^48^, direct coupling analysis (DCA)^49^, protein sparse inverse covariance (PSICOV)^50^ and statistical coupling analysis (SCA)^51, 52^. Among them, SCA has been utilized to identify appropriate boundaries for protein reengineering^53^. The basic idea of SCA is to identify groups of residues sharing the same modes of covariations, which are referred to as “sectors”^54^. Residues projecting to a single sector can be viewed as “evolving together”, and are likely to cooperate in function^55^. Therefore, sectors could serve as the “evolutionary units” for reengineering, as the borders of sectors provide rational cutting-points for preserving functional connections between residues^54^.

Applying SCA to NRPS sequences to gain insights on reengineering is a straightforward idea. However, one obstacle to this simple notion is the paucity of manually curated sequences: the majority of sequences in databases were automatically annotated by machine learning algorithms, and have not yet been reviewed by human experts. Moreover, different algorithms with varying standards for annotation were adopted by different researchers, which muddled the exact definitions of domains and interdomains in NRPSs. For examples, A and C domains annotated by the default option of antiSMASH are substantially shorter than Pfam-annotated domains^37^, and usually different with sequences from the Protein Data Bank (PDB)^56–58^. Even the domain annotations from antiSMASH, one of the most popular annotation platforms for NPRSs, slight changes happened between versions.

One solution for this “confusion by the annotation” is to focus on the invariant sequence pieces in NRPSs architecture, known as “core motifs”^59^. Despite the enormous diversity in NRPs, highly conserved sequence elements have been discovered across domains, and are usually essential to the domain function. These well-established core motifs include: 7 motifs in the C domain, 10 motifs in the A domain (of note, these are not the specificity-conferring code), 7 motifs in the E domain, 1 motif in the T domain and 1 motif in TE domain^59^. However, these motifs have been proposed for more than two decades. Since then, these motifs have not been systematically updated or confirmed. These conserved motifs could be used to standardize NRPS domains, allowing for statistical analysis of NRPS sequences from large databases without manual curation.

In our work, we constructed a “coordinate system” for NRPSs based on well-known conserved core motifs. Such motif-and-intermotif architecture allowed us to integrate sequence information of NRPSs from multiple databases and extract global statistics on the length, variation, and functional properties of each NRPS sequence. Following that, we were able to systematically evaluate a large number of sequences for their statistical properties and function-sequence couplings. For example, we identified new C domain subtypes and obtained the most comprehensive cross-kingdom classification of C domains to date. In addition, we uncovered several previously unrecognized conserved motifs, which could have functional implications as well as serve as candidates for novel core motifs. One of these novel motifs is located close to the A domains specificity pocket, with implications in structural flexibility, the importance of which was then supported by mutation experiments. Then, using coevolution analysis, we dissected two major challenges in reengineering NRPSs: first, there is no cutting point to separate multiple overlapping coevolving sectors, limiting domain/subdomain swapping; second, the specificity of the A domain intertwines with not only the specificity-conferring code but also the length of five loops in the A3-G intermotif, complicating the rational modification of A domain. Our findings provide a comprehensive overview and statistical insights into the sequence features of NRPSs, paving the way for future data-driven discoveries. In the end, an online platform entitled “NRPS Motif Finder” is offered for parsing the motif-and-intermotif architecture and C domain subtype classification (http://www.bdainformatics.org/page?type=NRPSMotifFinder).

## Result

### 1. Overview of the motif-and-intermotif architecture of NRPS

#### 1.1. Sequence properties of motifs and intermotifs in NRPS domains

In order to establish the foundation of our study, we first partitioned the NRPS architecture based on well-known core motifs. The sequence representations of 19 well-studied motifs were curated from literature^59^, then located to 7,329 NRPS domains from MiBiG database by alignment (see Method for details). As expected, most of the known core motifs are highly conserved, exhibiting clear sequence logos (first panel, Fig 1A-1D, Fig S1-S5). Motifs from the A, E, and T domains have sequence identities higher than 60%. However, C domain motifs exhibit a lower sequence identity compared to A, T and E domains. Among the C motifs, motif C3 is the most conserved, with 55% sequence identity across all C subtypes (Fig 1A, first panel). This is reasonable, as C3 has been reported to be critical for catalysis^60, 61^. With the exception of the Heterocyclization subtype, where the motif C3 is DxxxxD, the motif C3 in other subtypes is almost always HHxxxD. Motifs C6 and C7 are less conserved, with identities of 22% and 28% respectively. This lower conservation in C motifs is likely due to the presence of various functional subtypes within the C domain^60^, which we will discuss in further detail in the following sections 1.3-1.5.

**Figure 1.**
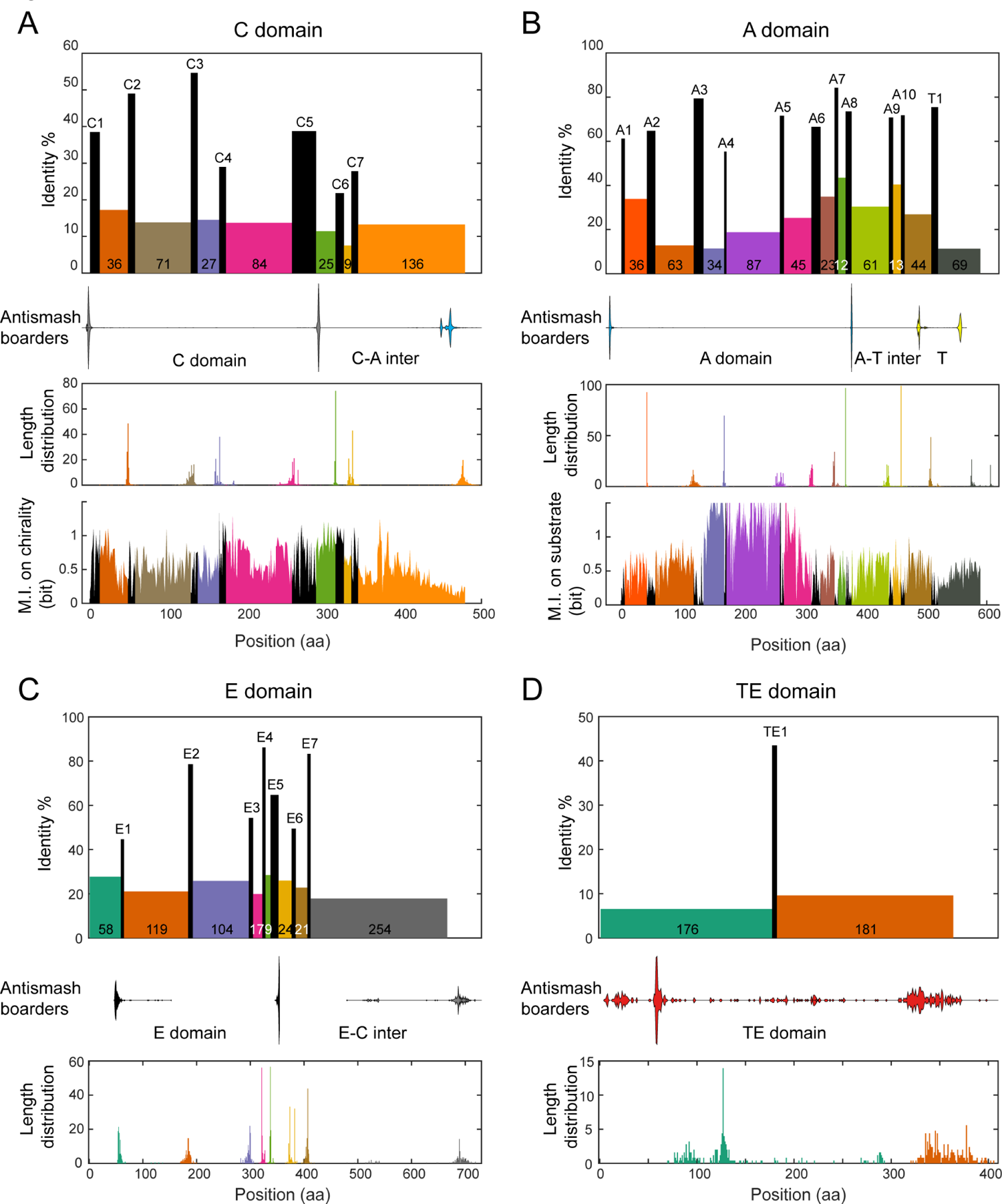
Overview of the motif-and-intermotif architecture of canonical NRPS domains. **A.** Organization of C domain. Rectangles in the first panel show the seven C motifs (black) and eight inter-motif regions (colored), including the last one as the inter-motif region between the C7 and A1 motifs. The heights of the rectangles indicate the average sequence identities. The widths of the rectangles are their average lengths in amino acid sequences, with the average lengths of inter-motif regions printed on their rectangles. Violin plots in the second panel mark the border positions of domains annotated by antiSMASH 5.1 by the default option (the start and end borders of the C domain: gray; the start of the A domain: blue). The third panel shows the distributions of the ending positions of the eight inter-motif regions, counting from their preceding conserved motifs. The fourth panel shows the mutual information (M.I.) between residues in the C domain and chirality subtypes. **B.** Same as that in **A**, but for the A domain and the T domain. It starts with the A1 motif and ends before the C1 motif. The fourth panel shows the mutual information between residues and A domain substrate specificity (see Method for details). **C.** Same as that in **A**, but for the E domain. It starts after the T1 motif, and ends before the C1 motif. **D.** Same as that in **A**, but for the TE domain. It starts after the T1 motifs, and ends with regions after the annotated TE domain.

The sequence region between the conserved motifs, referred to as the “intermotif region”, has a comparatively lower level of sequence identity, with values ranging from 10%-25%. The majority of the intermotif regions display narrow length distributions, with a standard deviation smaller than 10 aa (third panel, Fig 1A-1C), demonstrating the architecture invariance of NRPSs. However, the TE motif at the end of the cluster exhibits a much wider distribution (320-380aa, Fig 1D). An additional noteworthy exception is observed in the intermotif region between the motif T1 and its next motif C1, which exhibits a bi-modal distribution (Fig 1B). Upon closer examination of the domain annotations, it was found that the length of the T1-C1 region depends on the MiBiG annotation of the following C domain (Fig S3B): Specifically, regions followed by the LCL subtype tend to be 65-70 amino acids in length, whereas regions followed by the Dual subtype are typically 95-105 amino acids in length. These observations suggest that the architecture of intermotif regions may have functional implications. Its worth mentioning that by default, antiSMASH annotation s for the C domain do not include the motif C6 and C7; these for the A domain do not include the motif A9 and A10; and these for the E domain do not include the motif E6 and E7 (second panel, Fig 1A-1C). This explains the difference in NRPS domain length between MiBiG and Pfam (Fig S6).

#### 1.2. Exploring the coupling between sequence and function in the C and A domains by mutual information

Another focus of this study is how sequences couple with functions. Conserved motifs serve as “anchors” in multiple sequence alignment, facilitating a detailed evaluation of information content. Given that the sequences of the C domain have been previously demonstrated to be indicative of chirality subtypes^60^, we first quantified the Shannon mutual information (MI) between the aligned sequences and functional subtypes in the C domain, residue by residue (see Method for details). The region between the C1 and C7 motifs, represented in the last panel of Figure 1A, contains a substantial amount of information about chirality subtypes. After the C7 motif, the information about chirality decreases gradually until it reaches the baseline level as the sequence progresses toward the A1 motif (Fig 1A, last panel).

Generally, conserved motifs contain low levels of mutual information (Fig 1). Nevertheless, it’s noted that the C5-C6 region, including the C5 and C6 motif, has a high MI value with C domain subtypes (Fig 1A, last panel). We checked this region in the structure of LgrA, the only structure that contains multi-module NRPS in multiple conformations so far characterized by Reimer et al.^62^. We noticed that this region is located at the interface between the donor T domain and the recipient C domain of the next module (T_n_:C_n+1_), which has been suggested to play roles in chirality^63^. Multiple coevolution positions between the T_n_ domain and C_n+1_ domain were suggested by DCA in Reimer et al.’s work, with multiple key residues located in the C5 and C6 motif. Therefore, T_n_ and C_n+1_ are functionally linked by the C5-C6 motif in the T_n_:C_n+1_ interface, which explains the high MI value between C5-C6 intermotif region and C domain subtypes.

In light of the critical role played by the A domain in substrate selectivity, we also employed the same mutual information measurement to evaluate the relationship between each position in the A domain and substrate specificity (see Method for details). Results depicted in the last panel of Figure 1B show that positions with high information content start from the A3 motif and end before the A6 motif. This region overlaps the substrates binding pocket^56^. It is worth noting that the A4-A5 region has been well-known for containing the specificity-conferring code^28^.

#### 1.3. Cross-kingdom validation and comparison of motif properties in a larger dataset

The MiBiG database provides a solid foundation for the discovery of the fundamental properties of NRPSs, as all BGCs in the database have been confirmed to be active. Then, to further validate the conservation of the NRPS motifs, we expanded our analysis to a larger dataset. With that aim in mind, we downloaded all complete bacterial genomes (30,984) and all assembly-level (including “contig”, “scaffold”, “chromosome”, and “complete” levels) fungal genomes (3,672) from NCBI. After removing genomes with 99.6% similarity, we had 16,820 bacterial and 2,505 fungal genomes (as of 2022/10/23). BGCs in these genomes were annotated using antiSMASH v6. Unlike the MiBiG database, where the majority of pathways have been experimentally verified to be active, an unknown number of the domains in this dataset might be inactive. To account for this, we employed a preliminary dead domain filter by eliminating domains with non-standard motif lengths (see Method for details). In this large dataset, we analyzed 83,489 C domains, 95,582 A domains, 86,688 T domains, 14,502 E domains, and 23,590 TE domains from bacteria; and 34,269 C domains, 40,458 A domains, 26,651 T domains, 3,982 E domains, and 4,008 TE domains from fungi.

The presence of NRPS pathways extends across multiple kingdoms, with fungi being the only known eukaryotic producers. Examining the similarities and differences in the NRPS sequences of bacteria and fungi sheds light on the conservation and divergence of NRPS functions. Unlike the MiBiG database, where most pathways are of bacterial origin, the larger dataset contains a comparable number of C, A, and T domains from both fungi and bacteria, providing a valuable platform for the validation and comparison of motif properties.

Overall, the motif logo and intermotif length of the A, T, E, and TE domains were found to be similar between bacteria and fungi, albeit with some subtle differences (Fig S7-S14). For example, the sequence logo of bacteria E6 was identified as RX(V/I/L)PXXGXG(Y/F)G while fungi E6 was RX(V/I/L)PXXGXXYF (Fig S11); the fungal intermotif A10-T1 was found to be 13 amino acids longer than that of bacteria by the median (Fig S13), and the fungal intermotif E2-E3 is 15 amino acid shorter than bacteria by the median (Fig S14), and. The disparity in the E6 motif highlights a potential bias in our current understanding of NRPS, which is predominantly informed by sequences collected from bacteria: Previous research reported that the E6 consensus sequence was PxxGxGYG, only consistent with the E6 sequence logo we observed in bacteria^59^. To expand our knowledge, a more comprehensive understanding in NRPS motif and intermotif across kingdoms is required.

#### 1.4. Identification of new C subtypes in fungi guided by conserved motifs

The C domain of NRPS manifested discrepancies between bacteria and fungi. It has been well established that the C domain can exist in multiple subtypes, exhibiting different conserved motifs^60, 64, 65^. To conduct systematic comparisons of NRPS between fungi and bacteria, it is crucial to clarify these subtypes first. Currently, antiSMASH v6 can distinguish LCL, DCL, Cglyc, Dual, Starter, Cyc, and X subtypes^34^, and NaPDoS v2 can distinguish LCL, DCL, Dual, Starter, Cyc, modAA, and Hybrid subtypes^66^. However, these tools are primarily based on bacterial subtypes and do not consider fungal subtypes such as CT and Iterative^24, 67^. This lack of annotation for fungal C domain subtypes hinders genome mining of fungal NRPS. In this study, we curated sequences representing 18 subtypes of the C and E domains, 14 of which were sourced from the conserved domain database (CDD)^68^, and the remaining 4 were obtained from literatures^32, 65, 69, 70^ (Table S1). Phylogenetic analysis of these subtypes revealed their relatedness (Fig 2A). These subtypes form two major clades: L-clade and D-clade, consistent with previous reports^64^. Furthermore, our results revealed for the first time the evolutionary landscape of all known C-domain subtypes, offering deeper insights in several sub-clades. It can be observed that, the CT subtype may evolve from the FUM14 subtype; the LCL subtype may evolve from SgcC5, and Cglyc may evolve from DCL. The FUM14 subtype was named from *Fusarium verticillioides* FUM14 (also known as NRPS8), a bi-domain protein with an ester-bond forming NRPS C-domain^71^. SgcC5 is a free-standing NRPS condensation enzyme (rather than a modular NRPS), which catalyzes the formation of both ester- and amide-bond^72^. Cglyc subtype is the glycopeptide condensation domain^60^.

**Figure 2.**
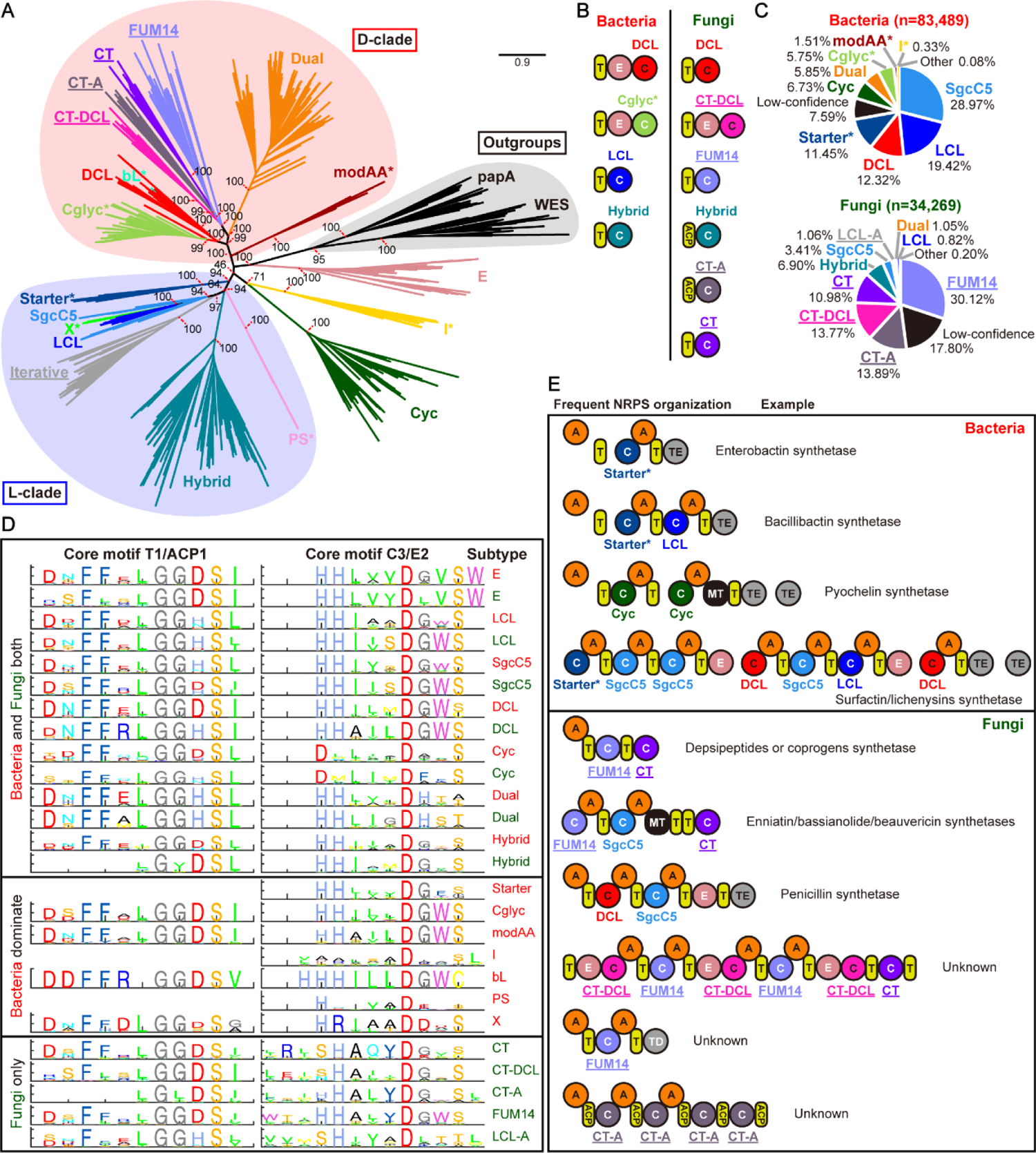
C domain subtype analysis and representative NRPS organizations in bacteria and fungi. **A.** Maximum-likelihood phylogenetic tree of the condensation domain superfamily. Subtype classification and sequences are described in the main text and the Method. Different subtypes are indicated by colors, with subtypes exclusive to fungi marked by underlines, and subtypes found predominantly in bacteria marked by asterisks. This is a rooted phylogenetic tree taking papA and WES as outgroups^64^, which are denoted by black shading. L-clade and D-clade are indicated by blue and red shading, respectively. **B.** Domains in adjacent with different C domain subtypes in bacteria and fungi. **C.** The statistics of subtype distribution in 83,489 bacterial C domains and 34,269 fungal C domains. C domains with HMM scores above the empirical threshold of 200 were annotated by their predictions, otherwise marked as “Low-confidence”. **D.** The sequence logo for the C3 or E2 motif from different C domain subtypes and the T1 or ACP1 motif adjacent to each subtype. Sequences from bacteria were marked by red, while sequences from fungi were marked by green. **E.** Frequent NRPS organizations with known representative examples in bacteria and fungi. Three NRPS organizations do not have known examples.

New insights on subclasses in C domains can be obtained by precisely characterizing this C domain phylogeny, particularly in fungal-related domains. For each C domain subtype, we constructed MSA from curated sequences by Muscle v5^73^ and built reference HMM models by hmmer v3^74^. We then utilized these HMM models to classify C domain subtypes, and extracted motifs for each subtype by previously mentioned method. During this process, we noticed that the sequence motifs of the curated CT subtype are remarkably variable, suggesting that this “CT” may not be a uniform subtype. Guided by the motif sequence and phylogenetic analysis, our analysis suggests that the original “CT” domain can be further classified into three subtypes by sequence similarity, which is also highly related to their locations in the NRPS pathway (Fig 2B): One of the subtypes is always located at the end of the NRPS, and we termed it “CT” because it fits the original definition of the CT subtype. Another subtype was termed “CT-DCL” because it is always behind an E domain and may have a similar function to the DCL in bacteria. Of note, we also observed that the annotated fungal DCL subtype is always located after the T domain. The third CT subtype is atypical because it is always behind an ACP (acyl carrier protein) domain rather than a T domain. Both the ACP and T domains are phosphopantetheinyl carrier domains, but the ACP domain has a conserved XGXDSL motif rather than the T domain’s GG(D/H)S(I/L) motif^59, 75^. Therefore, we termed this subtype “CT-A” (meaning atypical CT). These three CT subtypes form three different clades in the phylogenetic tree.

After the clarification of subtypes, the overall statistics regarding the distribution of C domain subtypes in bacteria and fungi can be analyzed. Our subtype prediction covered 92.41% C domains in bacteria and 82.20% C domains in fungi, by an empirical threshold of 200 score in HMM models (Fig 2C). Our predictions for the Starter and Dual subtypes are consistent with antiSMASH. However, antiSMASH frequently assign wrong annotations of “LCL” and “DCL” to subtypes that are uncommon in bacteria, which were corrected in our annotation (Table S2). Therefore, we redrew sequence logo of C domain in MiBiG by new subtype classification. The conservations of motifs increased significantly compared to the prior classification relying on MiBiG (Fig S1 and S15). In terms of subtype distribution, bacterial C domains mainly consist of SgcC5, LCL, DCL, Starter, Cyc, Dual, and Cglyc, while fungal C domains are mainly composed of FUM14, CT-A, CT-DCL, CT, and Hybrid. Among them, I (interface) only existed in bacterial C domain. Starter, Cglyc and modAA are almost exclusively found in bacteria, with a handful of occurrence in fungi (Table S3 and S4). FUM14, CT-A, CT-DCL, CT and an atypical LCL (designated as LCL-A, meaning atypical LCL which usually has a non-canonical conserved SHXXXDXX(T/S), rather than HHXXXDGXS) only exist in fungi.

These distinctions between fungal and bacterial C domains, as well as differences between subtypes, are readily apparent in the comparison of typical motif logos (Fig S16-S17), such as the C3 and T1 (Fig 2D). The C3 motif is essential for catalysis, and the T1 motif has been suggested to exhibit covariation with its preceding C domain^64^. Our findings highlight the variation in the location and motif of C domains in relation to the T domain between subtypes in fungi and bacteria (Fig 2D). Firstly, some C domain subtypes do not directly precede a T domain: the fungus Hybrid and CT-A subtypes are adjacent to an ACP domain, rather than a T domain; the bacteria DCL and fungus CT-DCL subtypes are located after an E domain, consistent with previous reports^64^; For the Cyc subtype, only 29.34% of them are adjacent to a T domain, while the rest are located at the start of the pathway. The Starter, I, and PS subtypes are also exclusively found at the start of the pathway. Secondly, we observed that the coupling between the C domain and its adjacent T domain cannot be solely explained by the L- and D-clades. Previous research based largely on bacterial data suggested that the LCL subtype (located in the L-clade of the phylogenetic tree in our analysis) is adjacent to a T domain with the LGGHSL motif, and the DCL subtype (located in the D-clade of the phylogenetic tree in our analysis) is adjacent to a T domain with the LGGDSI motif motif^63, 64^. Our analysis confirmed these relationships, but also showed that not all subtypes in the L- and D-clades follow this pattern: the X subtype in the L-clade is adjacent to a T domain with the LGGDSG motif; in the D-clade, and the Dual and fungal DCL subtypes in the D-clade are adjacent to T domains with the LGGHSI and LGGHSL motifs, respectively. These observations demonstrate that the T1 motifs are primarily linked to the specific subtypes of their adjacent C domains, rather than their clades.

In addition to the conserved motifs, the highly variable intermotif regions also differentiate fungi and bacteria based on some of their lengths. In the standardized motif-and-intermotif architecture, we compared the intermotif length distribution in different C domain subtypes between bacteria and fungi (Fig S18). The most notable difference is in the T1/ACP1-C1 region. C domain subtypes with more complex functions, such as Dual (92aa), modAA (74aa), bL (82aa), CT (86aa), CT-A(82aa), and FUM14 (85aa) have a substantially longer T1-C1 intermotif compared to C domains with simpler functions, such as LCL/ScgC5 (60aa in bacteria, 59aa in fungi). This finding implies that longer interdomain length may provide the necessary space for coordination between different domains in the megasynthetase. This distinction in C domain subtype compositions and intermotif lengths may offer potential applications in the future, allowing us to distinguish between bacterial and fungal NRPS in a piece of metagenome.

#### 1.5. The prevalent NRPS organizations in bacteria and fungi

With this detailed annotation of C domain subtypes, we can evaluate how domains are organized within NRPS pathways in bacteria and fungi. In this work, “organization” was defined as the composition and arrangement of domains within a NRPS pathway. For the accuracy of statistics, we focused on 12,364 bacterial and 6,438 fungal NRPS pathways that only contain the high-confidence C domains. Our findings show that in bacteria, 24 out of the 30 most frequent organizations are involved in the production of siderophores, accounting for 28.21% of the 12,364 pathways. Among these 24 siderophore pathway organizations, enterobactin^76^ (BGC0002476), the siderophore with the highest binding affinity to iron, has the highest occurrence (21.35% of 12,364). The second most frequent siderophore NRPS is bacillibactin synthetase^77^ (BGC0000309), making up 2.55% of the bacterial pathways, with an organization similar to enterobactin synthetase but with an additional module containing LCL subtype C domain. The third most frequent siderophore NRPS is pyochelin synthetase (1.16%, BGC0000412), containing two Cyc subtype C domains^78^. In additional to siderophores, biosurfactant surfactin/lichenysins synthetase are also frequent^79, 80^ (BGC0000433 and BGC0000381), representing 2 out of the 30 most frequent organizations and accounting for 1.24% of all pathways. This biosurfactant is comprised of there NRPS genes, with 1 Starter, 2 DCL, and 4 LCL/SgcC5 C domains. Interestingly, both pyochelin synthetase and surfactin/lichenysins synthetase contained 2 TE domains with the function of the second TE domain suggested to be for proofreading for the NRPS product^81^.

In fungi, the most frequent NRPS organization is also related to siderophore NRPS. Of the 20 most frequent NRPS organizations, 8 are depsipeptides or coprogens synthetase^67^, accounting for 11.90% of the 6,438 pathways. These synthetases typically consist of one NRPS gene with one FUM14 and one CT subtyped C domains, with some having an additional NRPS gene with an A domain (Fig 2E). These depsipeptides or coprogens synthetases are known for their iterative features, and their pathway organization suggests that the FUM14 subtype could also participate in the iterative process. However, unlike in bacteria, the products of many of the frequent NRPS organizations in fungi remain unknown (7 out of the 20 most frequent organizations, accounting for 7.72% of the pathways), highlighting the limitations in our current understandings of fungal NRPS. Other well-known fungal NRPS include enniatin/bassianolide/beauvericin synthetases^82–84^ (top 29-th in organizations, 0.36% of the pathways, BGC0000342, BGC0000312 and BGC0000313), which are also known for their iterative feature and have been the focus of reengineering for new products^85, 86^. These synthetases consist of one NRPS gene with one FUM14, one SgcC5, one CT C domain, again suggesting FUM14 subtype C domain may have an iterative function. Another known pathway (top 34-th, 0.33%) is the penicillin synthetase pathway^87^ (BGC0000404), which has a unique organization with its DCL located behind a T domain rather than an E domain, and its E domain positioned before a TE domain instead of a C domain. Almost all DCL in fungi have a similar organization to penicillin synthetase. Interestingly, in fungi, the function of DCL in bacteria seems to be replaced by CT-DCL. For example, the top 9-th most frequency fungal NRPS organization feature an E+CT-DCL arrangement (Fig 2E). In the entire database, such E+CT-DCL arrangements are quite common, with 66.14% (3121/4719) CT-DCL located behind an E domain, and 78.38% of E domain proceeding a CT-DCL (3121/3982). Another intriguing observation is that the top 12-th most frequent (0.87%) fungus NRPS organization ends with a TD domain, but the products of these pathways are unknown. Another unique organization among the fungus NRPSs is characterized by a prevalence of the CT-A subtype in their C domains, which can be found in 3.40% (219/6438) of the analyzed fungus NRPSs. While the CT-A subtype accounts for a substantial proportion (13.89%, 4759/34269) of the total fungal C domains, its presence in the analyzed 6438 fungus NRPS is limited, with only 618 instances observed. This suggests that most instances of the CT-A subtype may be associated with pathways of lower confidence C domains, which were excluded in our analysis of the pathway organizations. The results highlight the limitations of current annotations in large dataset, despite that the large dataset provided insights for confirming previously discovered information. Therefore, after analyzing this large dataset, we shift our focus back to the well-studied and annotated MiBiG database, where the well-established annotation offers a more suitable platform for in-depth discovery.

### 2. Discovery and functional identification of new conserved motifs

The presence of conserved motifs acts as anchor points in multiple sequence alignments, enabling an in-depth examination into the level of conservation across the entire sequence. From the MiBiG database, we calculated the amino acid frequency and gap frequency in sequences alignment of 1,161 C+A+T NRPS modules (1053 from bacteria, 81 from fungi, and 27 from “others”). The results are presented in Fig 3A, with a lower panel showing the conservation of the alignment. As expected, the majority of highly conserved positions are found within or close to known core motifs, thus can be seen as the extension of known motifs (Fig S19). However, there are three highly conserved regions that are relatively distant from well-known motifs (Fig 3A, upper panel), and hence cannot be considered extensions of these well-known motifs, suggesting potential new motifs.

**Figure 3.**
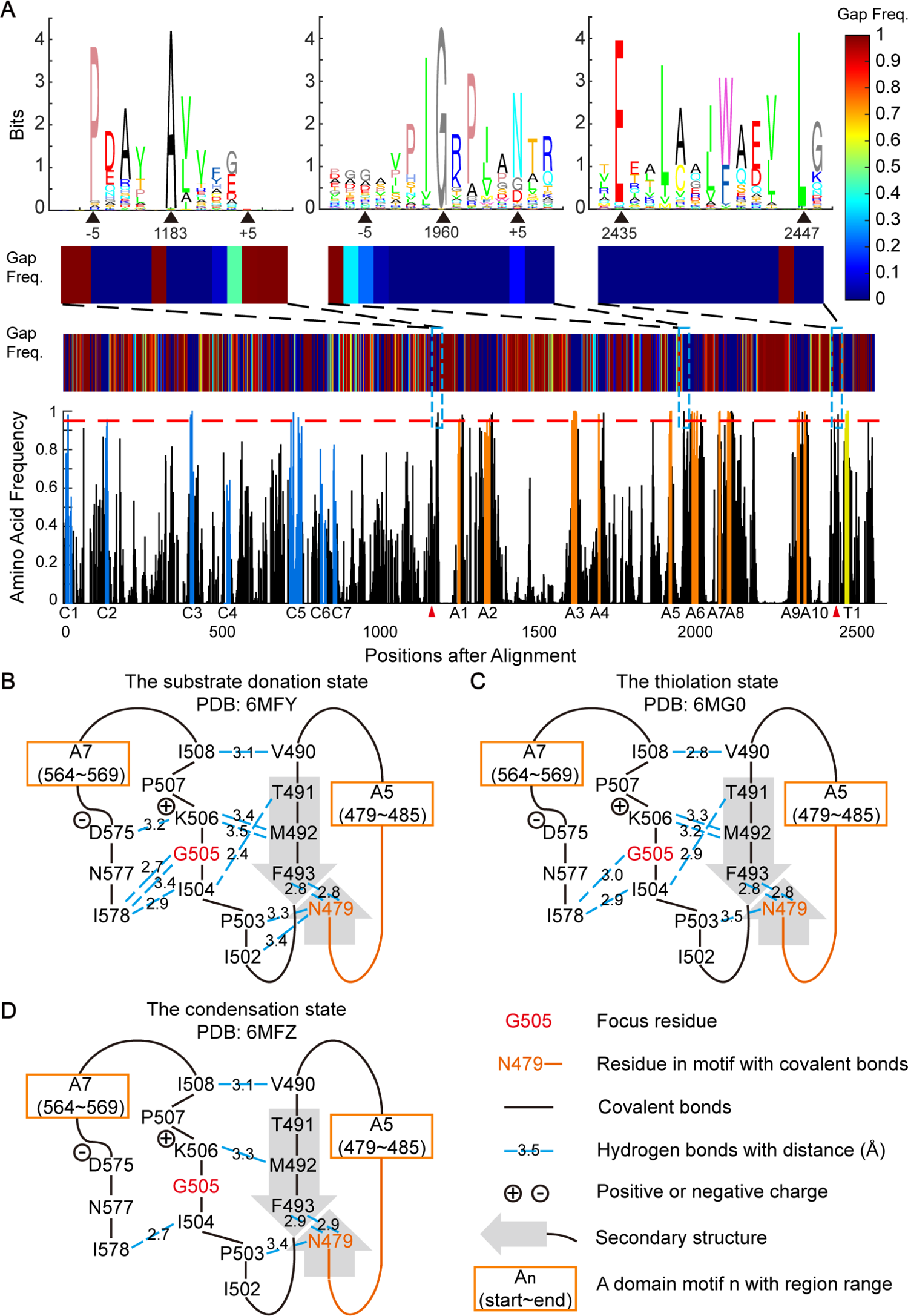
Analysis of amino acid frequency reveals potential new motifs with implications in structural flexibility. **A.** Amino acid frequency and gap frequency along the multiple sequence alignment of the NRPS CAT modules. In the bottom panel, bar heights indicate the frequency of the most frequent amino acid. Bars in the known core motifs from the C, A, T domains were colored blue, orange, and yellow, respectively. The horizontal red dashed line represents the 0.95 frequency level. Domain boundaries annotated by Pfam are divided by red triangles. The colored patch above amino acid frequency indicates gap frequency. Three potential new motifs (position 1183, 1960, and 2435/2447 in MSA) are marked by blue dashed box. The upper panel shows the sequence logo and the gap frequency near the three potential new motifs. **B.** Chemical interactions and secondary structures surrounding the second potential new motif in (A) in the substrate donation state (PDB: 6MFY). Hydrogen bonds near the most conserved Gly were shown in blue dashed-lines. Covalent bonds were shown as black lines. Secondary structures such as beta-sheets were shown as gray bold arrows. Known A domain motifs adjacent to related residues were shown in the orange box. **C.** Same as that in **B**, but in the thiolation state (PDB: 6MG0). **A. B.** Same as that in **B**, but in the condensation state (PDB: 6MFZ).

#### 2.1. The new “G-motif” in the substrate binding pocket relates to conformation flexibility

A conserved glycine located in the middle of the A5-A6 intermotif region has caught our attention. We also observed this conserved glycine in the A domain MSA in the large dataset (Fig S7 and S10). It is 21(±2) aa from the last residue of the A5 motif, and 23(±0.2) aa from the first residue of the A6 motif. This glycine is highly conserved in over 99% of the aligned A domains, with surrounding residues being moderately conserved (as shown in the second logo in the upper-most panel of Figure 3A). Notably, this glycine divides the A5-A6 intermotif region into two halves regarding substrate information, where the sequence preceding this glycine contains substantially more mutual information about substrate (Fig S20). For convenience, we will refer to this conserved glycine and its surrounding residues collectively as the “G-motif”.

We then located this G-motif to the G505 position in a recently solved NRPS structure, the linear gramicidin synthetase subunit A (LgrA)^62^. Interestingly, the G-motif resides on a loop structure, which is typically considered non-conserved. The LgrA structure has been solved with three function-related states: 6MFY, the substrate donation state; 6MG0, the thiolation state; and 6MFZ, the condensation state^62^. By calculating hydrogen bonds in these structures, we noticed that the G-motif (comprising residues I502, P503, I504, G505, K506, P507, and I508 in LgrA) interacts with N479 in the A5 motif (where the A5 spans N479-E485, with sequence NGYGPTE) and its backside residues (V490, T491, and M492). It also interacts with the backside residues of the A7 motif (D575 and I578; where the A7 spans Y564-R569 with sequence YRTGDR) (Fig 3B-3D). Of note, the number of hydrogen bonds for these interactions changes remarkably in different function-related states (Fig 3B-3D and Fig S21): In the substrate donation state 6MFY, the G-motif has 12 hydrogen bonds (average bond length 3.13±0.27Å); the thiolation state 6MG0 has 9 bonds (average length 3.09±0.24Å); and the condensation state 6MFZ has only 6 bonds (average length 3.13±0.27Å).

These changes observed in hydrogen bonds during confirmation changes prompted us to examine whether these interactions are functionally relevant. One prediction is that, in addition to the highly conserved glycine, the chemical properties of other residues involved in these interactions should also be conserved. Therefore, we turned back to the 1,161 MiBiG C+A+T alignment to check the chemical conservation of the residues interacting with the G-motif (Table S5). The cross-species conservation of chemical properties supported our prediction (Table 1). Of note, among these residues, only T491 and N479 use the hydroxyl or amide group in their side chains to form hydrogen bonds, while the other hydrogen bonds are formed by the commonly occurring α-carboxyl group and α-amino group in the main chains. The high degree of chemical conservation among these related residues suggests a strong selective pressure to maintain desired functions.

**Table 1.** Representative chemically conserved residues in or interacting with the G-motif. Conserved residues in or interacting with the G-motif have significant chemical properties. * This first letter is the prevalent amino acid in this position of MSA.

| The position of conserved residues |  | Amino acid composition | Properties of amino acid residues (side chains) |
| --- | --- | --- | --- |
| In the LgrA protein | In MiBiG C+A+T module MSA* | In MiBiG C+A+T module MSA |  |
| T491 | T1927 | T:61.67%<br>S:15.16% | Polar, -OH; the donor and acceptor of hydrogen bonds |
| I504 | I1959 | I:88.29%<br>V:5.25%<br>L:3.45% | Hydrophobic |
| G505 | G1960 | G:99.66% |  |
| K506 | R1961 | R:46.25%<br>K:23.84% | Positively charged; the donor of hydrogen bonds |
| I578 | L2094 | L:60.9%<br>I:29.03%<br>V:7.67% | Hydrophobic |

The small size and lack of a side chain of glycine make it unique among proteinogenic amino acids. We observed that two amino acids, N577 and F493 in LgrA, have a size near glycine in the G-motif (G505). One possible explanation for the high conservation of glycine in the G-motif is that its small size provides greater structural flexibility, reducing the likelihood of collisions with neighboring residues. To check this hypothesis in known structures of A domains, we analyzed all available structures of AMP-binding-domain-containing proteins from the PDB database (39 sequences existing in a total of 95 structures, including different conformations or ligands, Table S6). Of these 39 sequences, 30 are from NRPS pathways (selecting 20 different substrates), 6 from NRPS-like pathways (selecting 5 different substrates, with the substrates of the carboxylate reductase all being considered carboxylate), and 3 are from D-Ala-ligases (DltA) pathways (selecting substrate D-Ala). Notably, NRPS-like carboxylate reductases (CARs) do not contain the G-motif, showing a distinct evolution path despite also containing an AMP-binding domain (Table S6, “overview” sheet). Other than CARs, the G-motif is conserved in all 30 NRPS A domains, and 3 NRPS-like A domains (Fig S22). However, except for the NRPS-like protein poly-e-Lys synthetase (Pls-A, PDB: 7WEW), all AMP-binding-domain-containing proteins have equivalent N577 and F493 in LgrA, even for that don’t have a G-motif (Fig S23). The equivalent N577 and F493 have side chain sizes (Table S5). In these known structures, the G-motif is in close proximity to the adenylate part of the ligand (Fig S23), also suggesting a potential gatekeeper role of the G-motif. By simulated mutagenesis using PyMOL, we found that the position of glycine in the G-motif only permits small amino acids such as glycine and alanine. In simulation, mutating this residue into amino acids with larger volume or with inflexible loops resulted in a disturbance of the residue N397 and S491 in Figure 4A-B (equivalent F493 and N577 in LgrA, Fig 3B-3D).

**Figure 4.**
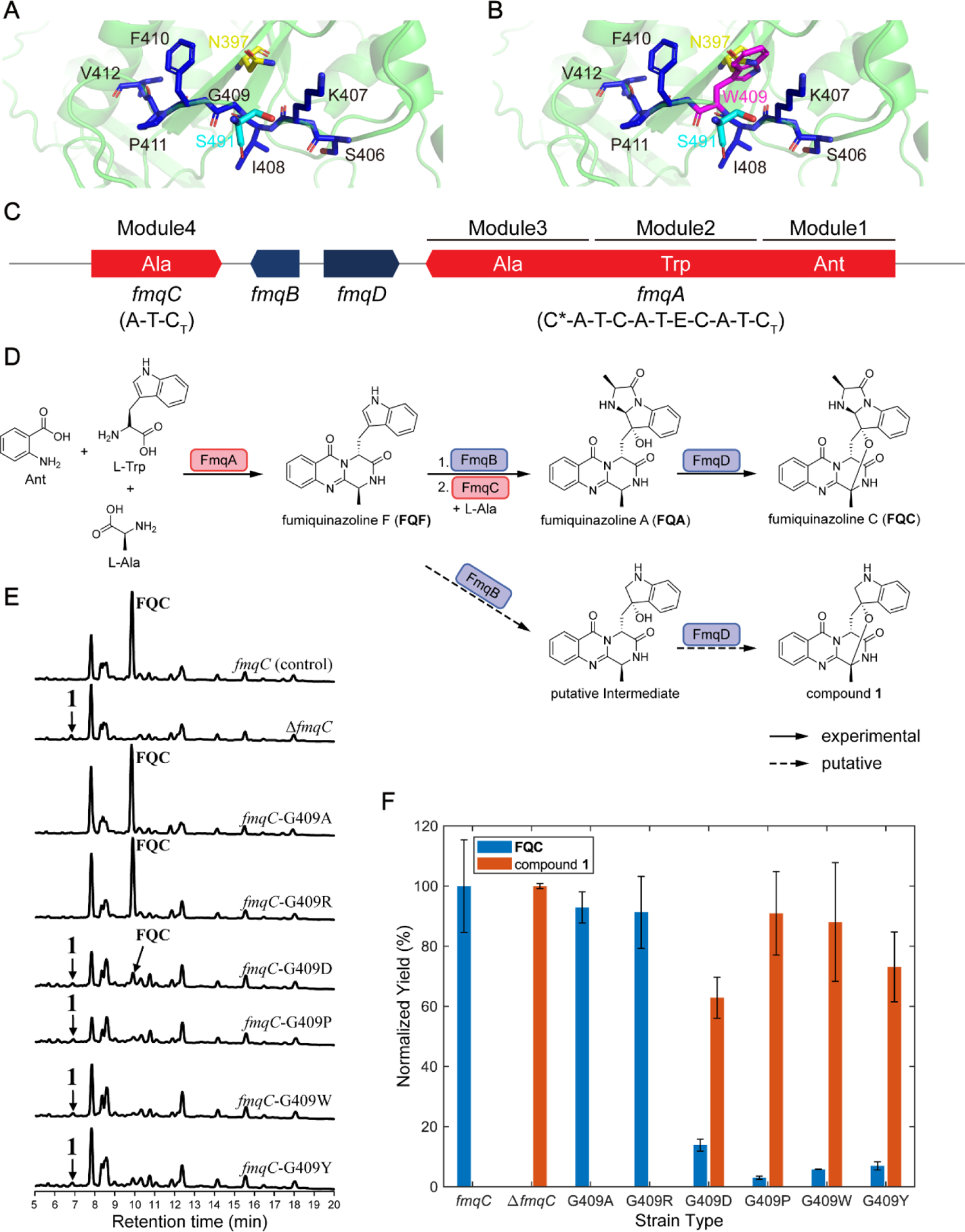
Mutations of G-motif G409 in FmqC support the importance of the conserved domain in the biosynthesis of fumiquinazoline C. **A.** Residues in and near the G-motif in the predicted structure of FmqC by Phyre2^91^. G409 is the conserved glycine in the G-motif. Residues in G-motif are marked by blue. The residues N397 and S491 (equivalent to F493 and N577 in LgrA, Fig 3B-3D) which may collide with G409 are marked by yellow and cyan. **B.** Same as that in A, but with simulated mutation of G409W. The mutated tryptophan is marked in magenta. **C.** The *fmq* gene cluster responsible for the production of FQC. Two NRPSs, *fmqA* and *fmqC*, are filled in red with their substrate selectivity marked. Ant: non-proteinogenic amino acid anthranilate. C* represents a truncated and presumably inactive C domain. C_T_ represents a terminal condensation-like domain which catalyzes macrocyclization reaction. **D.** The biosynthetic pathway for FQC is depicted, along with how it diverges into the production of compound **1** in the absence of functional FmqC. **E.** LC-MS analysis of the control *fmqC* (first row), Δ*fmqC* (second row) and six point mutation strains (3^rd^ to 8^th^ row). **F.** Normalized yield of FQC and compound **1** in different strains. For FQC, the yield is normalized by its production in the wild type strain. For compound **1**, the yield is normalized by its production in the *fmqC* gene deletion strain. Error bars show standard deviations.

In order to further verify the importance of this highly conserved glycine, we performed mutation experiments on the “G-motif” in a monomodular NRPS FmqC involved in the biosynthesis of fumiquinazoline C (FQC) in *Aspergillus fumigatus* (Fig 4C). In this system, a tripeptide precursor fumiquinazoline F (FQF) was first synthesized by FmqA and then the indole side chain of FQF oxidized by the FmqB. Then, the monomodular NRPS FmqC, activated L-Ala to create the product of fumiquinazoline A (FQA), thereby constructing a new C_2_–N bond and finally forming FQC^88^ (Fig 4D). The *fmqC* gene deletion resulted in the disappearance of FQC as the pathway diverged towards a putative compound (termed as compound **1**), as indicated in the first two rows of Figure S24. This is consistent with previous study in *A. fumtigatus* AF293^89^. We reintroduced *fmqC* into the original locus in the Δ*fmqC* background and generated one control strain with a full-length copy of *fmqC* in *A. fumtigatus*. The control strain could produce FQC just as the wild type (Fig S24). Then, the glycine in the G-motif of FmqC (G409) was mutated to six other representative amino acids (A, alanine, R, arginine, D, aspartic acid, P, proline, W, tryptophan and Y, tyrosine), to generate six FmqC (G409) mutants in *A. fumtigatus*, respectively. These mutations cover all combinations of structural effects estimated by Missense3D^90^ (Table S7). Following LC-MS analysis, the FQC yield of all mutants is lower than control. While the G-motif mutants G409A or G409R didn’t strongly reduce the yield of FQC, the other four mutants substantially reduced the yield of FQC (Fig 4E-4F). For example, when the glycine was mutated into proline (G409P), the yield of FQC was only around 3% of the control (Fig 4F and Table S8). Although we observed no accumulation of the precursor FQF in strains with substantially decreased FQC, we detected compound **1** as the divergent product in all of these strains instead (Fig 4D-4F). Our mutation experiment supported the critical role of the conserved glycine in the G-motif, and indicated that the flexibility of glycine is important in this position.

#### 2.2. Conserved motif at the start of the A domain

A conserved alanine located at the start of the Pfam annotated A domain was also identified through standardized multiple sequence alignment (Fig 3A, the first sequence logo in the first panel). This residue is positioned 11(±0.4) aa upstream of the A1 motif, and 124(±4) aa downstream of the C7 motif. This residue is found at A221 in the LgrA sequence. By analyzing hydrogen bonds based on the LgrA structures, we noticed that this conserved A221, V222 and another relatively conserved proline (P217 in LgrA) interact with residues in the A1 motif (L229, T230, Y231, and K232, where the A1 motif spans 229-234 in LgrA with sequence LTYKQL), as well as back side of residues from the A4 motif (L412 and I414; where the A4 motif spans 395-398 in LgrA with sequence FDGS). Direct interactions were also observed between A410, L406, and Y231 (Fig S25-S26). The chemical characteristics of residues anticipated to interact with the conserved alanine are also conserved in the alignment of 1,161 sequences, similar to the G-motif (Table S5). We termed this motif as “Aα1”, and it marks the start of the A domain. Previous studies have reported that Aα1 as a natural recombination site in Myxobacteria^92^ and applied it in synthetic biology for diverse NRPS products^93^.

#### 2.3. Conserved motif at the first helix of theT domain

The other two conserved residues outside of known motifs are glutamic acid and leucine, both of which are found at the start of the Pfam-annotated T domain (Fig 3A, the third sequence logo in the first panel). They were found at E700 and L711 in the LgrA sequence. The conserved glutamic acid is 10(±0.04) aa from the conserved leucine, which is 8(±0.6) aa before the first residue of the T motif. The T domain is known to be a distorted four-helix bundle with an extended loop between the first two helices^94^. According to the LgrA structure, the E700 and L711 are located at the N and C terminus of the first helix of the T domain, respectively. Distinct from the G-motif and the conserved A221, we found no hydrogen bonds outside the interior of this alpha helix. Instead, the E700 and L711 and other conserved aliphatic and aromatic amino acids take part in the formation of the hydrophobic core (Fig S27), agreeable with previous reports that the four helices were bound together by the hydrophobic interaction^95^. We termed this motif as “Tα1”, and it marks the start of the T domain.

### 3. SCA reveals overlapped co-evolving sectors across the C+A+T module

Coevolution analysis relies on proper sequence alignment, which can be facilitated by standardized NRPS structures. After examining the sequence properties of the standardized C+A+T module, we employed coevolution analysis to study the coupling between positions. Statistical Coupling Analysis (SCA), a commonly used technique for detecting “evolutionary units”, was applied to the 1,161 aligned and standardized C+A+T sequences from MiBiG. In the SCA result, only a few top modes (representing the collective covariations of a set of residues known as “sectors”) have eigenvalues (measures of the covariation magnitude) that are significantly larger than those obtained by random shuffling (26 out of 2560 total, Fig S28). The dominant first mode of SCA displays global correlations, which has been commonly regarded as phylogenetic association and generally ignored in subsequent analysis^54^. The remaining sectors, sorted in descending order based on eigenvalues, were referred to by their order and sign of the eigenvalues (see to Method for details). Sector II(+) and II(-) immediately after the phylogenetic mode contain 122 positions in the MSA with significant patterns of correlation (Fig 5A, two groups labeled green and magenta for II(+) and II(-), respectively). The II(+) sector and II(-) sector are orthogonal because they are the opposite directions of the same eigenvector. The II(+) sector primarily contains residues in the C domain, while the II(-) sector spans the entire NRPS module but does not contain any residues located in the pocket region of the A domain(Fig 5A).

**Figure 5.**
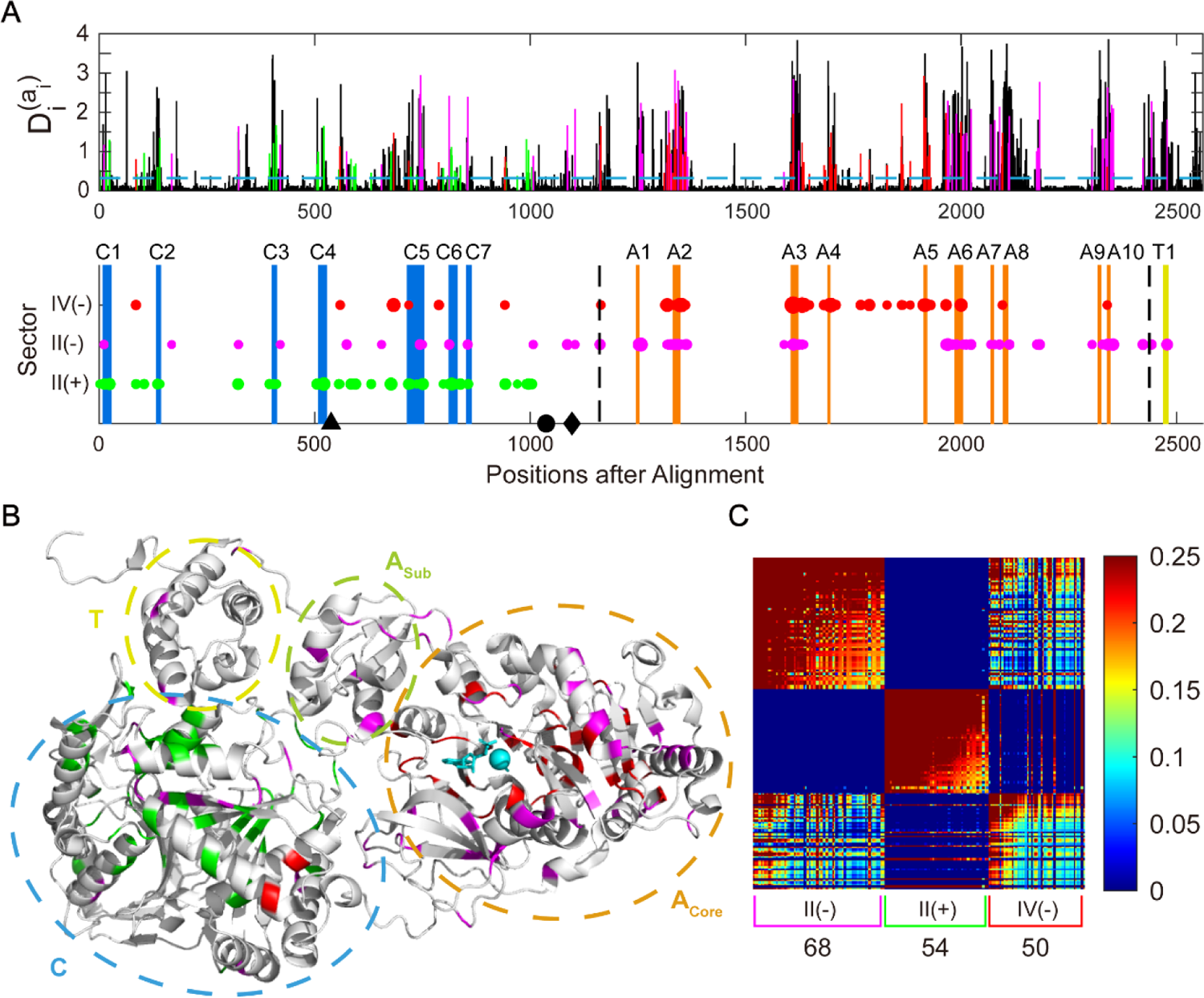
Statistical coupling analysis reveals overlapped sectors across the C+A+T module. **A.** The upper panel shows the conservation of residues in a multiple sequence alignment of 1,161 NRPS modules (containing the C, A, and T domains), quantified by the relative entropy D^(ai)^_i_ in SCA method. The mean conservation level (0.32) is marked by the blue dashed line. In the lower panel, there are three groups of positions (II(+) with green, II(-) with magenta, and IV (-) with red, termed “sectors”. Their corresponding conservations are marked in the same color in the upper panel. Blue bars mark C domain motifs from C1 to C7. Orange bars mark A domain motifs from A1 to A10. Yellow bar marks the T domain motif T1. Domain boundaries annotated by Pfam are divided by vertical black dashed lines. Black triangle marks reengineering point in the C domain reported by Bozhüyük et al.^26^, black circle marks reengineering point in the C-A inter-domain reported by Calcott et al.^27^ and black diamond marks reengineering point in the C-A inter-domain reported by Bozhüyük et al.^25^. **B.** Mapping three groups of correlated conservation positions into the three-dimensional structure of the NRPS module (PDB 4ZXI, containing the C, A, T, and TE domains. TE domain is hidden for clarity). Three sectors are marked in the same color as that in **(A).** C domain, A core domain, A sub domain, T domain are circled by blue, orange, yellow green, and yellow dotted line, respectively. Gly and AMP are substrates of this A domain. They and Mg2+ (for catalysis) are colored by cyan. **C.** Heatmap of the SCA matrix after reduction of statistical noise and of global coherent correlations (see Method for details). Each sector is marked by corresponding color bracket under the heatmap, with the number of contained residues listed. 68, 54, and 50 positions belong to II(-), II(+), and IV(-) sector, respectively. In each sector, residues are ordered by descending contributions, showing that sector positions comprise a hierarchy of correlation strengths.

Then we searched for sectors with a high proportion of residues from the A domain pocket region. The sector densely covering the A domain pocket region appears in the sixth biggest mode (Fig 5A, labeled as IV(-) sector with red dots), which includes three residues in the specificity-conferring code^28^. While the IV(-) sector varies nearly independently from the II(-) sector, residues contributing to the IV(-)and the II(-) sectors share a significant degree of covariation (Fig 5C), suggesting entanglements between substrate-specifying residues in the A domain and other regions of NRPSs. In detail, there are 5 positions (residue 85, 559, 718, 941, 942 in the MSA of Figure 5) shared between the II(+) and IV(-) sectors, and 6 positions (1164, 1351, 1358, 1605, 1631, 1635, 1965) shared between the II(-) and IV(-) sectors. In the II(+) sector, most positions are highly correlated. Therefore, overlapped positions with strong correlation were observed between the II(+) and IV(-) sector (Fig 5C). We noted that some residues of the C4-C5 and C5-C6 intermotifs are located in the II(+) sector and the IV(-) sector. The N-terminal of the C5-C6 intermotif is adjacent to the G-A6 intermotif region in the crystal structure, which is part of the A domain substrate binding pocket (A3-A6, Fig S29). Therefore, there is a possible coevolution in the overlapped region between the II(+) sector and the IV(-) sector.

Our coevolution analysis provides valuable insights for defining boundaries in NRPS reengineering. According to previous works on reengineering new proteins by recombination methods^22^, its recommended to avoid cutting into coevolution sectors when recombining sequences. However, our analysis indicated that there is no simple cutting point that can clearly separate all major sectors, as these sectors overlap with each other. Nevertheless, our findings are consistent with previously successful examples of reengineering NRPSs in their cutting point for recombination. As shown in the second panel of Figure 5A by black marks, Bozhüyük et al. cut the NRPS at the beginning of the IV(-) sector in their 2019 work of XUC (black triangle); Both Calcott et al. and Bozhüyük et al. in their 2018 work of XU cut NRPS after the ending of II(+) sector (black dot and diamond, respectively).

In addition, we also analyzed 685 C+A+T+C sequences and 245 C+A+T+E sequences by SCA (Fig S30A-S30B). In both sets of sequences, we found consistent sectors with the C+A+T analysis (Fig S31). In C+A+T+C, the first two sectors locate to two C domains, respectively, and the third sector spans across the four domains. In C+A+T+E, one sector locates to the C domain, and another sector spans across the four domains. Intriguingly, the third sector spans across the T and E domains, suggesting a possible cut point locating at the beginning of this sector, between the A and T domains (Fig S30).

### 4. Factors influencing the mapping from the A domain sequence to its substrate specificity

#### 4.1 Entanglement between substrate specificity and phylogenetic history

We then utilized a modified version of SCA to investigate the relationship between substrate specificity and residues in the A domain. Our analysis was based on 2,636 standardized A domain sequences with experimentally confirmed substrates, obtained from bacteria (2370 sequences), fungi (215), and other sources (51). These A domains were gathered from the supplemental material of SeMPI 2.0 research article^36^ (Table S9). These A domain sequences were aligned, and then their substrates were attached to the last column of the alignment (see Method for details). We primarily focused on modes that co-vary with the substrate column (Fig 6A). We found that residues contributing to substrate-related sectors are present throughout the A domain, with a higher concentration in the A3-A6 pocket region and a more scattered distribution in the A2-A3 region. Notably, the phylogenetic sector (Fig 6A, sector I) receives the second-largest contribution from the substrate column. It has long been recognized that phylogenetic relatedness results in consistent correlations across the whole sequence. As a result, substrate specificity, which is highly linked to the specificity-conferring code, is also linked to phylogenetic covariation throughout the A domain. This observation provides an explanation for the diversity of the specificity-conferring code even for the same substrate.

**Figure 6.**
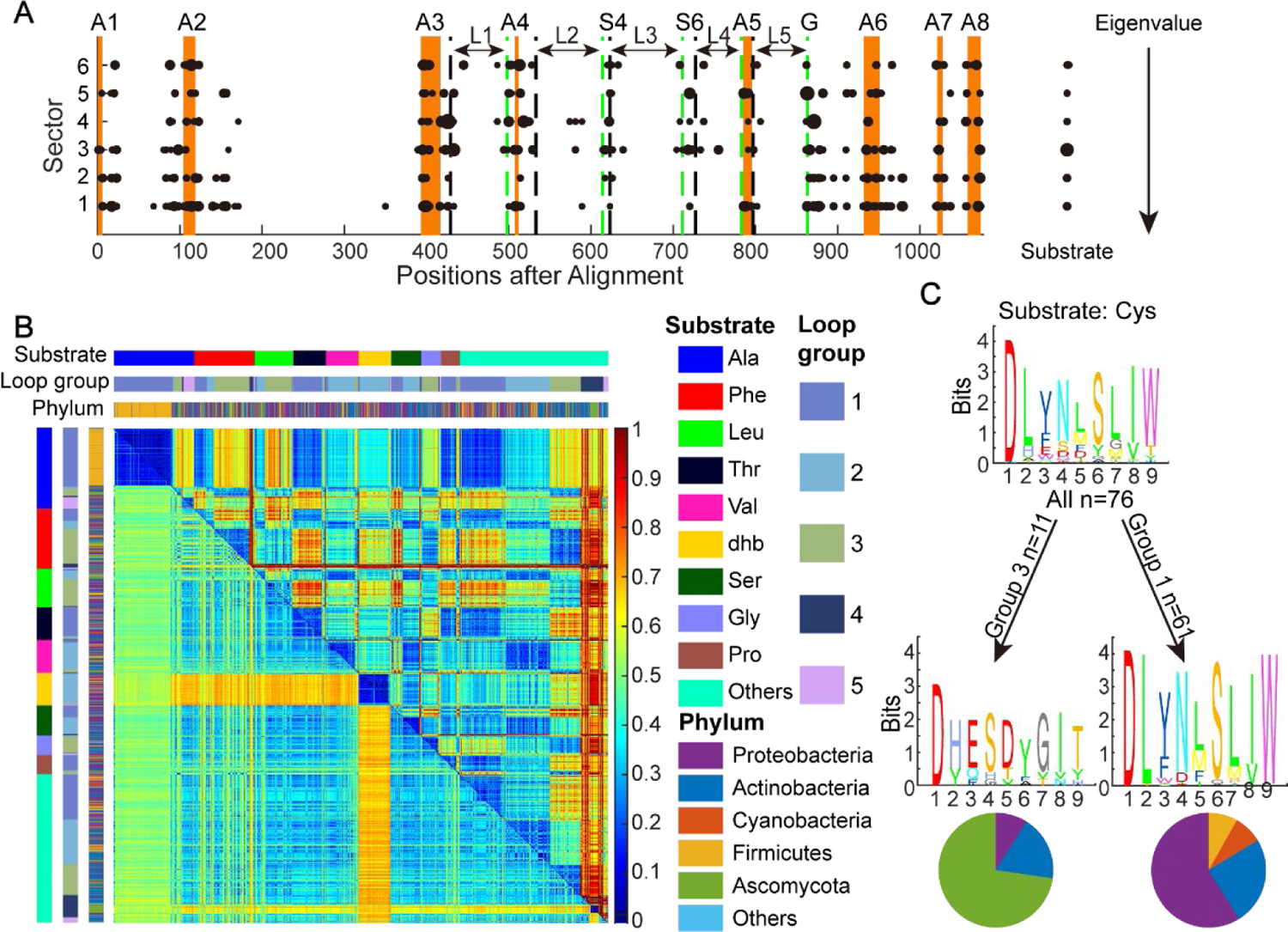
The specificity-conferring code of the A domain is correlated with loop length and phylogeny. **A.** SCA of 2,636 A domain sequences, together with their substrate specificities attached to the last column of the multiple sequence alignment. Six sectors with a high contribution from the substrate column (>0.05, the size of points on the left scales the substrate’s contribution to the sector, see Method for details) are sorted by their eigenvalues. The size of points scales its contribution to the sector. Orange bars mark the A domain motifs from A1 to A8. The start and end of the five loop regions are marked by black and green dotted lines, respectively. S4 and S6 are the 4^th^ and 6^th^ of the specificity-conferring codes. G is the G-motif. **B.** Distance matrix of A domain. Upper right on the heatmap is the Euclidean distance of the loop length as a 5-element vector. Lower left on the heatmap is the sequence distance of the A domain. The matrix is sorted by the substrate specificity followed by the loop length group. Substrates, groups of loop length, and phylum of these A domains, are shown by colors in side bars. **C.** Example showing that A domains conferring identical substrate exhibit distinct specificity-conferring codes, when they are categorized into different loop-length-groups. Phylum composition in each group is shown in the pie chart.

**Figure 7.**
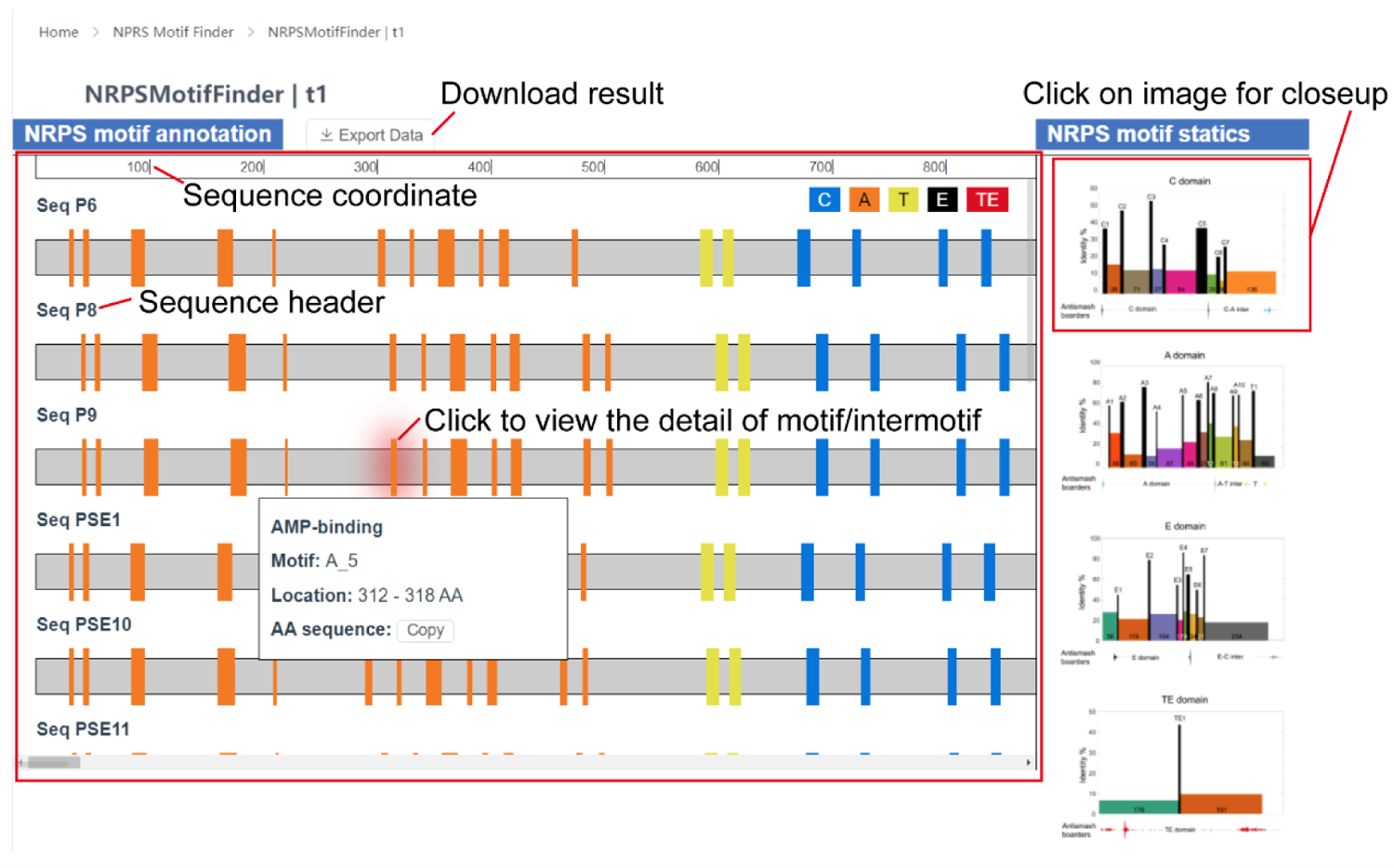
Demonstration on the result panel of the NRPS Motif Finder The NRPS Motif Finder result panel provides an interactive interface. The whole result could be navigated by scrolling the page, and details about motif and intermotif could be viewed by clicking the corresponding components. The general statistics about the NRPS architecture are displayed on the right for comparison. The results could be downloaded in table format. NRPS Motif Finder also supports the classification of 18 subtypes of the C domain, providing the first tool to annotate major fungal C domain subtypes. Furthermore, we provide all HMM files used in C domain subtype classification in supplemental materials for other researchers to use.

#### 4.2 Architecture of the substrate binding pocket relates to the diversity of the specificity-conferring code

Generally, SCA method gives higher weight to conserved positions^54^. Interestingly, some substrate-related sectors tend to locate in highly variable regions (Fig 6A and Fig S31). In aligning to PDB structures 1AMU and 4ZXI, we observed that these variable regions represent five protein loops in the A domain binding pocket (Fig 6A, loops region marked by arrows between motif A3-A6): A3-A4, A4-S4, S4-S6, S6-A5, and A5-G. A_n_ is the *n*-th core motif of the A domain. S_n_ is the *n*-th residue in the specificity-conferring code. G is the highly conserved glycine in the G-motif aforementioned).

Constructing an efficient MSA from highly varied regions is difficult. Therefore, to further understand the functions of these specificity-related loops, we compared their basic architectures, which are quantified by their length in amino acids. As the starts and ends of loops are highly conserved motifs and residues, loop lengths are relatively independent with MSA. These highly conserved sequence pieces help to anchor the alignment, so the length of the variable regions between motifs (i.e. the loop length) does not influence the alignment much. In variable regions, discrepancies in length correlate with larger sequence distances. Therefore, we estimated the Euclidean distances between pairs of A domains using the lengths of the five loops as a five-element vector. We observed that this loop-based distance is connected with, but not totally determined by, the similarity of sequence alignments and the phylogeny of A domains (Fig 6B).

This loop-based distance matrix was clustered hierarchically, generating five groups based on their loop-length vectors (Fig S32). Except for substrates Ala, Dhb, and Aad, one substrate typically corresponds to multiple loop-length groups. Importantly, the diversity of the specificity-conferring code is significantly reduced within each loop-length group. For example, for the substrate cysteine, the specificity-conferring code in the first loop-length group is readily distinguishable from that in the third loop-length group, despite the fact that these two groups contain A domains from distant phylogenies. Statistically, the diversity of the specificity-conferring code is reduced by identifying its loop-length vector group (Fig S33). To visualize the reduction, we decoupled the specificity-conferring code for specific substrates along the phylum and loop group (Fig S34-S41). In addition, we compared the loop length and loop group distribution between bacteria and fungi in MiBiG and the large dataset (Fig S42). We found a significant difference in loop group preference: the loop group 3 is dominant in fungi (85.86%) while it prefers the loop group 1 (37.89%), 2 (28.43%), and 3 (25.10%) in bacteria. In summary, we highlighted the importance of loop length for A domain substrate, which should be considered in the future A domain substrate prediction.

### 5. The NRPS Motif Finder online platform

Overall, such standardization enables statistical characterization of the sequence-function connections, supporting the use of known core motifs as “coordinates” of NPRS. To facilitate researchers in related fields, we constructed an online platform “NRPS Motif Finder” for parsing the motif-and-intermotif standardized architecture of NRPS from its coding sequences (http://www.bdainformatics.org/page?type=NRPSMotifFinder). It takes any query amino acid sequences in FASTA format and can be interacted with by clicking on the website result page for more information. In addition to the 26 well-established motifs, the NRPS Motif Finder also supports the three new motifs proposed in this article. The NRPS Motif Finder online version is based on Python, and the result could be downloaded in Excel format for further analysis. For researchers with basic programming skills, we recommend the NRPS Motif Finder Matlab version for timely updates (the source code is available in supplementary material).

## Discussion

About two decades ago, the core motifs of NRPSs were mapped out based on the early crystal structures and limited number of sequences available at the time^59^. Since then, modules, domains, and motifs were extensively utilized by annotation algorithms like antiSMASH^34^ and Pfam^37^ as well as experimental investigations^35, 36^, albeit with differing standards. Nowadays, with the rapid expansion of microbial sequencing data, systematic characterizations of NPRS are becoming possible. In this work, we presented a motif-and-intermotif architecture of NRPSs by partitioning the C, A, T, and TE domains by well-established core motifs. Guided by prior knowledge about these 19 motifs and domain characterizations, our standardized architecture presented a “common language” for processing NRPS sequences from various sources, enabling us to gain statistical insights from a large number of sequences without the need for manual curation.

Evolution imprints protein functions into their sequences. With the help of such a standardized architecture, novel insights acquired from sequence statistics exhibit amazing concordance with our point mutation experiments, as well as earlier findings derived from structure or reengineering efforts. For example, we identified novel conserved residues distant from known motifs, among which the G-motif seems to be the most intriguing. It centers on an exceptionally conserved glycine, where a single point mutation can abolish the enzymes function as we described in Figure 4. Interestingly, by using visual cues from protein structure, the G-motif has been used as the cut point of A subdomain for successfully reengineering A domains^96, 97^. Crüsemann et al. firstly proposed the A domain reengineering strategy by A subdomain swap guided by evolution in 2013^96^. Subsequently, Kries et al. proposed another A subdomain swap strategy in 2015, as they discovered that in the protein structure that the substrate binding region is a flavodoxin-like subdomain^97^. This subdomain starts from the middle of A3 and A4, and ends from the middle of A and A6 (the ending is exactly the G-motif, VPIGAPI in the PheA). Recently, Thong et al. also used this A subdomain swapping strategy by CRISPR-Cas9 and successfully re-engineered A domain substrate specificity by subdomains from a range of NRPS enzymes of diverse bacterial origins^98^.

NRPS undergoes substantial conformational changes as it exerts its multistep synthetase activity^62^. Identifying key residues associated with such a degree of structural flexibility is crucial for understanding NRPS functionality. In our analysis, three additional conserved motifs were identified. We hypothesized that the G-motif in the loop might be one of these key sites for structural flexibility, acting as a hinge when NRPS switches between distinct states. First, the number of hydrogen bonds around the conserved glycine changes substantially during the three function states. Also, glycine is the smallest of the 22 proteinogenic amino acids^99^, allowing the greatest structural flexibility. Additionally, it has been reported that the G-X-Y or Y-X-G oligopeptides motif provides the flexibility necessary for enzyme conformation change for catalysis, where X and Y are polar and non-polar residues, respectively^100^. G-R/K-P in the G-motif fit this G-X-Y structure. Furthermore, several enzymes have demonstrated the important function of conserved glycine residues in structure flexibility. For example, conserved G76 contributes to active-site loop flexibility in the pepsin^101^, and conserved G316 and G324 are the structural basis of efficient metal exchange in the cadmium carbonic anhydrase of marine diatoms^102^. In addition, the importance of this highly conserved glycine was verified in the fungal FQC biosynthesis system. We noted that the large amino acid in 409 position is mostly collided with N397 in simulated mutations (F493 in this position of MiBiG NRPS sequences, Fig 3B-3D). This may explain why mutating into large amino acids at the 409 position always reduce the FQC yield. Nevertheless, mutation into a small amino acid proline also strongly reduces the yield. It is possible that prolines side chain locks the dihedral angle Φ of the protein backbone at approximately −65° and causes significant conformational restriction^103^. Our results implied that the flexibility of glycine in the G-motif is essential for conformational change during A domain functions, therefore conformational restrictions in the G-motif impede A domain function.

We have also identified other conserved residues that may play a role in the functionality of NRPS. The conserved Ala and Pro at the start of the A domain (Aα1) may act as a bridge between the A1 and A4 motifs. Similarly, the conserved Glu acid and Leu in the first helix of the T domain (Tα1) may contribute to the proper folding of the T domain, much like the three conserved Gly residues contribute to the folding and function of green fluorescent protein^104^. From a functional perspective, these residues take part in formation of the hydrophobic core^94, 95^. In addition, 4’-phosphopantetheine cofactor required by T domain function is located in the vicinity of residues Y748, L751 and F752^105^ (numbered in LgrA. They are L65, L68 and F69 in the original research for holo-TycC3-PCP). Besides, Y748 and F752 also are the part of hydrophobic interface which interacts with C domain in docking process (in LgrA. V2534 and F2538 in original research for PCP2-C3 didomain from fuscachelin)^106^. In summary, the conserved residues we identified may contribute to the stability and flexibility of NRPS proteins in large conformational changes for product synthesis. In the future, it would be meaningful to experimentally explore these new motif candidates more thoroughly, to gain deeper insights into the conformational dynamics during NRPS functioning.

Another point of consistency between our sequence characterization and earlier reengineering studies is the SCA sectors that we identified. Among the three sectors we analyzed, the cutting point from Bozhüyük et al.’s 2019 work^26^ locates right at the beginning of the red sector, which contains residues enriched in the binding pocket of the A domain. The cutting points of Calcott et al. and Bozhüyük et al.’s 2018 work^25, 27^ both locate to the end of the green sector, which primarily represents covariation mode in the C domain. Congruence between sector boundaries and successful reengineering points substantiated SCAs potential to discern the “unit of evolution.”

Nonetheless, our analysis also dissected and reemphasized the complexities in reengineering NRPS. For example, sectors obtained from SCA intersperse with each other, even share covariation residues. Therefore, there is no universal cutting point in a NRPS module that does not disturb any of the major sectors. This observation is in line with difficulties in reengineering NRPS^22^. The success of Bozhüyük et al. and Calcott et al.^25–27^ may be partially attributed to the fact that their cutting points are right besides some of the major sectors, and they utilized sequences from NRPSs relatively close on phylogeny. It may be possible in the future to search phyla for NRPS systems with non-overlapping covariation sectors and use these systems as “building blocks” that can be recombined with greater freedom. As homologous recombination requires conserved sequence, the standardized architecture could aid in establishing the appropriate “cutting points” in such potential NRPS systems for reengineering.

The phylogenys entanglement with the substrate -specific binding pocket adds another layer of complexity in manipulating NPRSs. The contribution of the phylogeny sector to substrate specificity suggests non-degeneracy of the sequence space in determining substrate: the same substrate can be selected by distinctive binding-pocket sequences, which drift with phylogenies and may recombine with each other. Such non-degeneracy leads to the diversity of the specificity-conferring code, causing troubles in the targeted design of the A domain specificity. We demonstrated that knowing the length of the five loops in the pocket region reduces the specificity-conferring code diversity, and tried to provide an explanation by a causal diagram (Fig S43, the causal diagram is a method for causal inference^107^). It suggests there is a connection between the specificity-conferring code and loop length group for a specific substrate (Fig S34-41). However, a mechanistic understanding of substrate selectivity is still needed to guide the rational design of the A domain. The architectural distinctions and loop length group preference between fungal and bacterial A domains discussed may shed light on the detailed mechanism of substrate selection, and help to establish the framework for developing substrate prediction algorithms that are less reliant on experimentally validated substrates.

The standardized motif-intermotif architecture not only enables efficient analysis of large datasets, but also provides a useful framework for in-depth examination of individual NRPS pathways. To facilitate researchers in related fields, we presented the NRPS Motif Finder online platform with a user-friendly tool to construct the motif-and-intermotif architecture of NRPS. The resulting architecture can be used in A domain substrate prediction based on phylogenies of A3-A6 or A4-A5. Also, this tool can be used in guiding reengineering and new NRPS discovery, as G-motif and Aα1 both coincide with known cutting points^92, 93, 97^. In addition, NRPS Motif Finder supports the classification of C domain subtypes into 18 kinds, offering the first tool to cover most of the fungal C domain subtypes.

In terms of limitations, most of our results are based on computation. Despite the high degree of concordance between computational predictions and our point mutation experiments on the G-motif, more comprehensive experiments might be developed in the future to explore the roles of other conserved residues we identified. Additionally, given the diversity of NRPSs, despite our use of the largest A domain database with known substrates, A domains selecting rare substrates may be unrepresented. Moreover, the identification of the new C domain subtypes, while promoting the understanding of fungal NRPS, also highlighted the current limitation of the dataset, which is heavily biased towards bacterial BGCs. The catalytic mechanism and evolutionary history of new C domain subtypes are still unclear and require more investigation. Overall, our effort is a preliminary investigation into the possibilities of standardized architecture in modular enzymes. Much more discoveries could be achieved in the future with the rapid expansion of microbial sequencing data.

## Method

### Data acquisition

We downloaded the MiBiG database version 2.0^32^, then extracted sequence information from this database. In this research, only C/A/T/E/TE domains from NRPS clusters were considered. Finally, 326 NPRS sequences from 264 species were obtained, with 1,864 A domains, 1,765 C domains, 1,803 T domains, 310 E domains, 280 TE domains. In total, there are 1,161 C+A+T modules, 685 C+A+T+C modules, and 245 C+A+T+E modules.

A domains sequences with known substrates were gathered from the supplemental material of SeMPI 2.0 research article^36^. Some of the A domains from SeMPI overlap with those in MiBiG (http://sempi.pharmazie.uni-freiburg.de/database/MiBiG). Sequences which had deleted in MiBiG database or UniProt^108^ were removed from our dataset. Finally, we collected 2,636 A domain sequences. Sequence details see Table S9. Pfam seed alignments were downloaded for comparison with antiSMASH domains (PF00668 for C domain, PF00501 for A domain, PF00550 for T domain).

Sequences with high identity were removed (Fig S44). All analysis was applied to the remaining sequences. For the large dataset to confirm our results, we downloaded all complete bacterial genomes (30,984) and all assembly-level (including “contig”, “scaffold”, “chromosome”, and “complete” levels) fungal genomes (3,672) from NCBI (as of 2022/10/23). Genomes were deduplicated by Mash distance with a cutoff of 0.004 (>=99.6% genome similarity) using Mash tool v2.3^109^. Representative genomes were chosen by picking the genome with the longest genome size. After removing redundancy, we had 16,820 bacterial and 2,505 fungal genomes. The information of used genomes was summarized in Table S10. Species information was obtained by TaxonKit^110^.

### Length threshold for multi-domain NRPS form MiBiG

Considering the NRPS sequence length distribution, we filtered sequences by the following thresholds. For C+A+T, the sequence length should be more than 1000 and less than 1300; for C+A+T+C, the sequence length should be more than 1500 and less than 1750; for C+A+T+E, the sequence length should be more than 1350 and less than 2350. In practice, 14 sequences in the C+A+T+C type are removed because their secondary C domain, rather than the first C domain, align with the first C domain of other sequences. And a few sequences from C+A+T+E were removed because they are incomplete at start.

### Multiple sequence alignment

Most of the MSA in our work were constructed by Cluster Omega 1.2.4^111^. For the construction of C domain subtype reference HMM files, we prepared MSA by Muscle v5 refer to previous studies^60, 73^.

### antiSMASH annotation

Data obtained from MiBiG database has already been annotated by antiSMASH 5.1^32^. BGCs in the large dataset were annotated by antiSMASH 6^34^.

### Conserved motif detection for NRPS domain

As shown by Figure S45, first, manual curation of the conserved motifs from literatures was performed for all domains. That resulted in 7 motifs for C, 10 motifs for A, 1 motif for T, 7 motifs for E, and 1 motif for TE^59^. A table of these conserved motifs according to previous literatures were summarized in Table S11. Second, we obtained reference sequences for each type of NRPS domain. The reference sequences of the A, T, and TE domains were from surfactin A synthetase C which has resolved crystal structure (PDB: 2VSQ)^57^. The reference sequence of the E domain was retrieved from the initiation module of tyrocidine synthetase A (PDB: 2XHG)^58^. The reference sequences of C domain are described in the section of C domain subtypes classification. The motif locations on all reference sequences above were manually identified, according to the results from step 1. Third, for every query sequence, global alignment was performed against possible reference sequences. Regions on the query sequence that align with the conserved motifs in the reference sequence were recorded as “motifs”. For clarification, only sequences with the prevalent motif length were used in the construction of motif sequence logos. The prevalent motif lengths of each domain were recorded in Table S11.

### C domain subtypes classification

First, we curated sequences from the E domain and 18 subtypes of the C domains as reference. 14 of 18 subtypes of the C domains were sourced from the CDD^68^, and the remaining 4 were obtained from literatures^32, 65, 69, 70^ (Table S1). The original “CT” subtype sequences were classified into CT, CT-DCL and CT-A subtypes guided by our phylogenetic anlaysis. We prepared MSA for each subtype by Muscle v5^73^ and constructed reference HMM files by hmmer v3.1b2^74^. The MSA results of each C domain subtype were attached in the supplementary materials.

Finally, we aligned the query sequence to profile using hidden Markov model alignment. In the NRPS Motif Finder online version (python), it is achieved by the hmmscan function in hmmer3. In the NRPS Motif Finder Matlab version, it is achieved by the hmmprofalign function in Matlab.

### Calculation of mutual information

In generating the fourth panel of Figure 1A, mutual information between the residues and the chirality subtypes of C domains was calculated. The C domain subtype was defined by the antiSMASH annotation, including “LCL”, “DCL”, “Dual”, “Starter”, and “Heterocyclization”. In the multi-alignment of C domains, the reduced Shannon entropy on the amino acid composition at position *i* given the subtype, was calculated as the mutual information about chirality at position *i.* The fourth panel of Figure 1B was obtained by calculating the mutual information between substrate specificity and amino acid composition at each position of the multi-alignment of A domains.

### The position of motif boundaries for multi-sequence

The motif boundaries in Figure 3 and 5-6 and Figure S30-S31 show the average motif boundaries positions from each single sequence. Motif boundaries positions are consistent in different sequence after MSA other than the C domain, where the conserved motifs vary by subtypes (Table S11).

### The calculation of conservation in NRPS protein

For a quantitative perspective, we use ConSurf^112^ and AACon^113^ to calculate the conservation of specific positions in the NRPS protein. The results of two methods are similar in very conserved positions, but a little different in middle conserved positions. The results are displayed in Table S5.

### Statistical coupling analysis

The statistical coupling analysis was performed with the binary approximation method^54^. The cleaned correlation matrix is obtained using the method described in the same article^54^. To investigate substrate-related positions, we attached substrates as the last column of the multiple sequence alignment. Then we applied SCA to this modified MSA. The least frequent substrate was taken as background of this position.

### Sector identification in SCA

We use the same notation in Halabi et al.^54^. Except two sectors of the second mode with opposite eigenvalues, third sector was chosen empirically. The bra-ket notation is such that |k⟩ denotes the *k^th^* eigenvector and 〈i|k〉 the weight for position *i* along eigenvector *k*. The threshold ε to separate significant weights along an eigenvector from statistical noise is 0.05, which was used in Halabi et al.^54^.

For C+A+T, green sector is defined as 〈*i*|2〉 > ε; magenta sector is defined as 〈*i*|2〉 < *ε*; red sector is defined as 〈*i*|6〉 < *ε*.

For C+A+T+C, green sector is defined as 〈*i*|2〉 > *ε*; magenta sector is defined as 〈*i*|4〉 > *ε*; red sector is defined as 〈*i*|2〉 < *ε*.

For C+A+T+E, green sector is defined as 〈*i*|2〉 < *ε*; magenta sector is defined as 〈*i*|3〉 > *ε*; red sector is defined as 〈*i*|2〉 > *ε*.

For A domain, from bottom to top, sectors are defined as 〈*i*|1〉 > *ε*, 〈*i*|2〉 > *ε*, 〈*i*|3〉 > *ε*, 〈*i*|4〉 > *ε*, 〈*i*|5〉 > *ε* and 〈*i*|6〉 < *ε*.

### Loop length clustering

The sum of the Euclidean distance of five loops between different A domain sequences was calculated for distance matrix. To avoid the influence of local maximum value on the heatmap, the upper-bound in distance was set to 12. Then hierarchical clustering analysis is applied to this distance matrix (Fig S32).

### Loop length profile calculation method

Related to Figure S32, 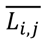 is normalized length of loop i in j^th^ row. *L_i,j_* is actual length of loop i in j^th^ row. *L_i_* is a column vector containing all length of loop i.

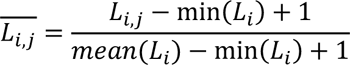

### Experimental Materials and General Methods

The plasmids and strains utilized in this work are listed in Table S12. The oligonucleotide primers synthesized by Shanghai Sango Biotech are given in Table S13. PCR reactions were carried out with FastPfu high-fidelity DNA polymerase (Transgene Biotech). Escherichia coli strain DH5α was propagated in LB medium with appropriate antibiotics for plasmid DNA, and plasmid DNA was prepared using the Plasmid Mini Kit I (Omega). Automated DNA sequencing was performed by Shanghai Sango Biotech. PCR screening for transformants were carried out using 2×GS Taq pcr mix (Genesand). LC-MS analysis was performed on an Agilent HPLC 1200 series system equipped with a single quadrupole mass selective detector, an electrospray ionization (ESI) and an Agilent 1100LC MSD model G1946D mass spectrometer by using a Venusil XBP C18 column (3.0 x 50 mm, 3 mm, Bonna-Agela Technologies, China) and equipped with an electrospray ionization (ESI) source.

### Gene cloning, plasmid construction, and genetic manipulations

#### (a) Creation of *fmqC* deletion strains in *A. fumigatus*

For site-directed mutagenesis of G-motif in *A. fumigatus*, *fmqC* was fully deleted in Cea17.2 to increase the rate of homologous recombination and reduce the difficulty of screening transformants. The FmqC (accession: EDP49773.1) deletion strain was created in Cea17.2 by replacing the *fmqC* with hygromycin phosphotransferase gene (*hph*) using modified double joint PCR described previously^114^ consisting of the following: 1 kb DNA fragment upstream of *fmqC*, a 1.9 kb DNA fragment of *hph*, and a 1 kb DNA fragment downstream of *fmqC*. Third round PCR product were purified for protoplast transformation (fragment concentration greater than 300ng/µL). Polyethylene glycol (PEG) based transformation of *A. fumigatus* was done as previously described^115^. The fragment concentration was 3 µg per 100 µL of protoplasts. The mutants were confirmed by using diagnostic PCR and the correct transformants were used for subsequent analysis.

#### (b) Creation of *fmqC* mutation strain in *A. fumigatus*

Seven plasmids were generated, one which included a full-length copy of *fmqC* (pYJY25), and the others were mutated *fmqC* (pYJY26-31). The pYJY25 plasmid was assembled by amplifying 1 kb DNA fragment upstream of *fmqC*, *fmqC*, *A. fumigatus pyrG* as the selectable marker, and a 1 kb DNA fragment downstream of *fmqC*. The fragments were assembled into a full plasmid by homologous recombination using MultiS One Step Cloning Kit (Vazyme). pYJY26-31 were assembled using the same fragments and method, except that the targeted mutations of 409Gly in G-motif was introduced by using primers containing the mutation and amplifying *fmqC* in two separate PCR reactions. After plasmid construction, the fused 8kb PCR fragments with the point mutation were purified and used for transformation to TYJY81. Transformants were confirmed by sequencings to obtain TYJY82-88 for the subsequent experiments.

### Detection of secondary metabolites by LC-MS analysis

*A. fumigatus* strains were cultivated in glucose minimal medium (GMM)^116^ culture medium at 28 °C for 3 days in the dark. The media and mycelia were extracted with ethyl acetate, and then evaporated under reduced pressure. The extract was dissolved in 1 mL methanol, and 5 μL of the solution was directly injected for LC-MS analysis. Water with 0.1% (v/v) formic acid and acetonitrile were used as mobile phase with a flow rate of 0.5 mL/min. For analysis of the extracts, a linear gradient of acetonitrile in water (10-45%, v/v) in 30 min was used and washed with 100% (v/v) acetonitrile for 5 min.

## Data Availability

NRPS Motif Finder is an online platform for construction of NRPS motif-and-intermotif architecture (http://www.bdainformatics.org/page?type=NRPSMotifFinder). The source codes are available in supplementary material.

## Supporting information

C domain subtype reference HMM files

C domain subtype reference sequence MSA

NRPS Motif Finder Matlab version code

NRPS Motif Finder online version code (Python)

The result file of the phylogenetic tree of C domain and E domain by IQ-TREE

## Acknowledgements

This work was supported by grants from Peking-Tsinghua Center for Life Sciences. We thank Haonan Zheng for helpful discussions. We also appreciated the tech team at the BDA Informatics Suite (www.bdainformatics.org) for the online platform.

## Author contributions

Ruolin He collected the data, performed the majority of computational analysis in this research, drafted the manuscript, and suggested mutation sites for the experiments. Jinyu Zhang was responsible for execution and analysis of the mutation experiments. Yuanzhe Shao wrote python scripts for online platform. Shaohua Gu offered insightful comments and assisted in revising the manuscript. Chen Song offered insightful comments in structural analysis of the G motif. Long Qian assisted in constructing the NRPS Motif Finder platform. Wen-Bing Yin oversaw the project, designed experiments and revised the manuscript. Zhiyuan Li conceptualized and oversaw the project, conducted the initial analysis of the NRPS architecture, and revised the manuscript.

## The legend of Supplementary Materials

**Table S1.** Source of C domain subtype reference sequences

**Table S2.** Subtype prediction by NRPS Motif Finder and annotation by antiSMASH v5 for C domains in MiBiG

**Table S3.** Subtype prediction by NRPS Motif Finder and annotation by antiSMASH v6 for C domains in 16,820 bacterial genomes

**Table S4.** Subtype prediction by NRPS Motif Finder and annotation by antiSMASH v6 for C domains in 2,505 fungal genomes

**Table S5.** Amino acid composition and conservation in three potential motifs

**Table S6.** All available structures of AMP-binding-domain-containing proteins from the PDB database

**Table S7.** Structural effects of point mutations in G-motif G409 of FmqC estimated by Missense3D

**Table S8.** Product yields in different strains by HPLC/MS analysis

**Table S9.** 2,636 A domains sequences information

**Table S10.** Bacterial and fungus genome information used in this study

**Table S11.** The definition of conserved motifs in NRPS domains and the positions in reference sequences.

**Table S12.** Fungal plasmids and strains used in this study

**Table S13.** PCR primers used in this study

## Figure

**Figure S1.** Sequence logo of the seven C motifs among the multialignment of 1758 C domains (first row), and among six subtypes of C domains (second row) in MiBiG

**Figure S2.** Sequence logo of the ten A motifs among the multialignment of 1859 A domains in MiBiG

**Figure S3.** Sequence logo of T1 motif and the length distribution of the T1-C1 region in MiBiG

**Figure S4.** Sequence logo of the seven E motifs among the multialignment of 310 E domains in MiBiG

**Figure S5.** Sequence logo of the TE1 motifs among the multialignment of 280 TE domains in MiBiG

**Figure S6.** Length distributions of C, A, and T domains, in MIBIG and in Pfam seeds

**Figure S7.** Sequence logo of the twelve A motifs and two T motifs among 95,582 A domains and 86,688 T domains in bacteria

**Figure S8.** Sequence logo of the seven E motifs among 14,502 E domains in bacteria

**Figure S9.** Sequence logo of the TE1 motifs among 23,590 TE domains in bacteria

**Figure S10.** Sequence logo of the twelve A motifs and two T motifs among 40,458 A domains and 26,651 T domains in fungi

**Figure S11.** Sequence logo of the seven E motifs among 3,982 E domains in fungi

**Figure S12.** Sequence logo of the TE1 motifs among 4,008 TE domains in fungi

**Figure S13.** Comparison of NRPS A and T domain architecture between bacteria and fungi

**Figure S14.** Comparison of NRPS E domain architecture between bacteria and fungi

**Figure S15.** Sequence logo of the seven C motifs among the multialignment of 2,572 C domains (first row), and among 15 subtypes of C domains (second row) in MiBiG

**Figure S16.** Sequence logo of the seven C motifs among 77,152 C domains (first row), and among 13 subtypes of C domains (second row) in bacteria

**Figure S17.** Sequence logo of the seven C motifs among 34,269 C domains (first row), and among 11 subtypes of C domains (second row) in fungi

**Figure S18.** Comparison of NRPS C domain architecture between bacteria and fungi

**Figure S19.** Sequence logo of highly conserved positions near known motifs

**Figure S20.** The mutual information between residues in the A5-A6 and A domain substrate specificity

**Figure S21.** The G-motif in different function states of LgrA structure

**Figure S22.** The sequence logo of G-motif in A domains with known structures and FmqC

**Figure 23.** The equivalents of N397, G409 (G-motif), and S491 in FmqC mapped to known structures

**Figure S24.** LC-MS analysis in the strain construction

**Figure S25.** Aα1 motif in different states of LgrA structure

**Figure S26.** The interaction near Aα1 motif in different states of LgrA structure

**Figure S27.** Conserved resides of T domain in the condensation state

**Figure S28.** Eigenvalue spectra for the SCA matrix of C+A+T modules MSA and random MSA

**Figure S29.** The interaction of C5-C6 with other domains in LgrA structure

**Figure S30.** SCA analysis for C+A+T+C and C+A+T+E four domains NRPS sequences

**Figure S31.** The gap frequency in the MSA of 2,636 A domains

**Figure S32.** Clustering and groups of five loops

**Figure S33.** The entropy and conditional entropy of the specificity-conferring code given different constraints.

**Figure S34.** The sequence logo of the specificity-conferring code for substrate alanine in the dimension of phylum and loop group

**Figure S35.** The sequence logo of the specificity-conferring code for substrate phenylalanine in the dimension of phylum and loop group

**Figure S36.** The sequence logo of the specificity-conferring code for substrate leucine in the dimension of phylum and loop group

**Figure S37.** The sequence logo of the specificity-conferring code for substrate valine in the dimension of phylum and loop group

**Figure S38.** The sequence logo of the specificity-conferring code for substrate tyrosine in the dimension of phylum and loop group

**Figure S39.** The sequence logo of the specificity-conferring code for substrate 2-amino-adipic-acidin the dimension of phylum and loop group

**Figure S40.** The sequence logo of the specificity-conferring code for substrate glutamine in the dimension of phylum and loop group

**Figure S41.** The sequence logo of the specificity-conferring code for substrate diaminobutyric acid in the dimension of phylum and loop group

**Figure S42.** Loop length and group distributions in bacteria and fungi

**Figure S43.** Causal analysis of A domain substrate specificity

**Figure S44.** Protein sequence pairwise distance distribution

**Figure S45.** Workflow of detecting conserved motifs in NRPS domain

## Other supplementary materials

1. C domain subtype reference HMM files
2. C domain subtype reference sequences (MSA)
3. The result file of the phylogenetic tree of C domain and E domain by IQ-TREE
4. NRPS Motif Finder online version code (Python)
5. NRPS Motif Finder Matlab version code

