## Supplemental Table 8, Supplemental Table 12, Supplemental Table 13, Supplemental Figure 1-45 for "Knowledge-guided data mining on the standardized architecture of NRPS: subtypes, novel motifs, and sequence entanglements"

This file contains **Table S8**, **Table S12-S13** and **Figure S1-S45**. See attached Excel files for **Table S1-S7**, **Table S9-S11** and attached compressed files for **Other supplementary materials 1-5**.

#### The legend of Supplementary Materials

Table S8

**Table S8.** Product yields in different strains by HPLC/MS analysis

| Strains | Label | FQC relative concentration* | Compound 1 relative concentration* |
|---|---|---|---|
| TYJY81.1 | Δ*fmqC* | 0 | 39.826 |
| TYJY81.2 | Δ*fmqC* | 0 | 39.861 |
| TYJY81.3 | Δ*fmqC* | 0 | 40.401 |
| TYJY82.1 | *fmqC* (control) | 199.258 | 0 |
| TYJY82.2 | *fmqC* (control) | 230.414 | 0 |
| TYJY82.3 | *fmqC* (control) | 270.880 | 0 |
| TYJY83.1 | G409A | 225.824 | 0 |
| TYJY83.2 | G409A | 221.679 | 0 |
| TYJY83.3 | G409A | 203.156 | 0 |
| TYJY84.1 | G409R | 198.312 | 0 |
| TYJY84.2 | G409R | 245.373 | 0 |
| TYJY84.3 | G409R | 195.700 | 0 |
| TYJY85.1 | G409D | 27.931 | 28.286 |
| TYJY85.2 | G409D | 37.183 | 23.218 |
| TYJY85.3 | G409D | 31.764 | 23.974 |
| TYJY86.1 | G409P | 6.403 | 30.642 |
| TYJY86.2 | G409P | 6.214 | 36.839 |
| TYJY86.3 | G409P | 8.460 | 41.718 |
| TYJY87.1 | G409W | 13.351 | 39.269 |
| TYJY87.2 | G409W | 13.653 | 40.303 |
| TYJY87.3 | G409W | 13.613 | 26.148 |
| TYJY88.1 | G409Y | 18.441 | 32.551 |
| TYJY88.2 | G409Y | 13.979 | 25.974 |

*Concentration related to the compound in the 18 minutes in LC-MS result (Figure 4E).

Table S12

**Table S12.** Fungal plasmids and strains used in this study

| Strains | Description | Reference |
|---|---|---|
| **Cea 17.1** | *Δku80* | [1] |
| **Cea 17.2** | *Δku80*, *pyrG1* | [1] |
| **TYJY81** | Δ*fmqC*::*hyg* | This study |
| **TYJY82** | *fmqC*::*pyrG* | This study |
| **TYJY83** | *FmqC409*$^{gly->ala}$::*AfpyrG*, *pyrG1* | This study |
| **TYJY84** | *FmqC409*$^{gly->arg}$::*AfpyrG*, *pyrG1* | This study |
| **TYJY85** | *FmqC409*$^{gly->asp}$::*AfpyrG*, *pyrG1* | This study |
| **TYJY86** | *FmqC409*$^{gly->pro}$::*AfpyrG*, *pyrG1* | This study |
| **TYJY87** | *FmqC409*$^{gly->trp}$::*AfpyrG*, *pyrG1* | This study |
| **TYJY88** | *FmqC409*$^{gly->tyr}$::*AfpyrG*, *pyrG1* | This study |
| **pYH-wA-pyrG** | *URA3, WA flanking, AfpyrG, Amp* | [2] |
| **pYJY25** | *FmqC*::*AfpyrG* in pYH-wA-pyrG | This study |
| **pYJY26** | *FmqC409*$^{gly->ala}$::*AfpyrG* in pYH-wA-pyrG | This study |
| **pYJY27** | *FmqC409*$^{gly->arg}$::*AfpyrG* in pYH-wA-pyrG | This study |
| **pYJY28** | *FmqC409*$^{gly->asp}$::*AfpyrG* in pYH-wA-pyrG | This study |
| **pYJY29** | *FmqC409*$^{gly->pro}$::*AfpyrG* in pYH-wA-pyrG | This study |
| **pYJY30** | *FmqC409*$^{gly->trp}$::*AfpyrG* in pYH-wA-pyrG | This study |
| **pYJY31** | *FmqC409*$^{gly->tyr}$::*AfpyrG* in pYH-wA-pyrG | This study |

TXX = original transformant, pXX = plasmid

Table S13

**Table S13.** PCR primers used in this study.

| **Primer** | **Oligonucleotide sequences (5'-3')** | **Uses** |
|---|---|---|
| Hyg-FOR | cattccaatcgataccgtcgac | *G418* amplification |
| Hyg-REV | gtggataaccgtattaccgcc | |
| fmqC-5F-F | cacttcaagtctgccaaagcg | *fmqC* 5′ flanks amplification |
| fmqC-5F-R | caatatcagttaacgtcgacggtatcgattggaatgtcgatcaagtgtcgttgtctccg | |
| fmqC-3F-F | atcagctcactcaaaggcggtaatacggttatccacgaagggacctgagataggtttgt | *fmqC* 3′ flanks amplification |
| fmqC-3F-R | aacctggcaatcatgacgat | |
| fmqC-NEST-F | tcagcacacccttaccgaag | *fmqC* deletion cassette amplification |
| fmqC-NEST-R | acaaatccttgcgctgttct | |
| fmqC-RT-F | gttctgcacctgtgcactc | *fmqC* transformant screening |
| fmqC-RT-R | ggtgcatcggcttcatcttc | |
| pyrG-F | gagagttattctgtgtctgacgaaa | *pyrG* amplification |
| pyrG-R | attctgtctgagaggaggcac | |
| fmqC-WA-F | ctccttctcctgatcgataggtatcctgcatgtgtgcg | construct mutant plasmids |
| fmqC-pyrG-R | cacagaataactctcagtgcgatatatttgtgctgct | |
| pyrG-3F-F | cctctcagacagaataattccgtttctttcctgcgg | construct mutant plasmids |
| 3F-WA-R | gtgattcgcgtcatgcggccgccaatcaacactgctctaccgac | |
| fmqC-G to A-F | gatcgcattccccgtcagcgtggtaccctgggtagtgg | FmqC409$^{gly->ala}$ |
| fmqC-G to A-R | cgctgacggggaatgcgatcttgctcggatgactc | |
| fmqC-G to R-F | gatcagattccccgtcagcgtggtaccctgggtagtgg | FmqC409$^{gly->arg}$ |
| fmqC-G to R-R | cgctgacggggaatctgatcttgctcggatgactc | |
| fmqC-G to D-F | gatcgacttccccgtcagcgtggtaccctgggtagtgg | FmqC409$^{gly->asp}$ |
| fmqC-G to D-R | cgctgacggggaagtcgatcttgctcggatgactc | |
| fmqC-G to P-F | gatcccattccccgtcagcgtggtaccctgggtagtgg | FmqC409$^{gly->pro}$ |
| fmqC-G to P-R | cgctgacggggaatgggatcttgctcggatgactc | |
| fmqC-G to W-F | gatctggttccccgtcagcgtggtaccctgggtagtgg | |
| fmqC-G to W-R | cgctgacggggaaccagatcttgctcggatgactc | FmqC409$^{gly->trp}$ |
| fmqC-G to Y-F | gatctacttccccgtcagcgtggtaccctgggtagtgg | |
| fmqC-G to Y-R | cgctgacggggaagtagatcttgctcggatgactc | FmqC409$^{gly->tyr}$ |

Figure S1

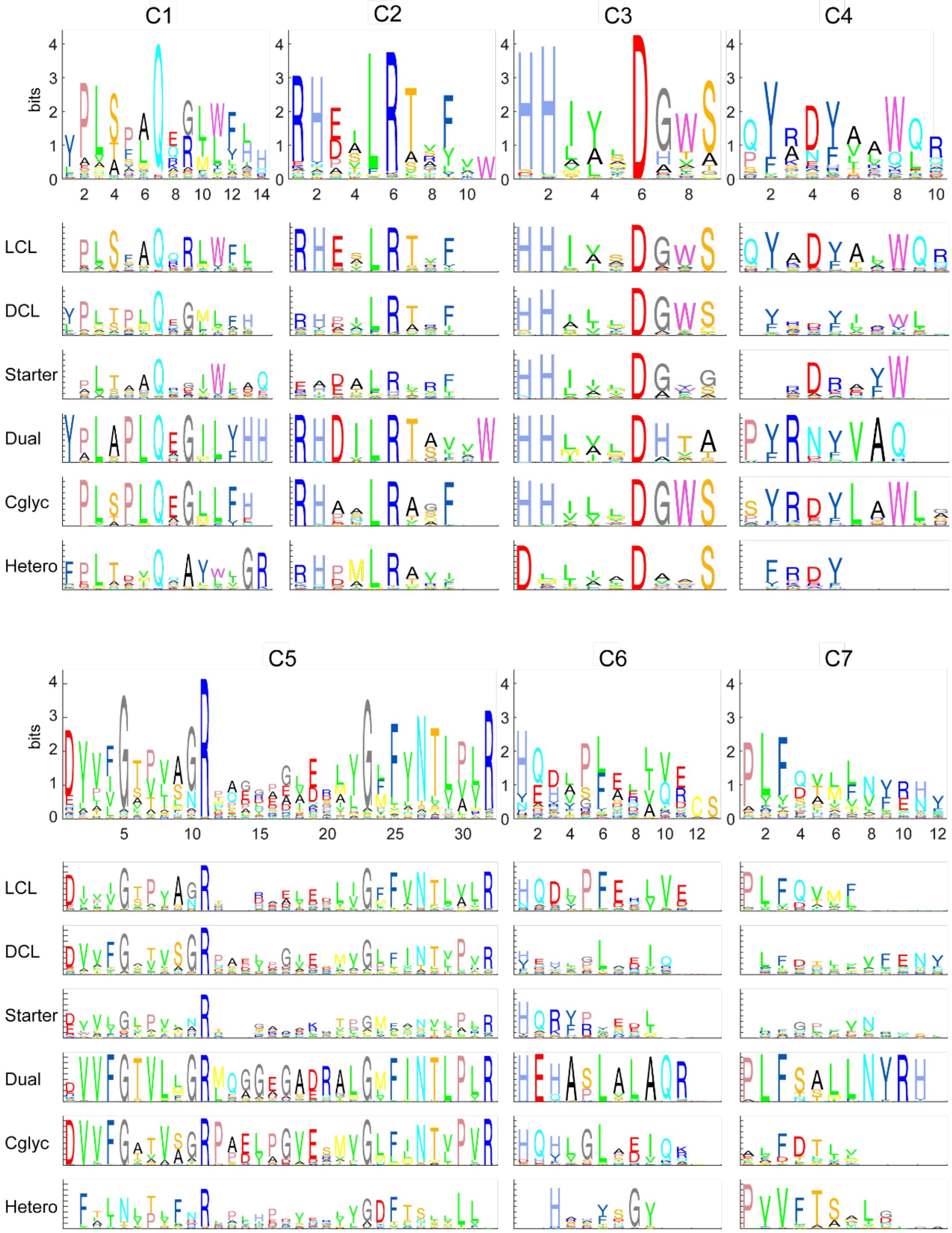


**Figure S1. Sequence logo of the seven C motifs among the multialignment of 1758 C domains (first row), and among six subtypes of C domains (second row) in MiBiG**
The ranges of y-axis in sequence logo figures all are 0~4.4 bits. The numbers of each C domain subtypes are 809 (LCL), 385 (DCL), 114 (Starter), 300 (Dual), 114 (Cgly), and 36 (Heterocyclization).

Figure S2

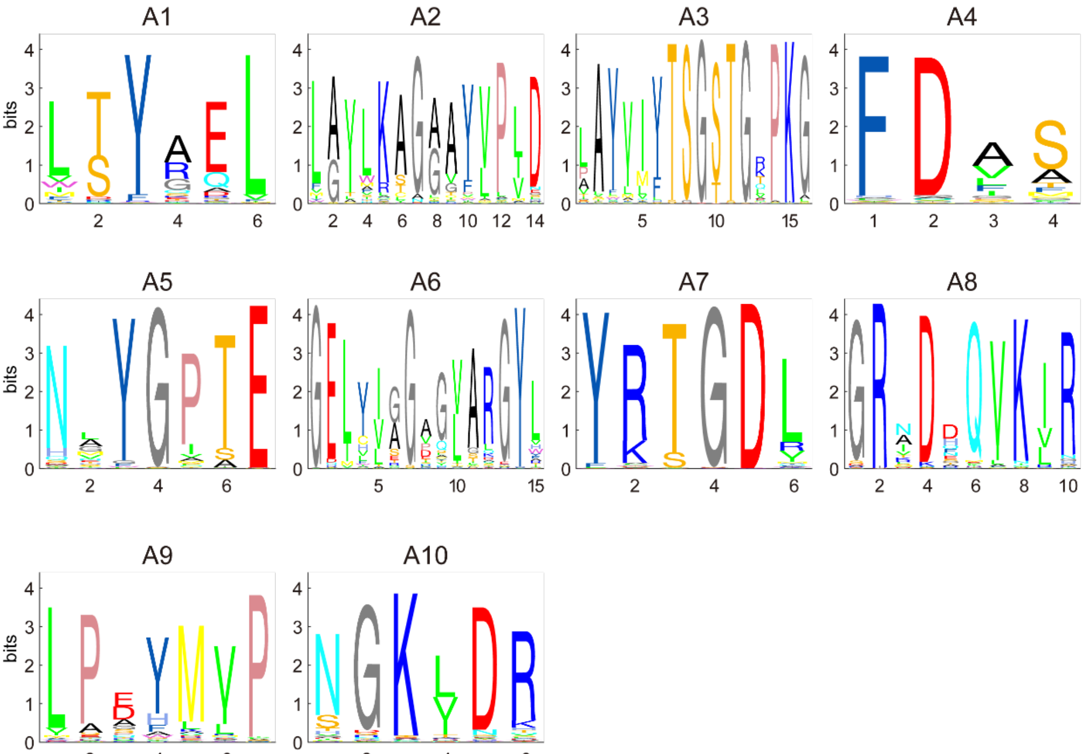


**Figure S2. Sequence logo of the ten A motifs among the multialignment of 1859 A domains in MiBiG**

The y-axis ranges in sequence logo figures all are 0~4.4 bits.

Figure S3

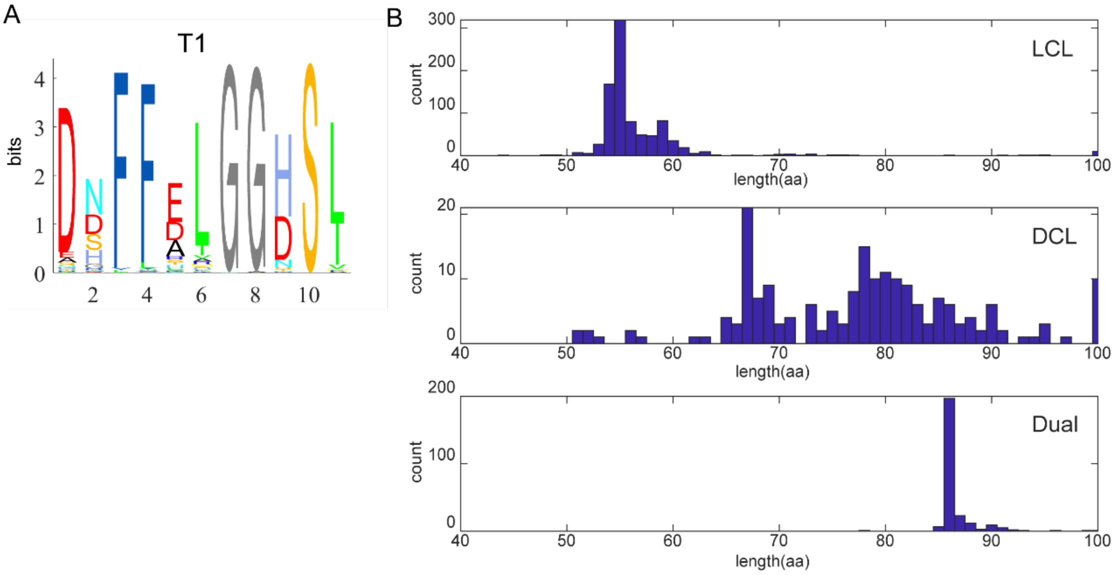


**Figure S3. Sequence logo of T1 motif and the length distribution of the T1-C1 region in MiBiG**

**A.** Sequence logo of the T1 motif. The y-axis range in sequence logo figure is 0~4.4 bits.

**B.** Length distribution of the T1-C1 region, for different subtypes of C.

Figure S4

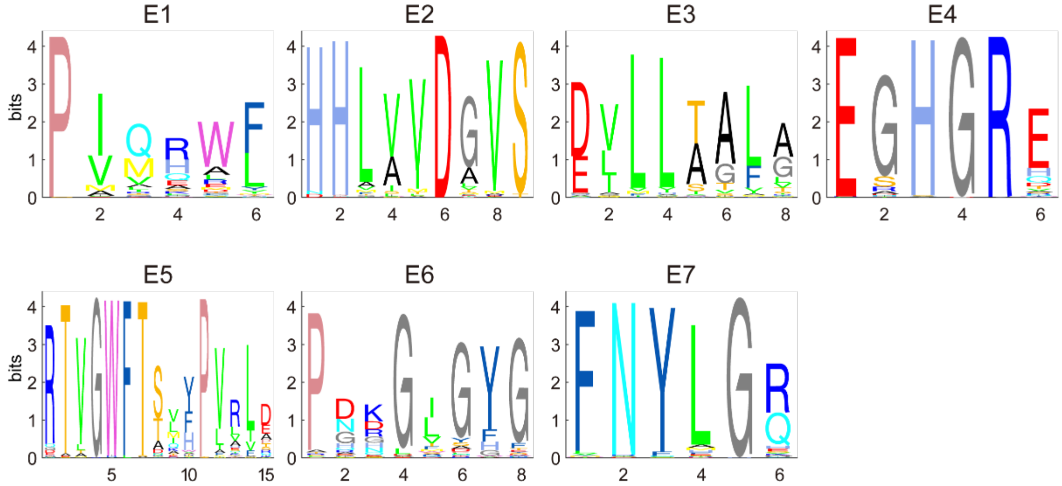


**Figure S4. Sequence logo of the seven E motifs among the multialignment of 310 E domains in MiBiG**

The y-axis ranges in sequence logo figures all are 0~4.4 bits.

Figure S5

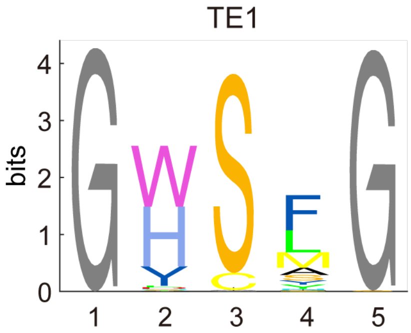


**Figure S5. Sequence logo of the TE1 motifs among the multialignment of 280 TE domains in MiBiG**

The y-axis range in sequence logo figure is 0~4.4 bits.

Figure S6

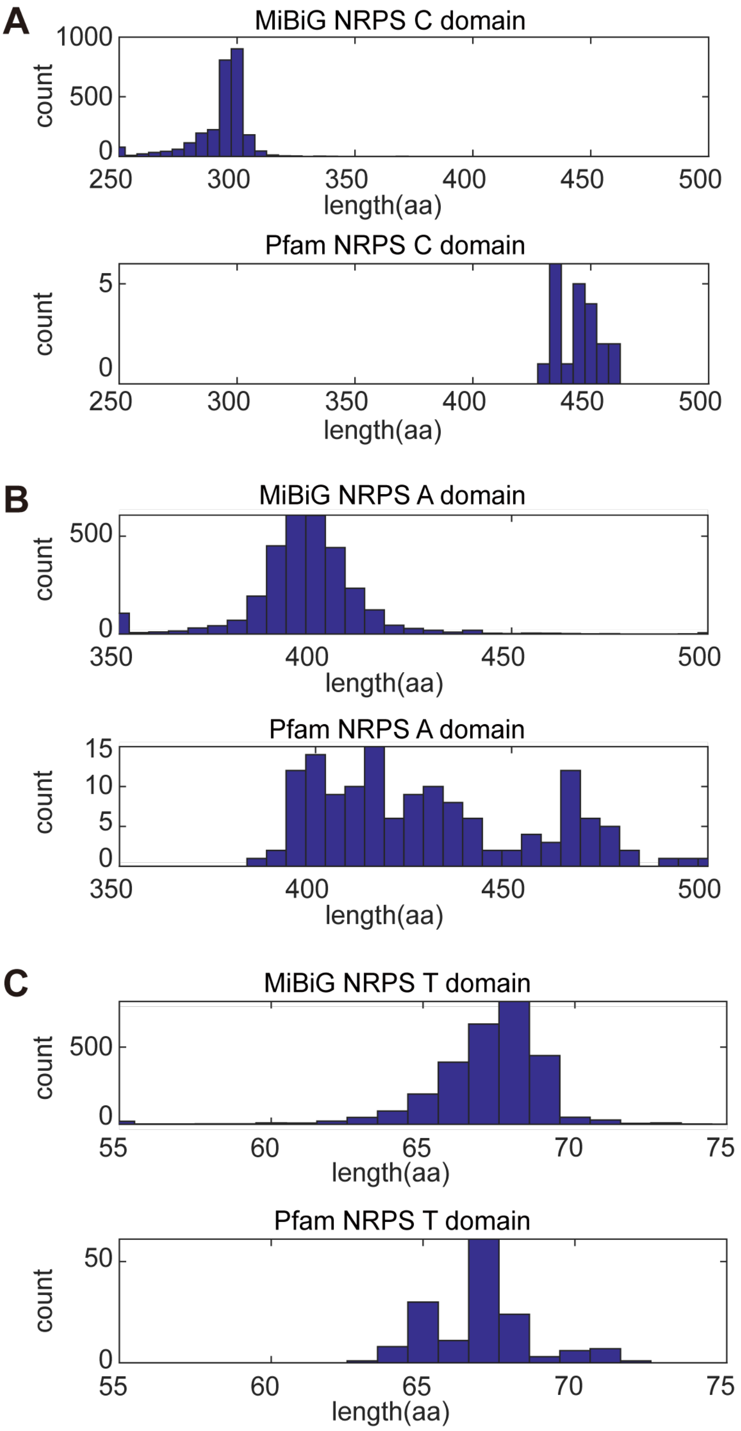


**Figure S6. Length distributions of C, A, and T domains, in MIBIG and in Pfam seeds**

Figure S7

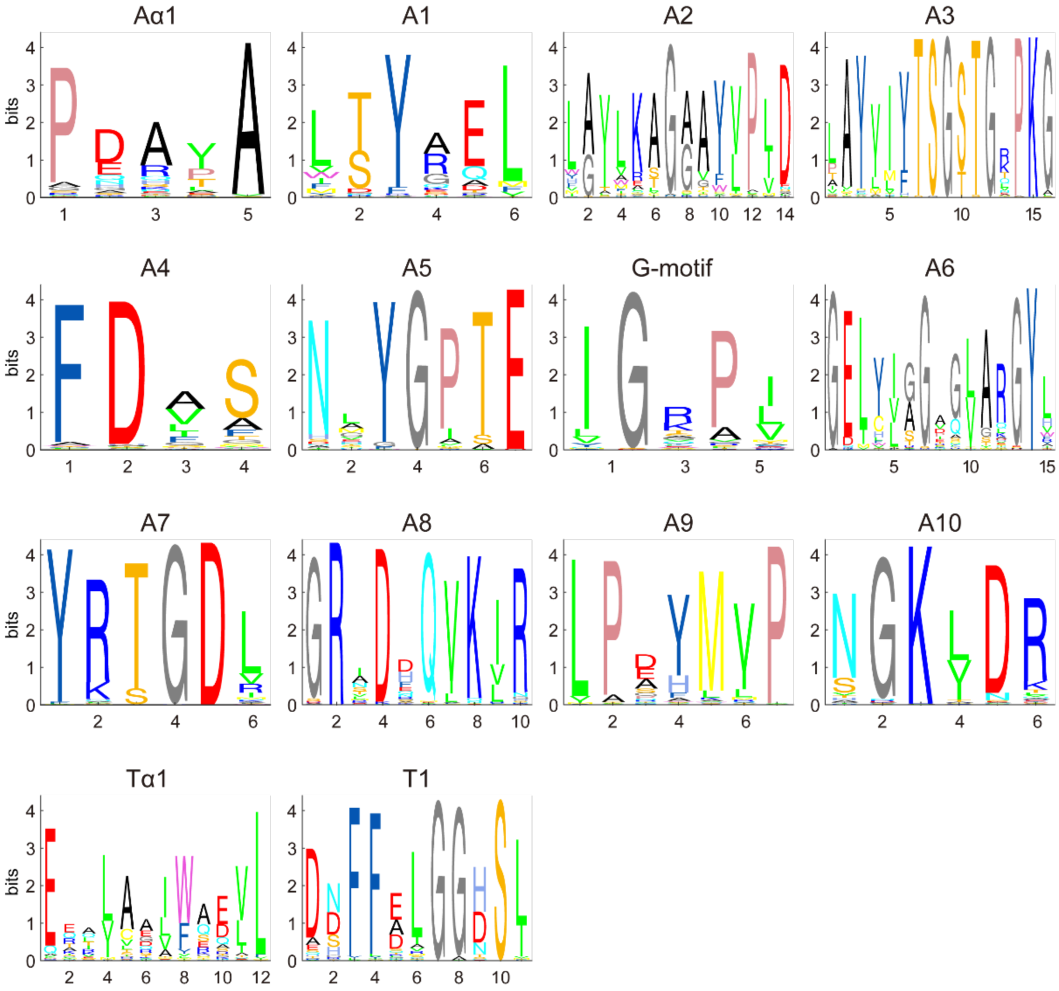


**Figure S7. Sequence logo of the twelve A motifs and two T motifs among 95,582 A domains and 86,688 T domains in bacteria**

The y-axis ranges in sequence logo figures all are 0~4.4 bits.

Figure S8

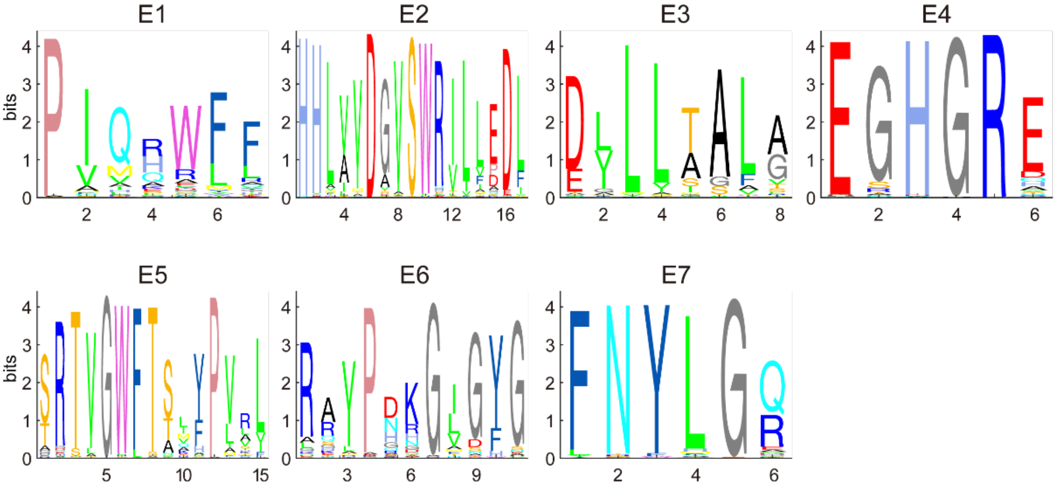


**Figure S8. Sequence logo of the seven E motifs among 14,502 E domains in bacteria**

The y-axis ranges in sequence logo figures all are 0~4.4 bits.

Figure S9

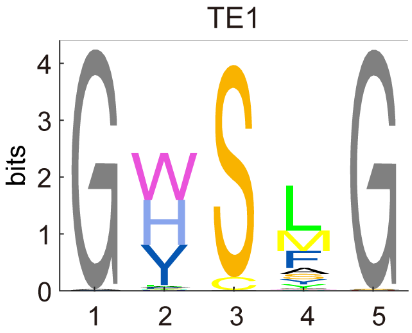


**Figure S9. Sequence logo of the TE1 motifs among 23,590 TE domains in bacteria**

The y-axis range in sequence logo figure is 0~4.4 bits.

Figure S10

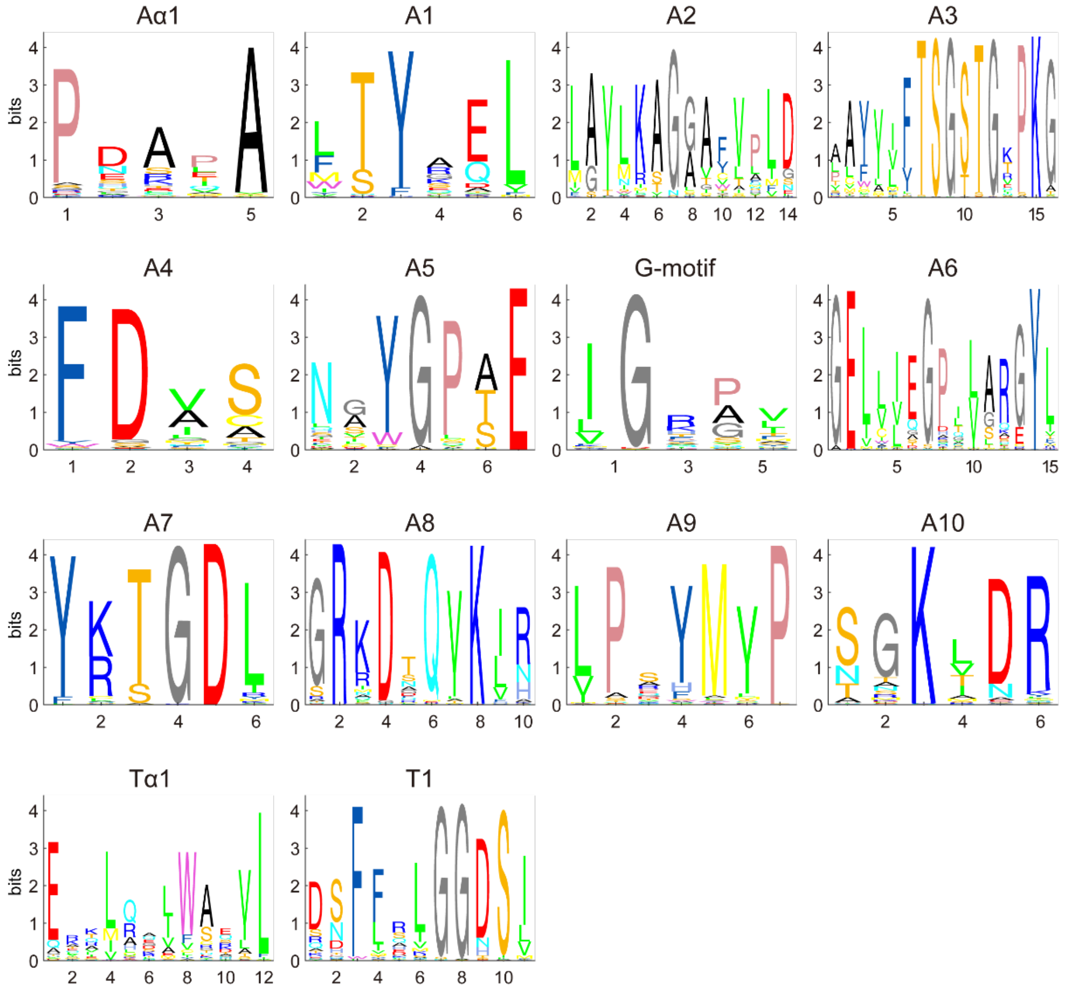


**Figure S10. Sequence logo of the twelve A motifs and two T motifs among 40,458 A domains and 26,651 T domains in fungi**

The y-axis ranges in sequence logo figures all are 0~4.4 bits.

Figure S11

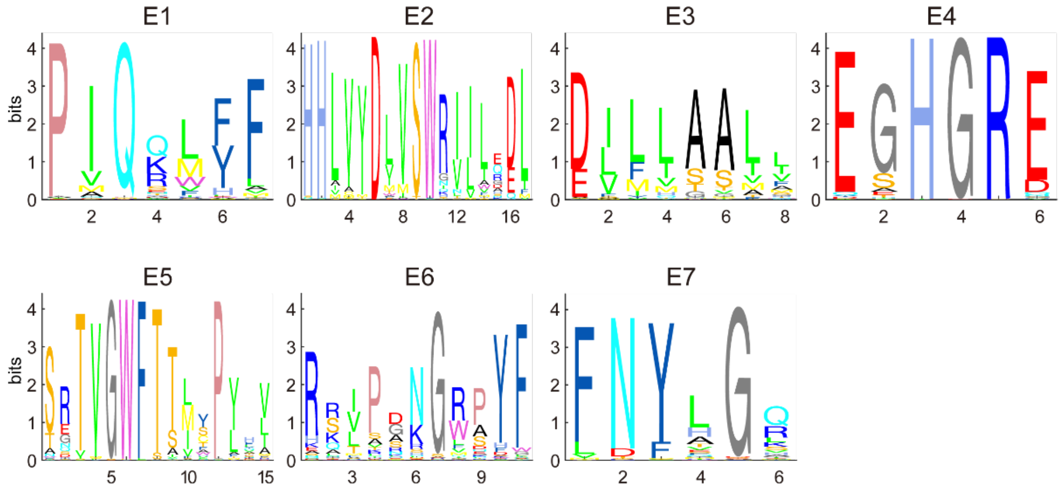


**Figure S11. Sequence logo of the seven E motifs among 3,982 E domains in fungi**

The y-axis ranges in sequence logo figures all are 0~4.4 bits.

Figure S12

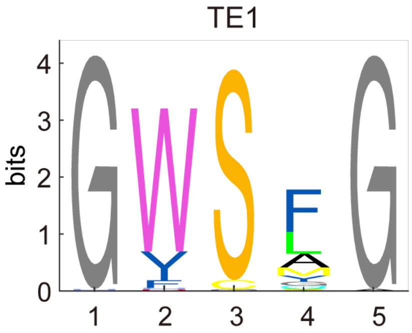


**Figure S12. Sequence logo of the TE1 motifs among 4,008 TE domains in fungi**

The y-axis range in sequence logo figures is 0~4.4 bits.

Figure S13

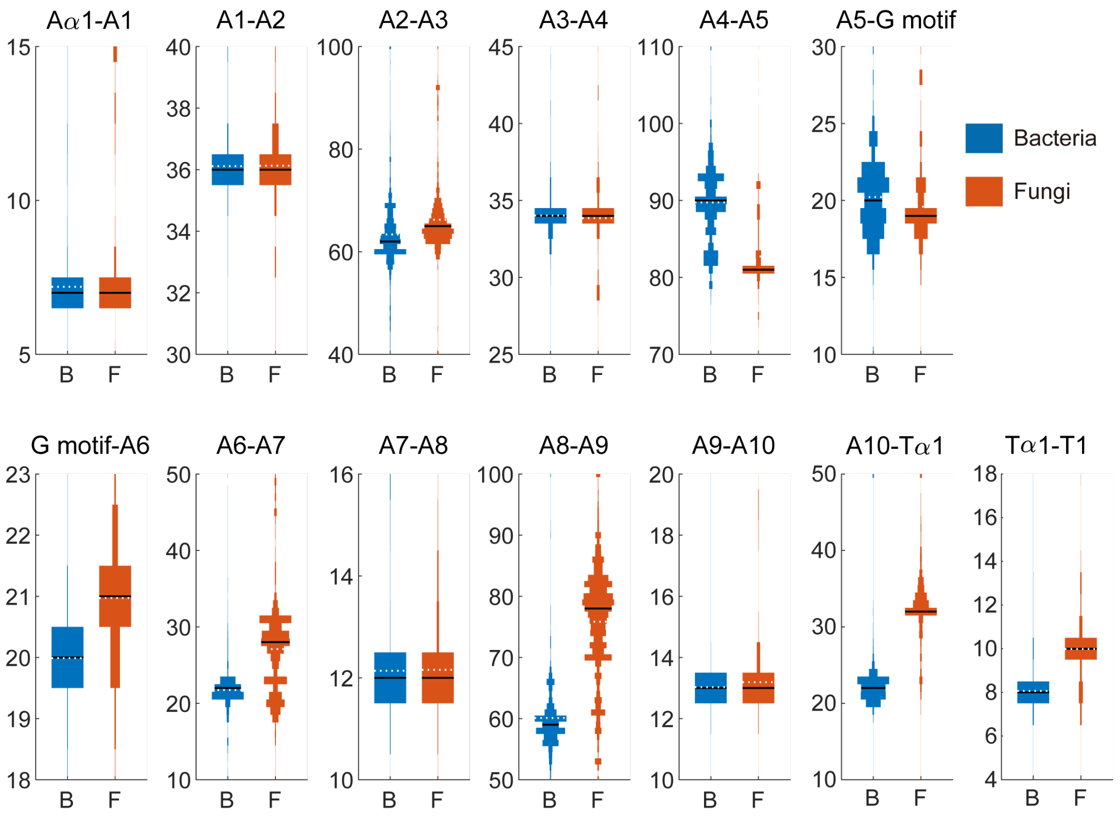


**Figure S13. Comparison of NRPS A and T domain architecture between bacteria and fungi**

For comparison, only A domains which have same motif length with reference A domain are used. Sequence numbers of intermotifs (A1-A2, A2-A3, A3-A4, A4-A5, A5-G motif, G motif-A6, A6-A7, A7-A8, A8-A9, A9-A10, Tα1-T1) are 75,407 in bacteria and 17,890 in fungi. Sequence numbers of A1-Tα1 intermotif (actually interdomain) in bacteria are 69,440 while they in fungi are 12,194 because only part of A domains are adjacent with the T domain. Sequence numbers of Tα1-T1 intermotif in bacteria are 85,755 while they in fungi are 25,069

Figure S14

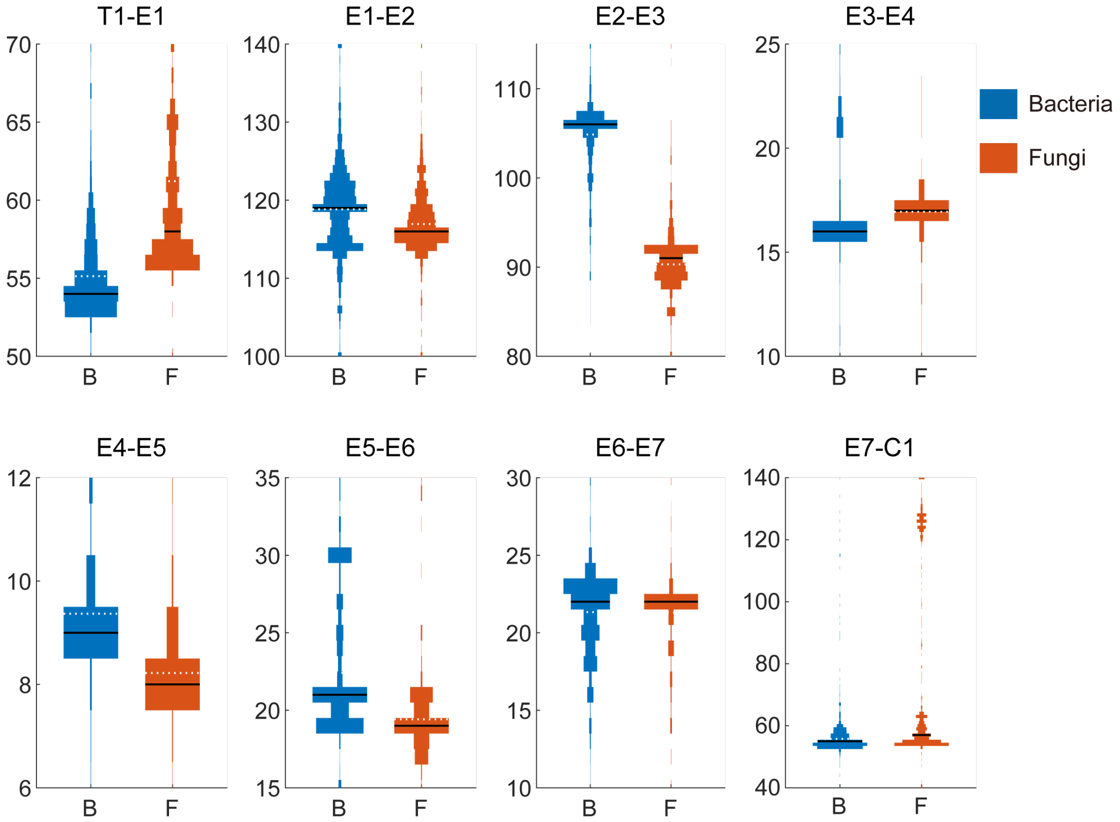


**Figure S14. Comparison of NRPS E domain architecture between bacteria and fungi**
For comparison, only A domains which have same motif length with reference A domain are used. Sequence numbers of intermotifs (E1-E2, E2-E3, E3-E4, E4-E5, E5-E6, E6-E7) are 12,875 in bacteria and 2,852 in fungi. Sequence numbers of intermotifs (actually interdomain) in bacteria are 12,618 for T1-E1 and 8,353 for E7-C1 while they in fungi are 2,088 for T1-E1 and 2,530 for E7-C1.

### Figure S15

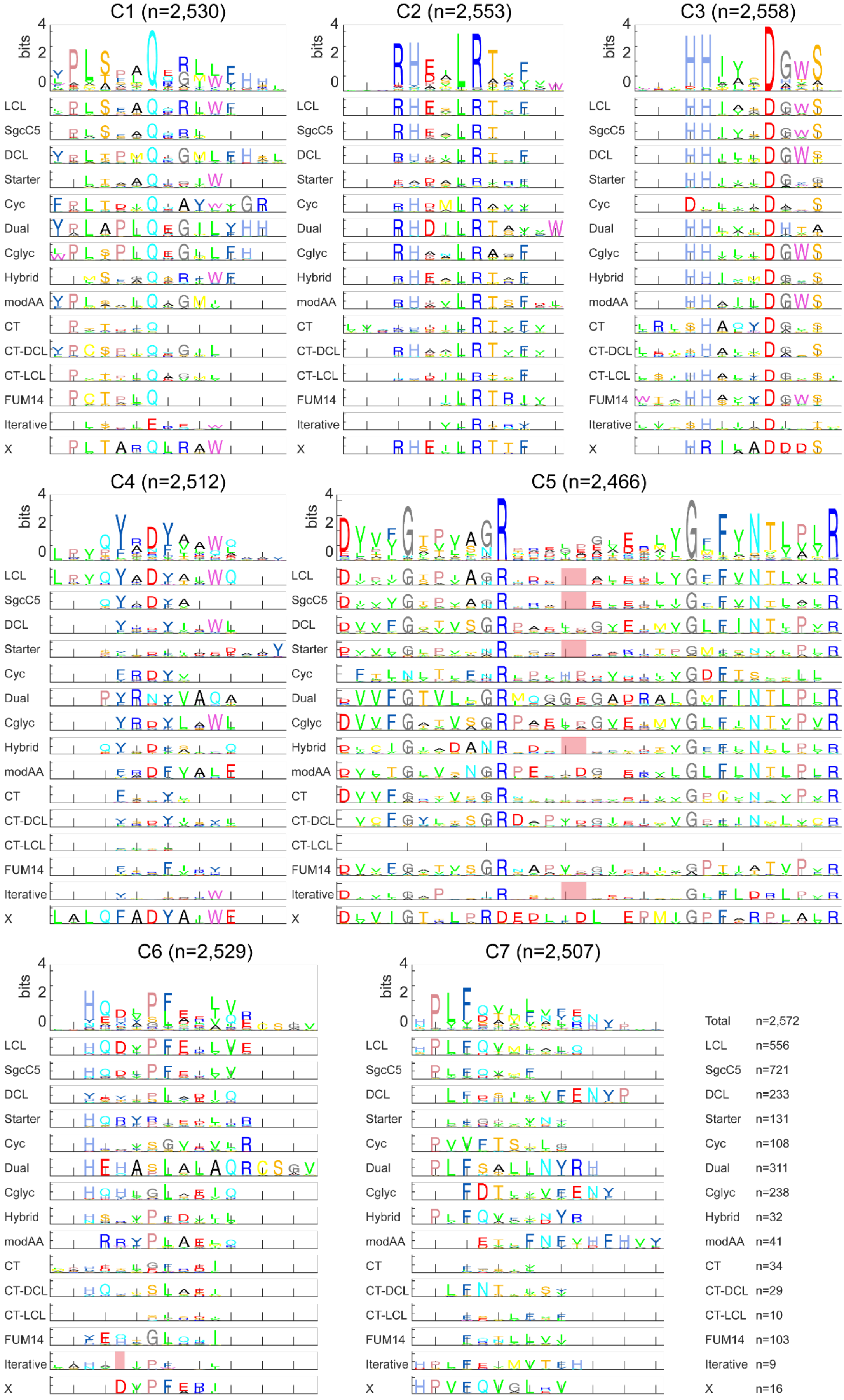
C1 (n=2,530)
C2 (n=2,553)
C3 (n=2,558)
C4 (n=2,512)
C5 (n=2,466)
C6 (n=2,529)
C7 (n=2,507)
bits
LCL
SgcC5
DCL
Starter
Cyc
Dual
Cglyc
Hybrid
modAA
CT
CT-DCL
CT-LCL
FUM14
Iterative
X
Total n=2,572
LCL n=556
SgcC5 n=721
DCL n=233
Starter n=131
Cyc n=108
Dual n=311
Cglyc n=238
Hybrid n=32
modAA n=41
CT n=34
CT-DCL n=29
CT-LCL n=10
FUM14 n=103
Iterative n=9
X n=16

**Figure S15. Sequence logo of the seven C motifs among the multialignment of 2,572 C domains (first row), and among 15 subtypes of C domains (second row) in MiBiG** There are 2,572 C domains with subtype prediction scores more than the threshold (200) and a count of domain subtype sequences of more than 3. The y-axis ranges in sequence logo figures all are 0~4.4 bits. The numbers of each C domain subtype are labeled at the end. For clarification, only motifs that have prevalent length are used in plotting. The actual total C domain number used in the figure for the specific motif is shown in the title. There are some gaps at both ends of the sequence for alignment. In motif C5, there are some interior gaps labeled in red for aligning with motif C5 of other subtypes.

#### Figure S16

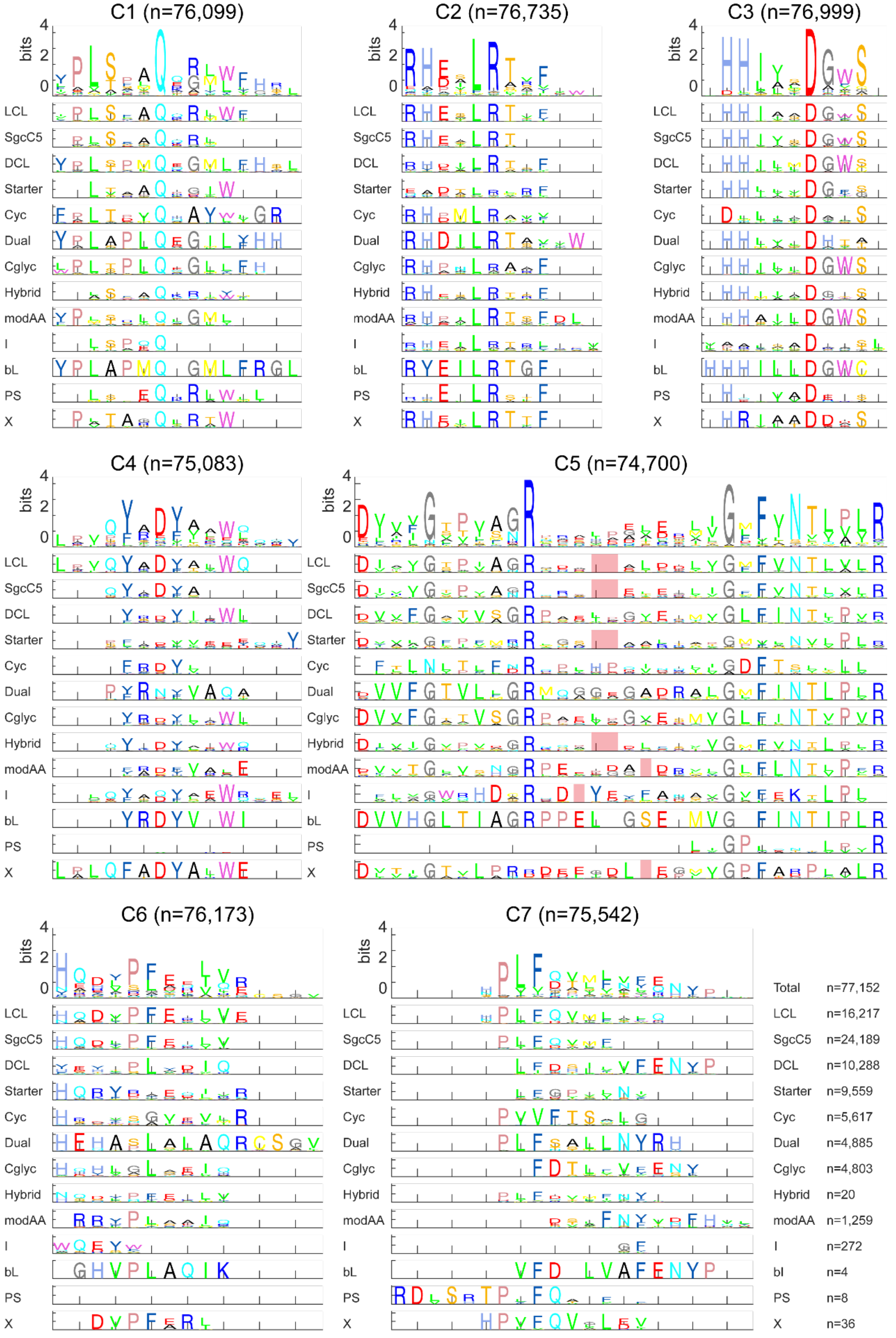


**Figure S16. Sequence logo of the seven C motifs among 77,152 C domains (first row), and among 13 subtypes of C domains (second row) in bacteria**

There are 77,152 C domains with subtype prediction scores more than the threshold (200) and a count of domain subtype sequences of more than 3. The y-axis ranges in sequence logo figures all are 0~4.4 bits, except it's 0~1 for bL the subtype and 0~2.1 for the PS subtype because these subtypes are few in the sequence number. The numbers of each C domain subtype are labeled at the end. For clarification, only motifs that have prevalent length are used in plotting. The actual total C domain number used in the figure for the specific motif is shown in the title. There are some gaps at both ends of the sequence for alignment. In motif C5, there are some interior gaps labeled in red for aligning with motif C5 of other subtypes.

#### Figure S17

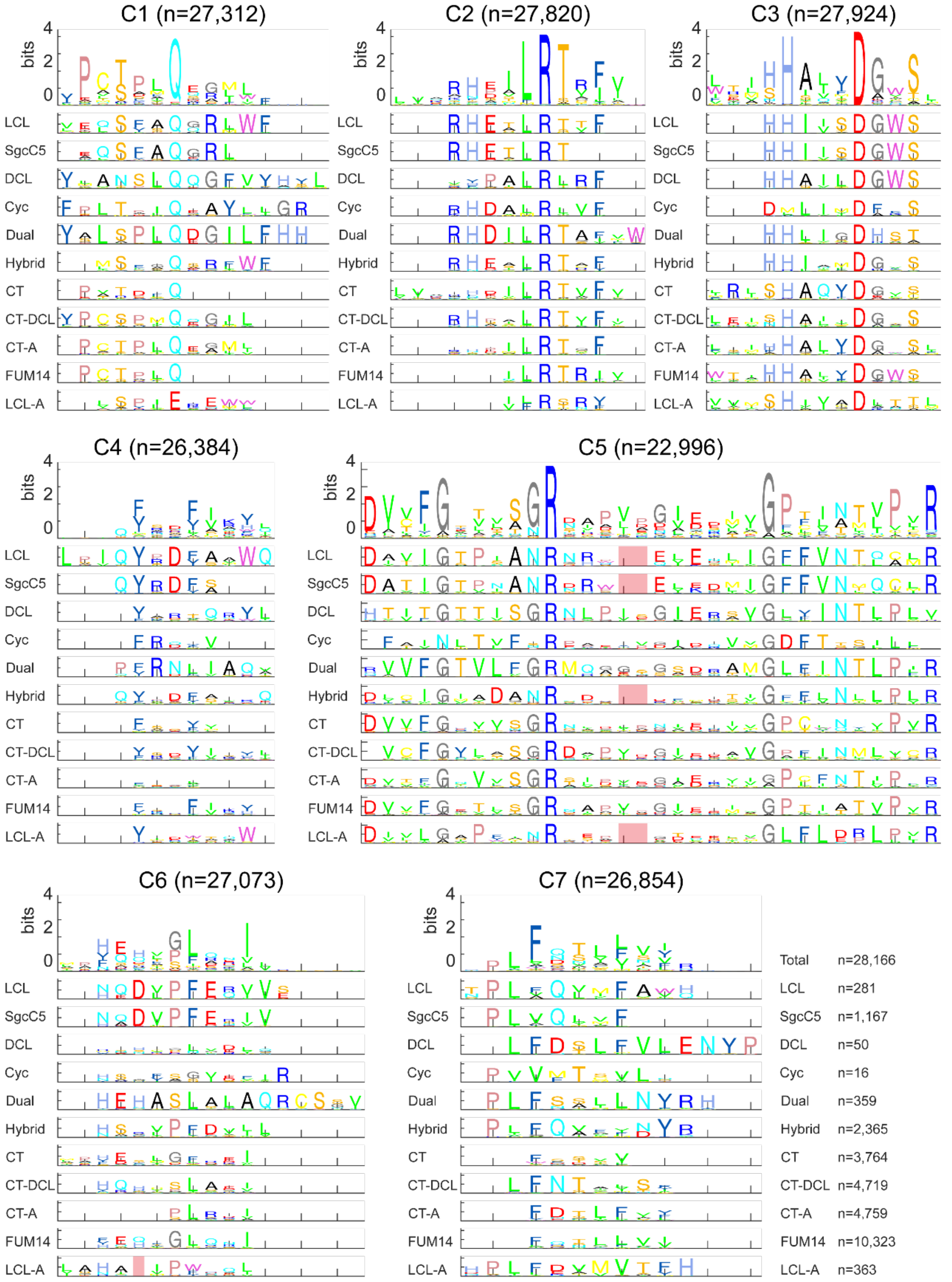


**Figure S17. Sequence logo of the seven C motifs among 34,269 C domains (first row), and among 11 subtypes of C domains (second row) in fungi**

There are 34,269 C domains with subtype prediction scores more than the threshold (200) and a count of domain subtype sequences of more than 3. The y-axis ranges in sequence logo figures all are 0~4.4 bits. The numbers of each C domain subtype are labeled at the end. For clarification, only motifs that have prevalent length are used in plotting. The actual total C domain number used in the figure for the specific motif is shown in the title. There are some gaps at both ends of the sequence for alignment. In motif C5, there are some interior gaps labeled in red for aligning with motif C5 of other subtypes.

Figure S18

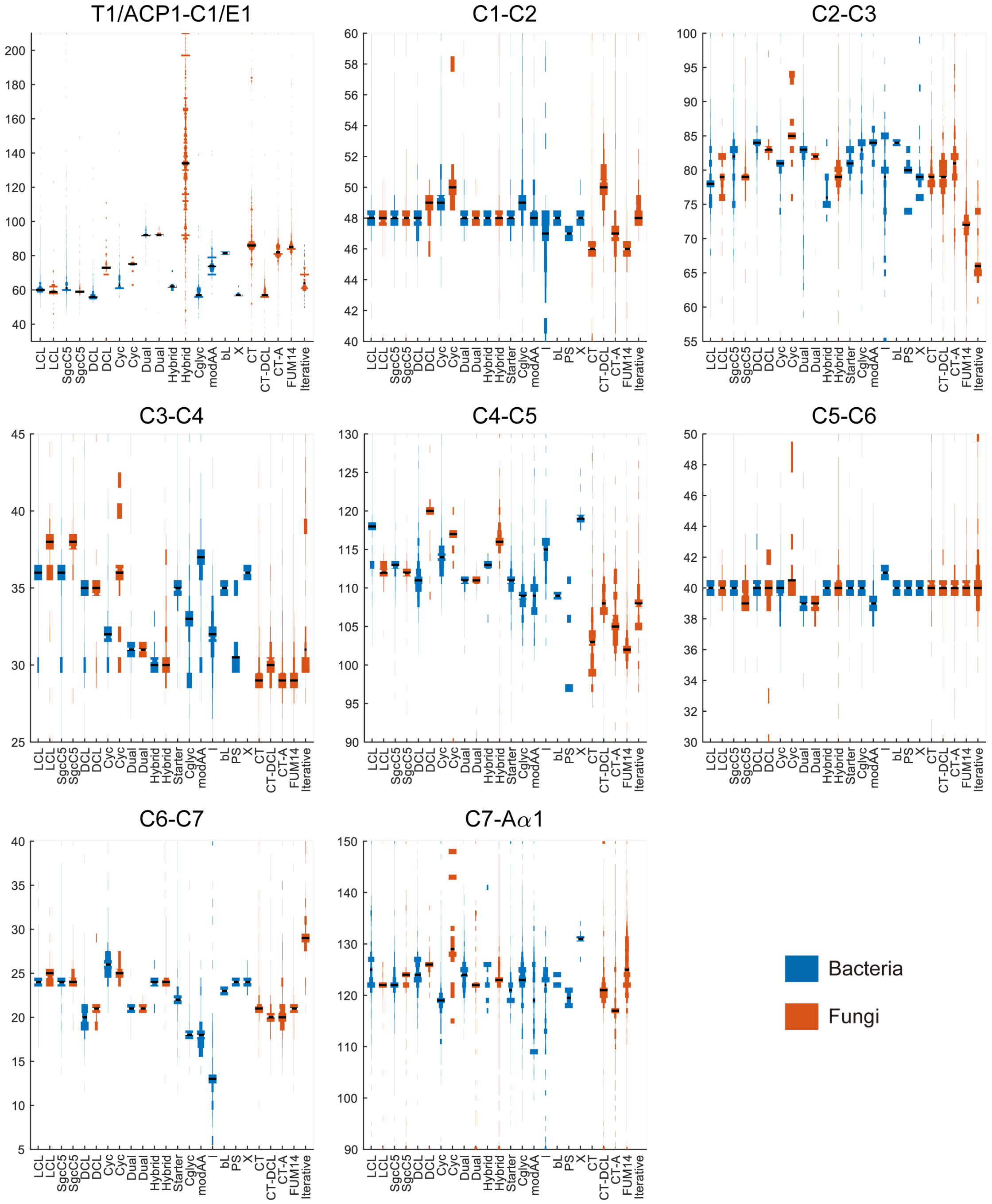


**Figure S18. Comparison of NRPS C domain architecture between bacteria and fungi**

For comparison of intermotif length between different C domain subtypes and E domain, we chose the conserved positions which exist in all C domain and E domain as start and end of intermotif. T1/ACP1-C1/E1 ends before the conserved “Q” in C1 and E1 (the conserved “E” in LCL-A subtype). C1-C2 starts with the conserved “Q” in C1 (the conserved “E” in LCL-A subtype), and ends before the second conserved “R” in C2. C2-C3 starts before the second conserved “R” in C2 and ends before the conserved “D” in C3. C3-C4 starts before the conserved “D” in C3 and ends before the second conserved “Y” in C4. C4-C5 starts before the second conserved “Y” in C4 and ends before the conserved “G” in C5. C5-C6 starts before the conserved “G” in C5 and ends before the conserved “P” in C6.

C6-C7 starts before the conserved “P” in C6 and ends before the conserved “F” in C7 (the conserved “F” in LCL). C7-Aα1 starts before the conserved “F” in C7 (the conserved “F” in LCL) and ends before Aα1. Sequence numbers of intermotifs (C1-C2, C2-C3, C3-C4, C4-C5, C5-C6, C6-C7) are 77,152 in bacteria and 34,269 in fungi. Sequence numbers of intermotifs (actually interdomain) in bacteria are 28,185 for T1-C1 and 33,176 for C7-Aα1 in LCL subtype C domain, 6,967 for E7-C1 and 6,752 for C7-Aα1 in DCL subtype C domain, 3,860 for T1-C1 and 4,495 for C7-Aα1 in Dual subtype C domain, 6,967 for T1-C1 and 6,752 for C7-Aα1 in starter subtype C domain and while they in fungi are 434 for T1-C1 and 616 for C7-Aα1 in LCL subtype C domain, 808 for E7-C1 and 1,931 for C7-Aα1 in DCL subtype C domain and 24 for T1-C1 and 21 for C7-Aα1 in Dual subtype C domain.

Figure S19

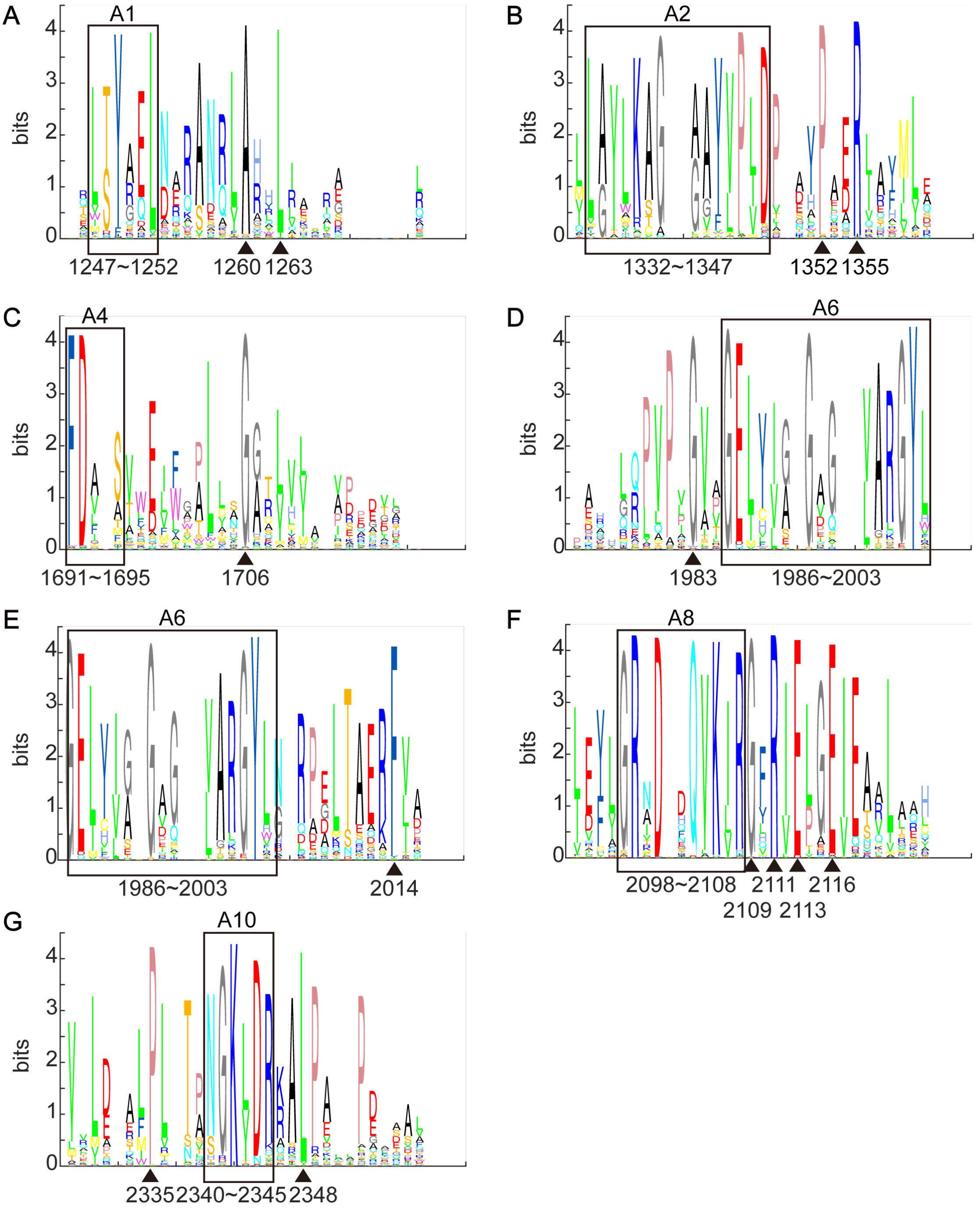


**Figure S19. Sequence logo of highly conserved positions near known motifs**

Black box shows known core motifs in A domain. Black triangle shows highly conserved positions in multialignment from the 1,161 C+A+T NRPS sequences from MiBiG database.

Figure S20

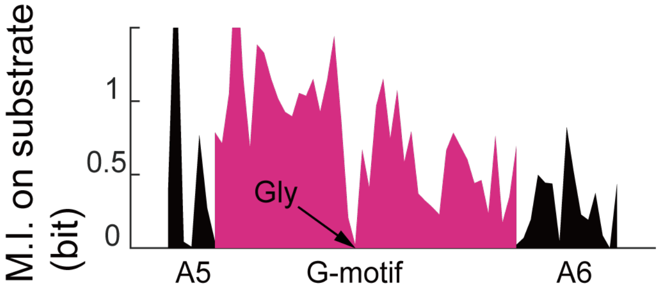


**Figure S20. The mutual information between residues in the A5-A6 and A domain substrate specificity**

Same as that in the fourth panel of Figure 1B, but this plot focuses on regions between motif A5 and motif A6. The position of conserved Gly is indicated by the black arrow.

Figure S21

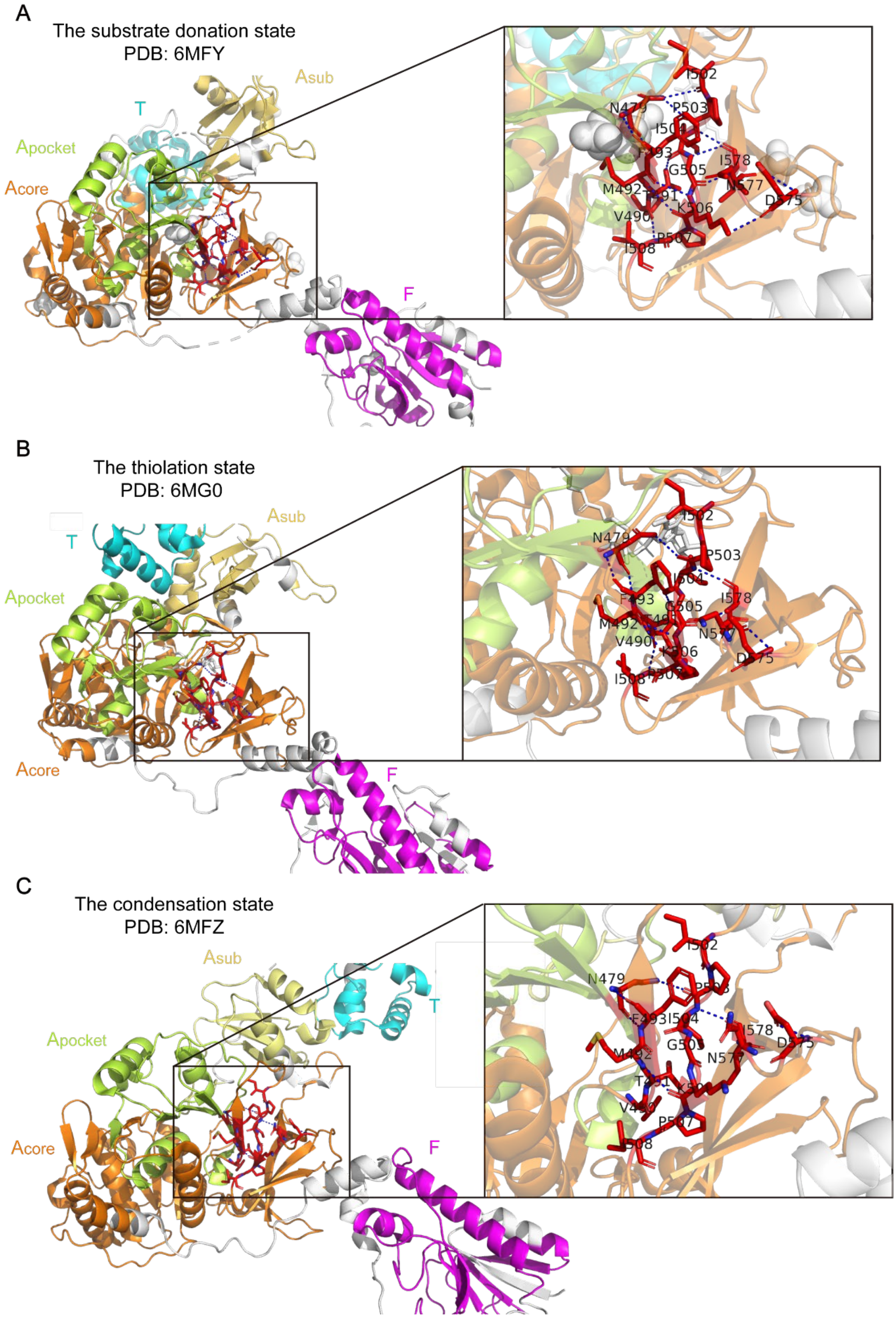


**Figure S21. The G-motif in different function states of LgrA structure**

Related to Figure 3B-3D. The view zoomed in towards the region near the G-motif emphasized by red sticks. Hydrogen bonds were shown in blue dashed-line. Different domains marked by different colors (F: formylation domain, colored by magenta; T: thiolation domain, colored by cyan; A: adenylation domain, $A_{core}$ (orange) covers A1-A8 of A domain, $A_{sub}$ (yellow) covers A9-A10, $A_{pocket}$ (yellow green) covers A3-A6 of A domain).

Figure S22

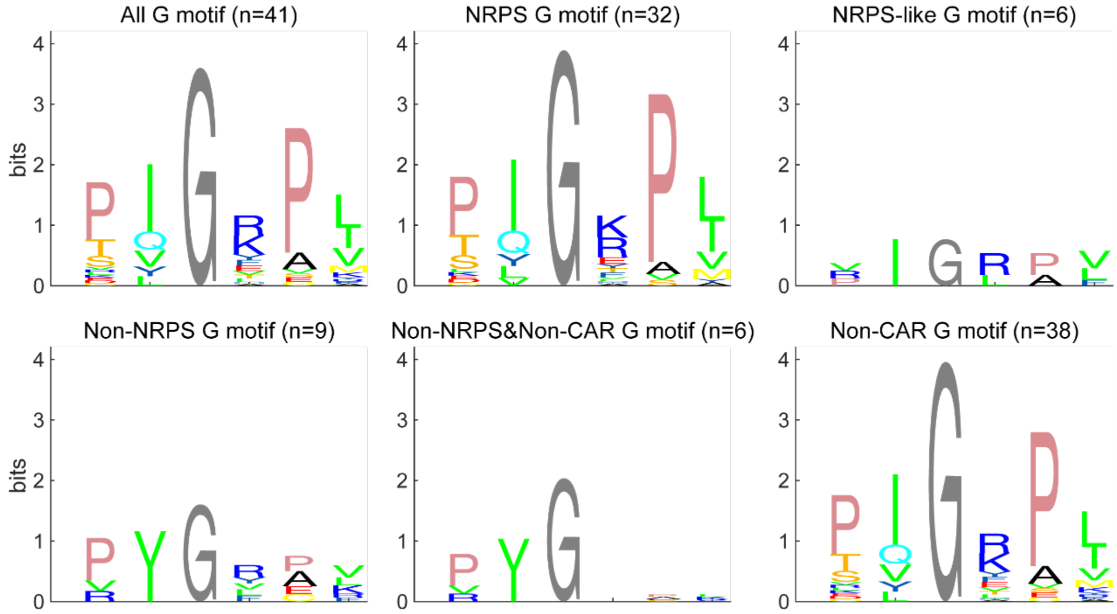


**Figure S22. The sequence logo of G-motif in A domains with known structures and FmqC**

Non-NRPS means A domains from these proteins which aren't NRPS. Non-CAR means A domains from these proteins which aren't CAR. Non-NRPS&Non-CAR means A domains from these proteins which aren't NRPS or CAR.

Figure S23

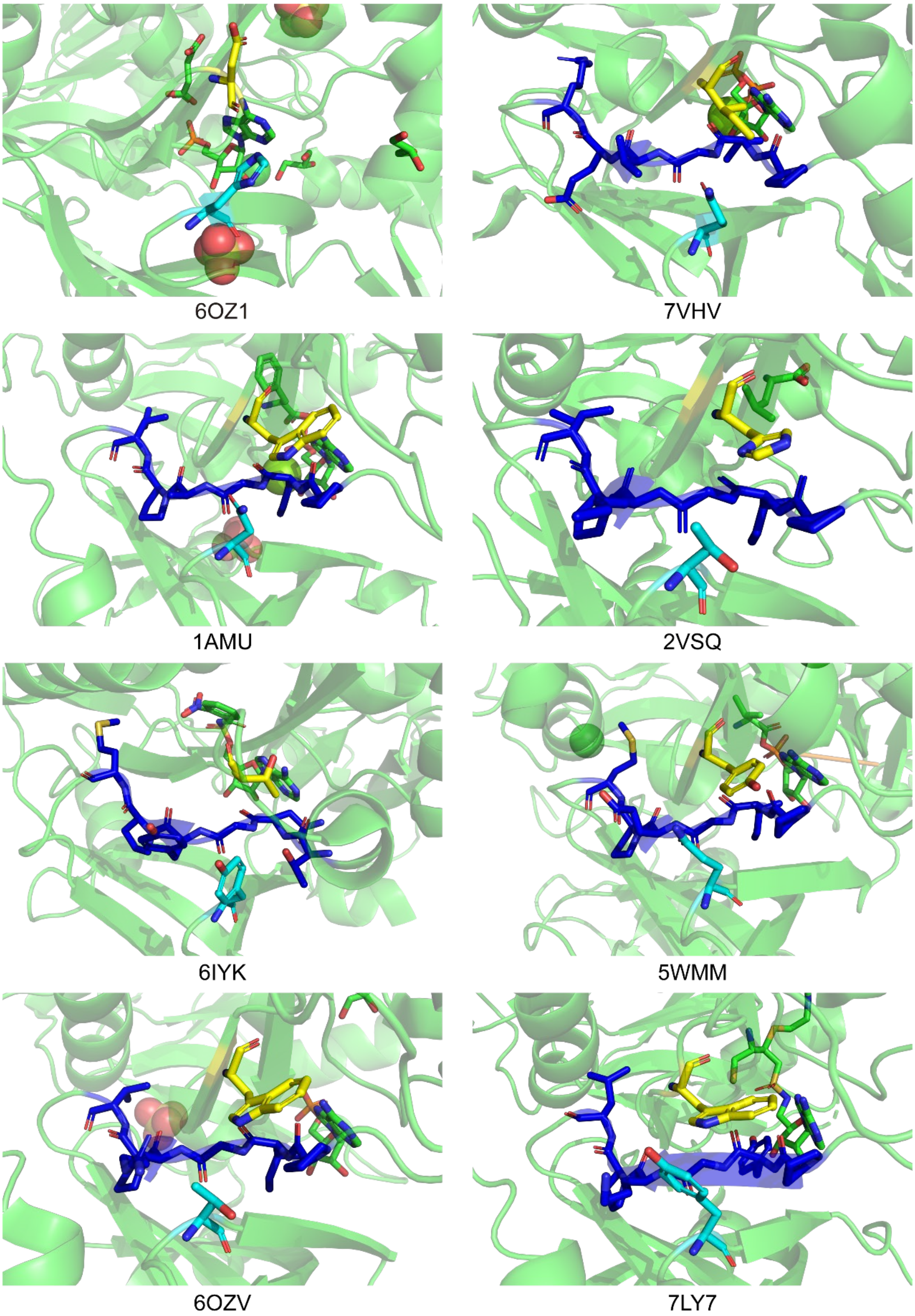


**Figure 23. The equivalents of N397, G409 (G-motif), and S491 in FmqC mapped to known structures**

From the first to the 8-th figure, the known structures are CAR protein (PDB: 6OZ1), DltA (PDB: 7VHV), and NRPS (PDB: 1AMU, 2VSQ, 6IYK, 5WMM, 6OZV and 7LY7). Residues in G-motif were marked by blue. The equivalent N397 and S491 were marked by yellow and cyan. The ligand molecule was marked in green. The PDB IDs of proteins are shown below. Only the CAR protein doesn't contain the G-motif, but it still has equivalents of N397 and S491. G-motif is in close proximity to the adenylate part of ligand, suggesting a potential gatekeeper role.

Figure S24

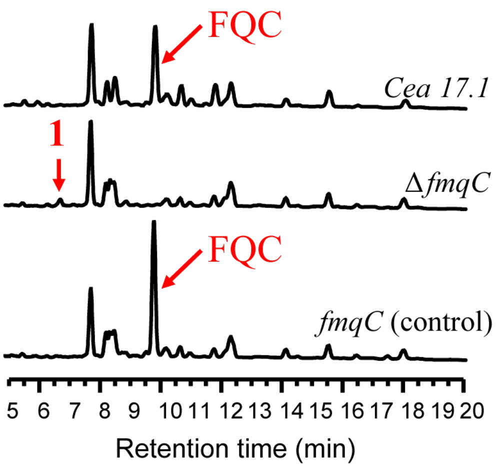


**Figure S24. LC-MS analysis in the strain construction**

Wild type (Cea17.2, first row), Δ*fmqC* (second row) and the control *fmqC* (third row)

Figure S25

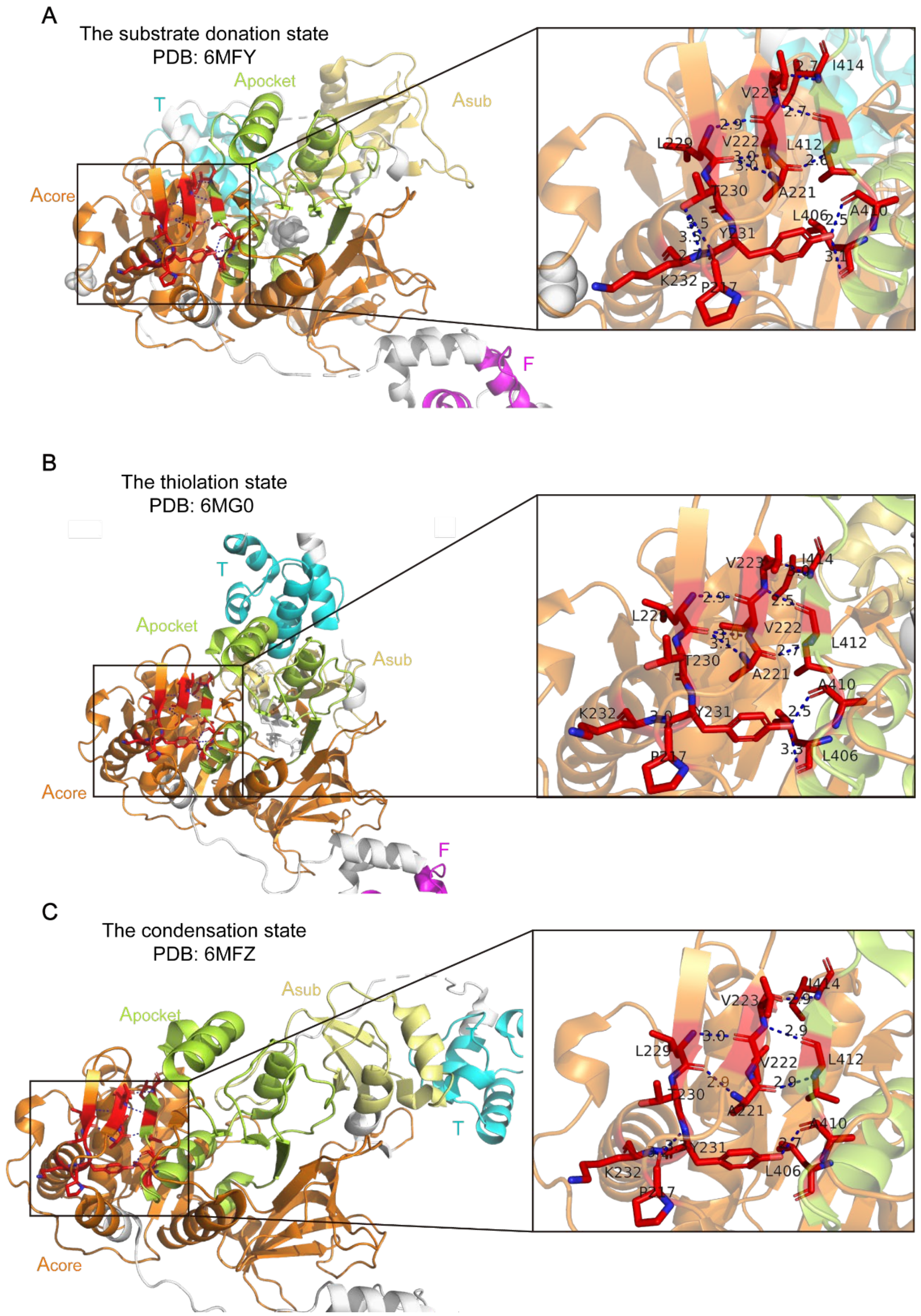


**Figure S25. Aα1 motif in different states of LgrA structure**

Similar with Figure S21, but for the Aα1 motif. The view zoomed in towards the region near the Aα1 motif emphasized by red sticks.

Figure S26

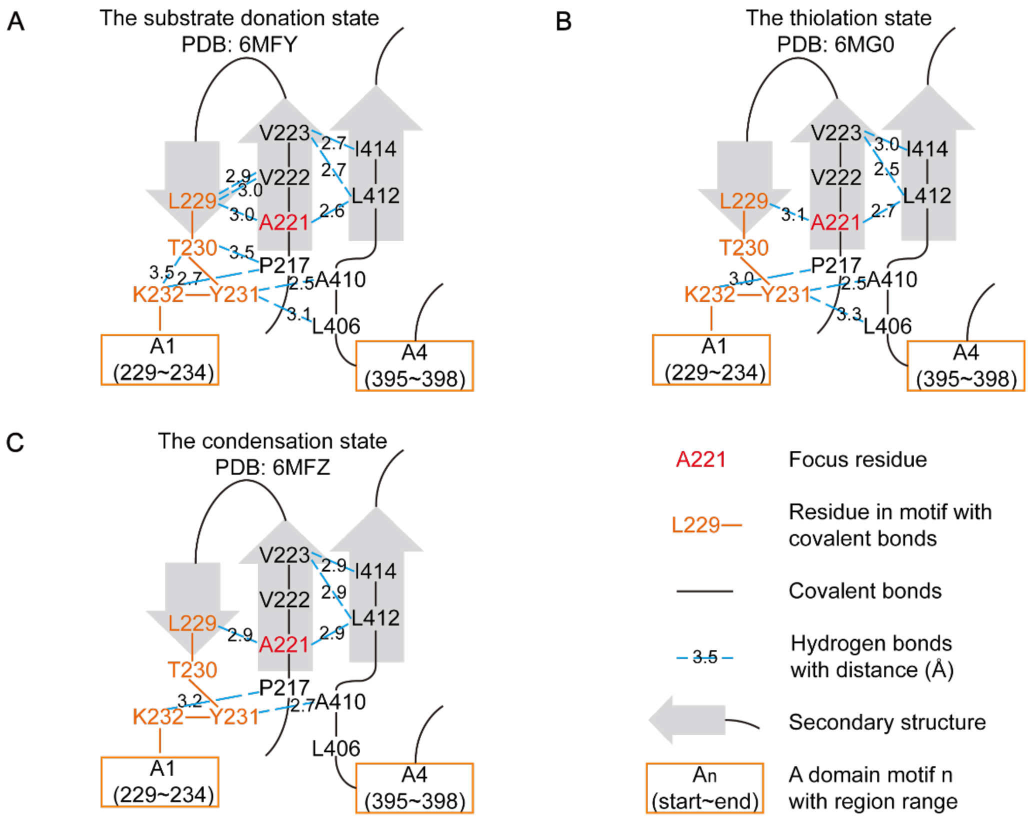


**Figure S26. The interaction near Aα1 motif in different states of LgrA structure**

Related to Figure S25. Similar with Figure 3B, but shows chemical interactions and secondary structures surrounding the Aα1 motif at the substrate donation state (A), the thiolation state (B) and the condensation state (C). Of note, in these residues, only T230 and Y231 use the hydroxyl group in the side chain to form hydrogen bonds. Other hydrogen bonds, on the other hand, are formed by the common α-carboxyl group and α-amino group in the main chains.

Figure S27

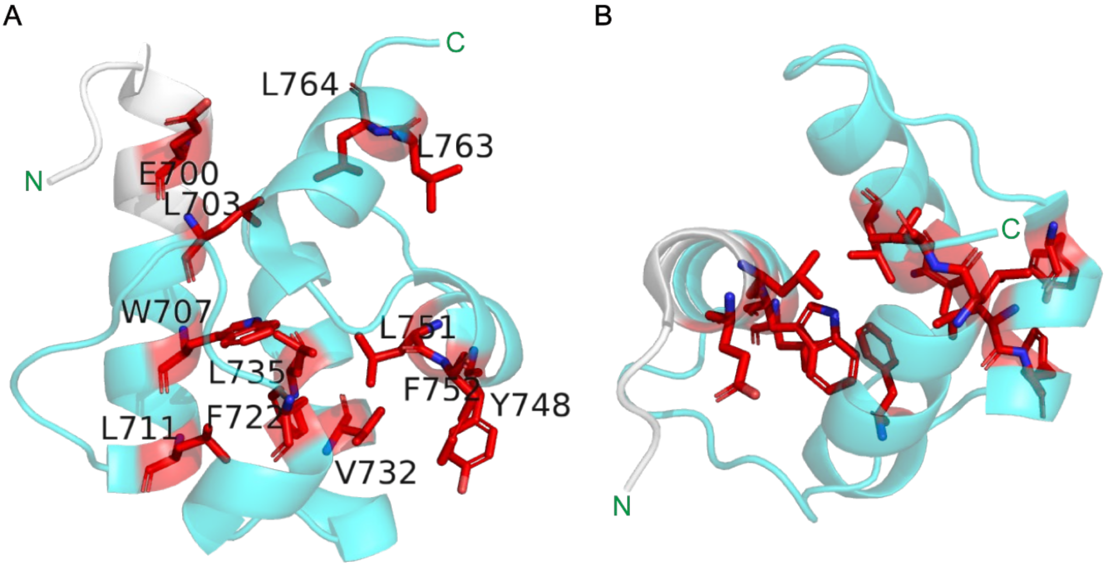


**Figure S27. Conserved resides of T domain in the condensation state**
Structure is obtained from LgrA in the condensation state (PDB: 6MFZ). Cyan color shows the T domain defined by Pfam. A small white region isn't covered by Pfam, although it is visually one part of the first helix of T domain. N-terminal and C-terminal were marked by green texts. Conserved resides were marked by red sticks with their one letter label shown. Reside labels in the top view were hidden for visual clearness.

**A.** Side view of T domain.

**B.** Top view of T domain.

#### Figure S28

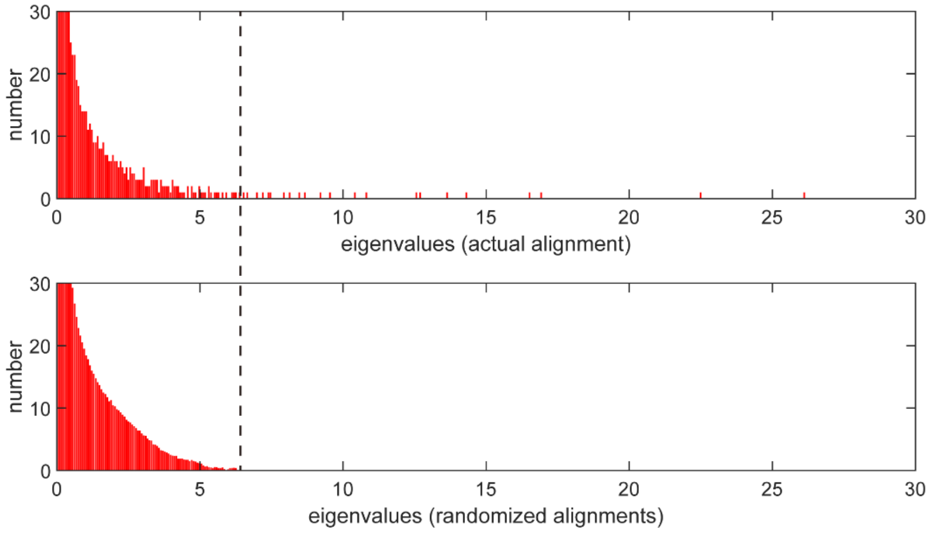


**Figure S28. Eigenvalue spectra for the SCA matrix of C+A+T modules MSA and random MSA**

Eigenvalue spectra for the SCA matrix corresponding to the 1,161 C+A+T modules (top panel) and for 100 trials of randomizing sequences alignment (bottom panel). The randomization process scrambles the order of amino acids in each alignment column independently, which did not change amino acid frequencies at positions. Black dashed-line marked the maximum of eigenvalues from randomized alignments. This analysis shows that only a small part of the spectrum (26 out of 2560 total eigenvalues) is significant given sample size.

Figure S29

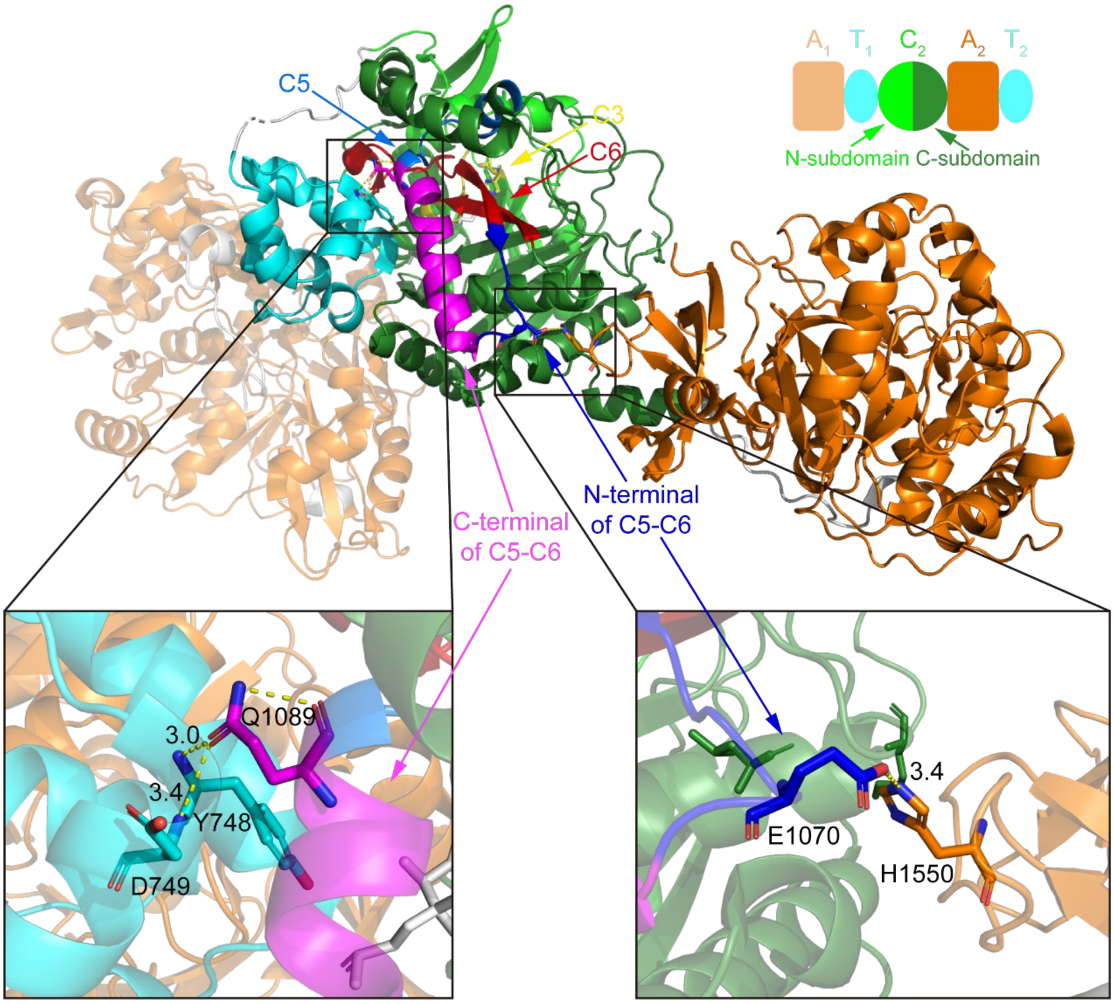


**Figure S29. The interaction of C5-C6 with other domains in LgrA structure**

The formylation domain in the first module (F1) of LgrA is hidden for visual clearness. The colors of each domain are noted on the top of figure. The C domain is split into the N-terminal subdomain (N-subdomain, covering C1-C4) and the C-terminal subdomain (C-subdomain, covering C5-C7), referring to previous research (PMID: 23756159). The active site histidine (the second histidine in C3 motif HHxxxD), the residues in C5-C6 intermotif interacting with T domain or A domain and the residues in T domain and A domain interacted by these residues are shown as stick format. Hydrogen bonds were shown in yellow dashed-line with distances by black.

#### Figure S30

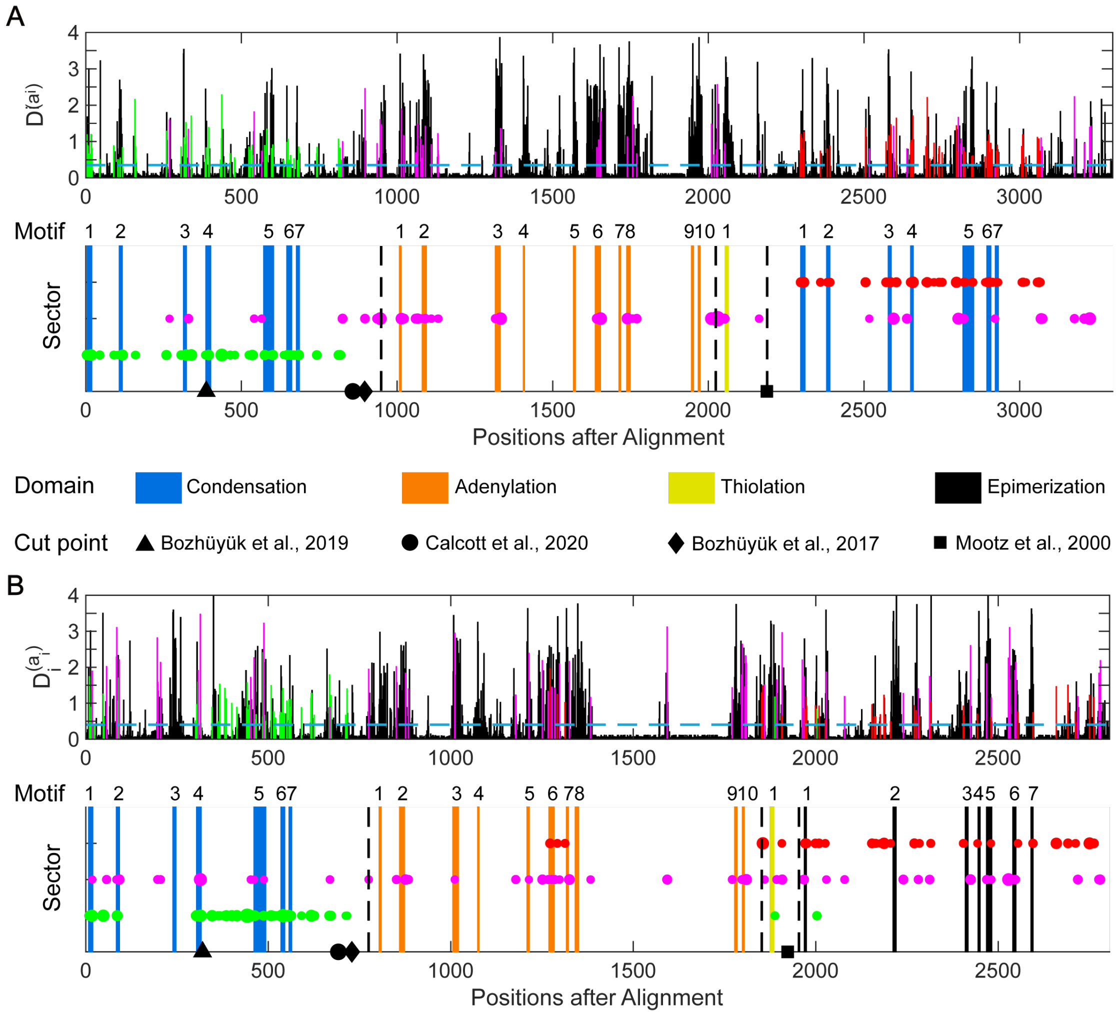


**Figure S30. SCA analysis for C+A+T+C and C+A+T+E four domains NRPS sequences** Similar with Figure 3A, we also analyzed 685 C+A+T+C (**A**) and 245 C+A+T+E (**B**) module NRPS sequences by SCA. Although sequence numbers are less than C+A+T composition NRPS (1,161), they also could provide insights for NRPS reengineering. Except known cutting points, the SCA result of C+A+T+E NRPS sequences indicated the junction between A and T domains may be a potential cutting point. Of note, cutting point proposed by Mootz et al.[3]. was for C+A+T+C in original research.

Figure S31

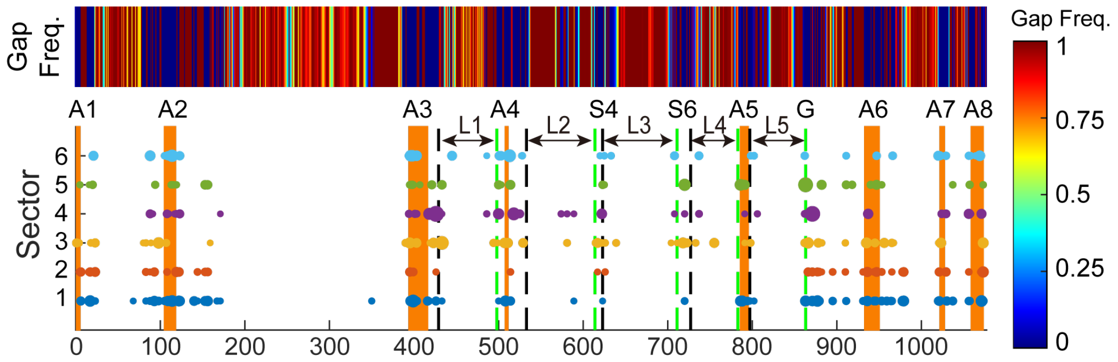


**Figure S31. The gap frequency in the MSA of 2,636 A domains**

Related to Figure 6A, upper panel shows the gap frequency in the MSA of 2,636 A domains. Some residues in the substrate-related sectors were found in the highly variable loop regions (L1-L5). These regions contain high numbers of gaps in MSA, and are usually loops in the structure.

Figure S32

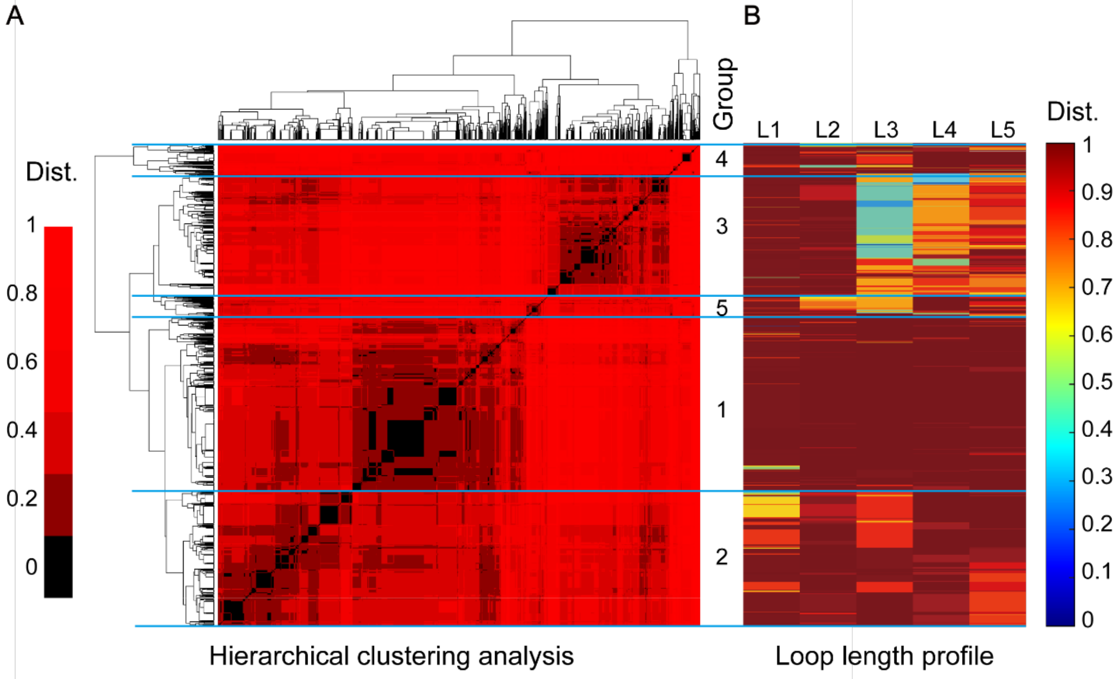


**Figure S32. Clustering and groups of five loops**

**A.** Hierarchical clustering of the A domains based on the Euclidean distances of their lengths in five loops. A domains were categorized into five groups based on their loop-length vectors. For visual clearness, in calculation, the Euclidean distances which are more than 12 are set as 12 before normalizing.

**B.** Loop length profiles of in five groups shown in **A**. More details in Method.

Figure S33

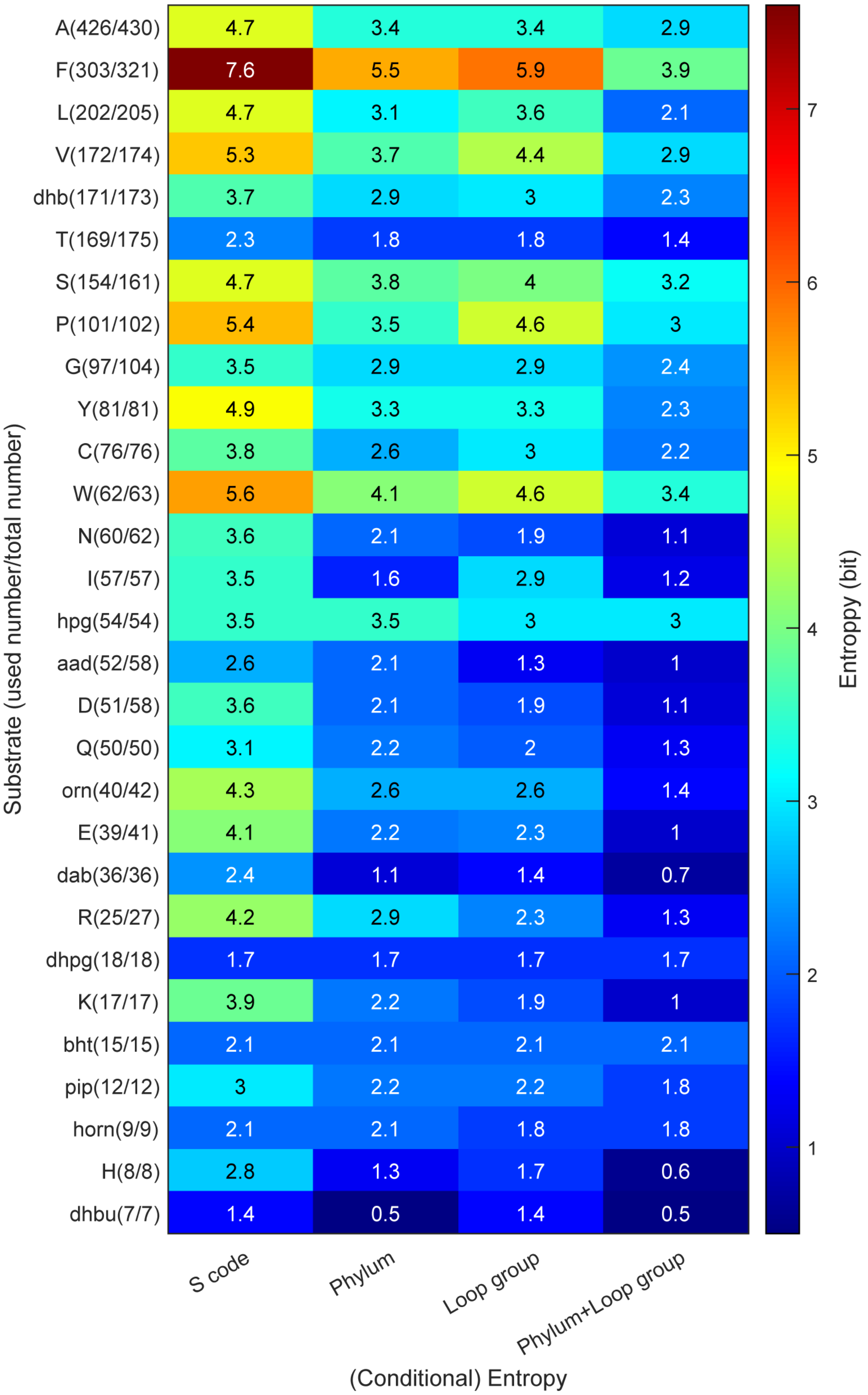


**Figure S33. The entropy and conditional entropy of the specificity-conferring code given different constraints.**

The first column shows the entropy of the specificity-conferring code for different substrates. The proteinogenic amino acids are named according to the standard amino acid one-letter code. Abbreviations of non-proteinogenic amino acid substrate: aad=2-amino-adipic-acid, bht=beta-hydroxy-tyrosine, dab=diaminobutyric acid, dhb=2,3-dihydroxy-benzoic acid, dhbu=2,3-dehydroaminobutyric acid, dhpg=3,5-dihydroxy-phenyl-glycin, horn=hydroxy-L-ornithine, hpg=4-hydoxy-phenyl-glycine, orn=ornithine and pip=pipecolic acid. Only A domains from 5 main phylum are used to calculate entropy (2564/2623=97.8% sequences). The sequence number of each substrate was marked in bracket after substrate names on the y labels (the first is the number used in calculation of entropy and the second is the total number of this substrate in our datasets). Second to fourth columns show the conditional entropy of the specificity-conferring code given information about the phylum, the loop group, and the phylum with loop group, respectively. Information from the phylum and the loop group both could reduce the uncertainty of the specificity-conferring code, and they together could further reduce the uncertainty.

#### Figure S34

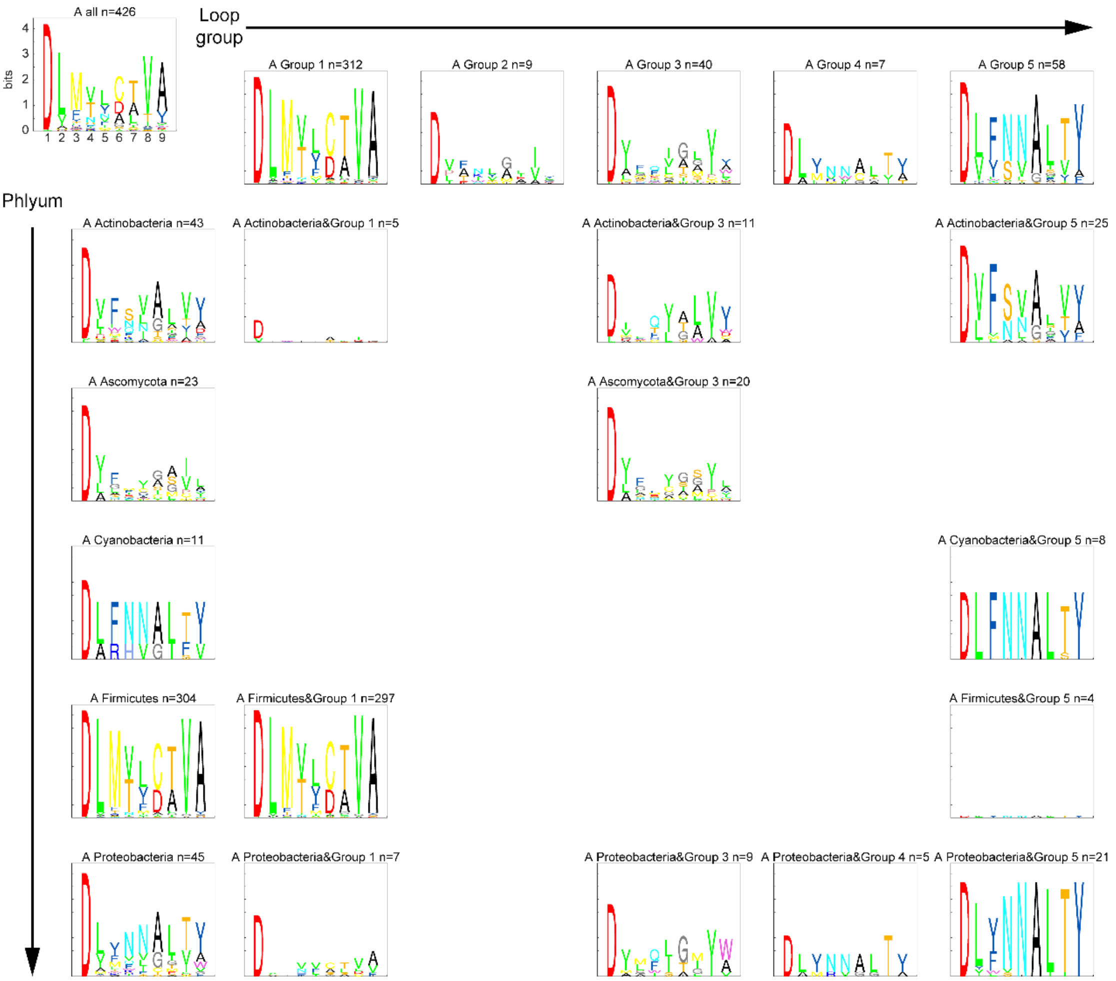


**Figure S34. The sequence logo of the specificity-conferring code for substrate alanine in the dimension of phylum and loop group**

9 of 10 the specificity-conferring code are displayed. The last one is conserved lysine (K) in the A10 motif. It wasn't shown because our A domain sequences only cover A1-A8. Sequence logo will not be plotted, if the number of sequences is less than 3. Substrate abbreviation: A=alanine.

#### Figure S35

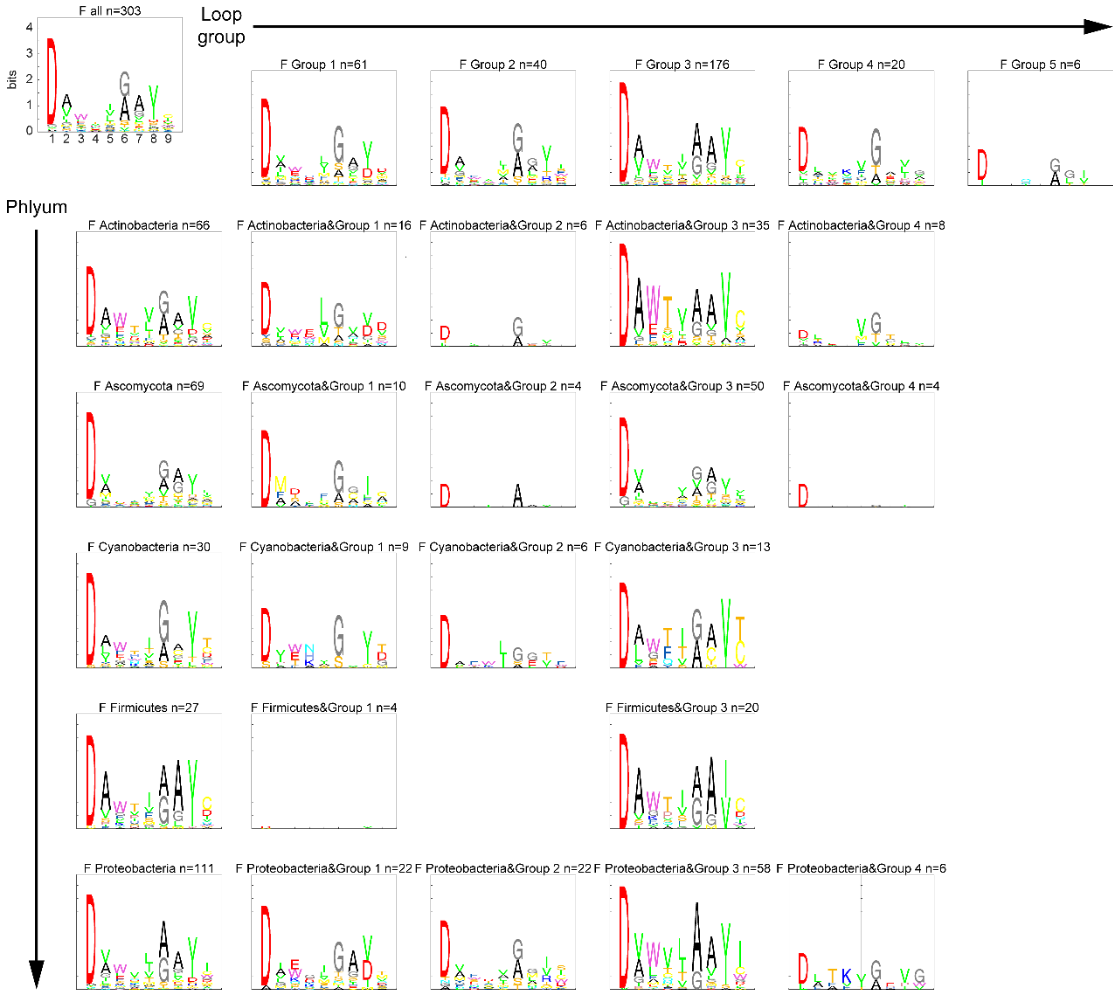


**Figure S35. The sequence logo of the specificity-conferring code for substrate phenylalanine in the dimension of phylum and loop group**

Similar with Figure S34, but for substrate phenylalanine (F).

Figure S36

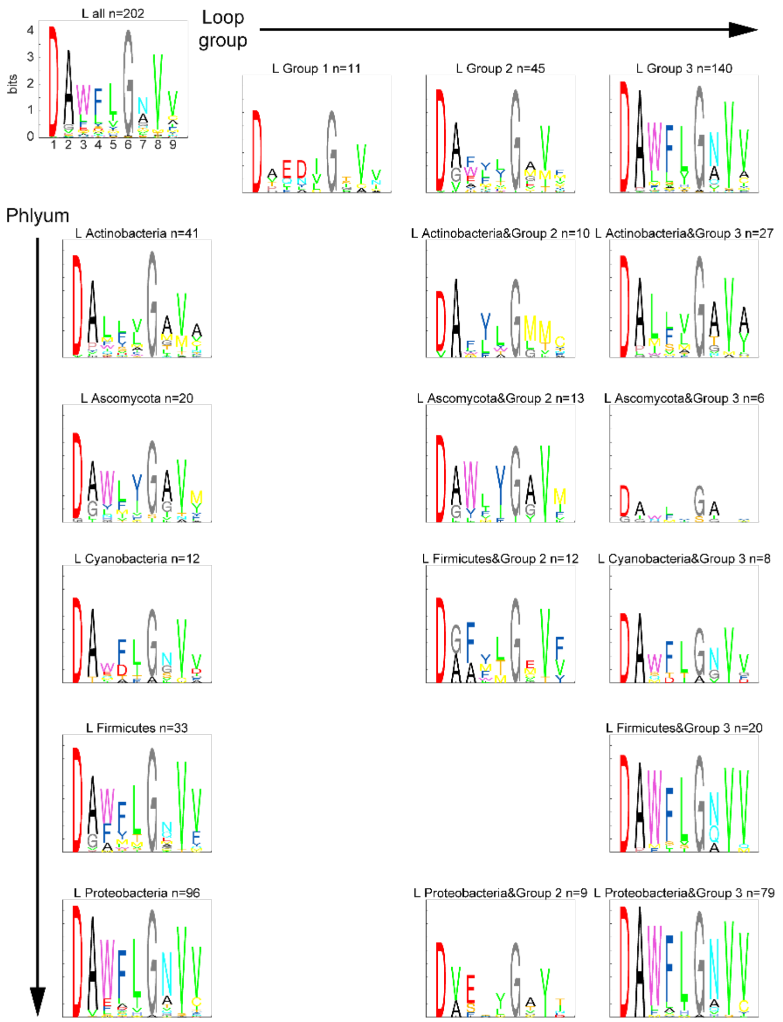


**Figure S36. The sequence logo of the specificity-conferring code for substrate leucine in the dimension of phylum and loop group**

Similar with Figure S34, but for substrate leucine (L).

Figure S37

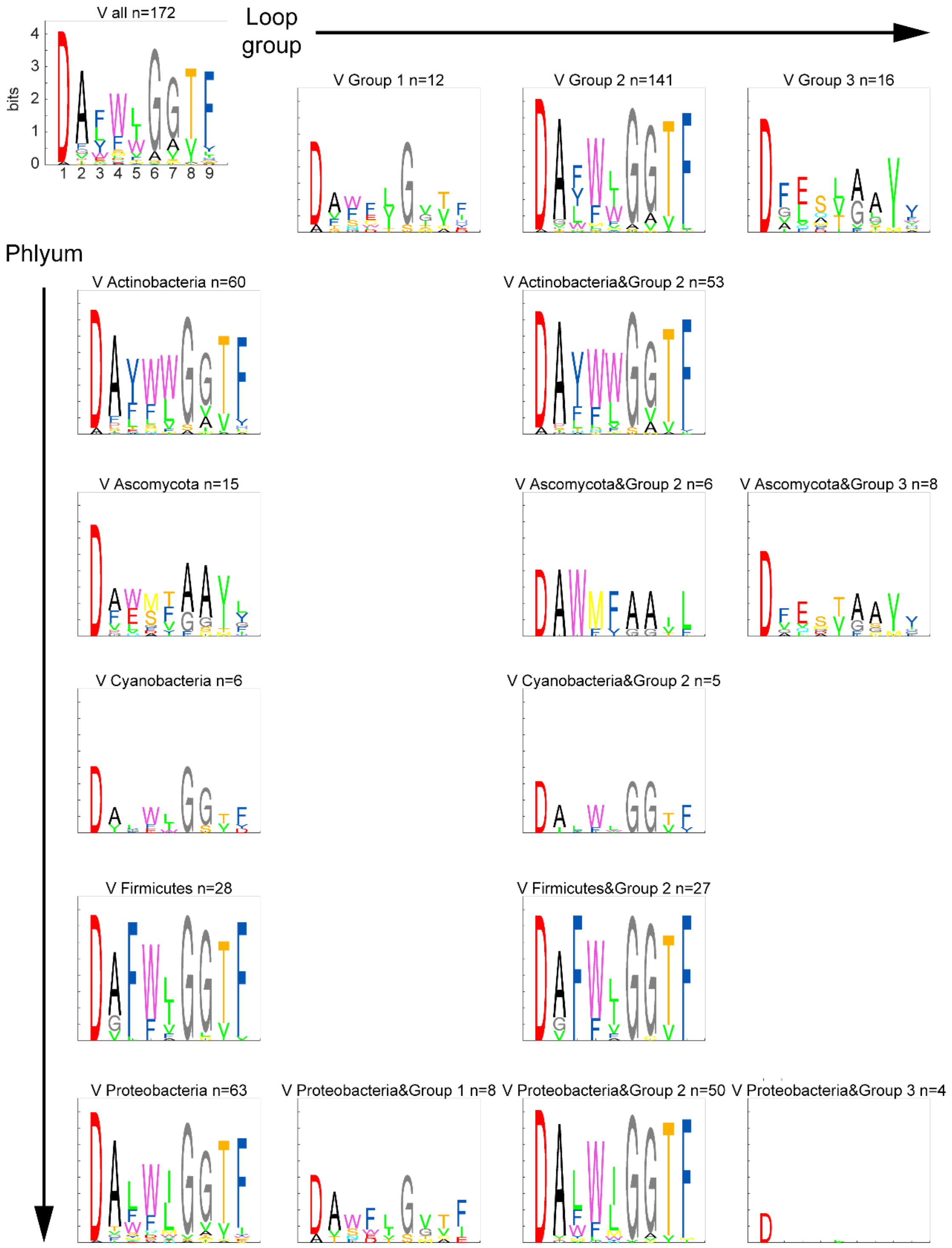


**Figure S37. The sequence logo of the specificity-conferring code for substrate valine in the dimension of phylum and loop group**

Similar with Figure S34, but for substrate valine (V).

#### Figure S38

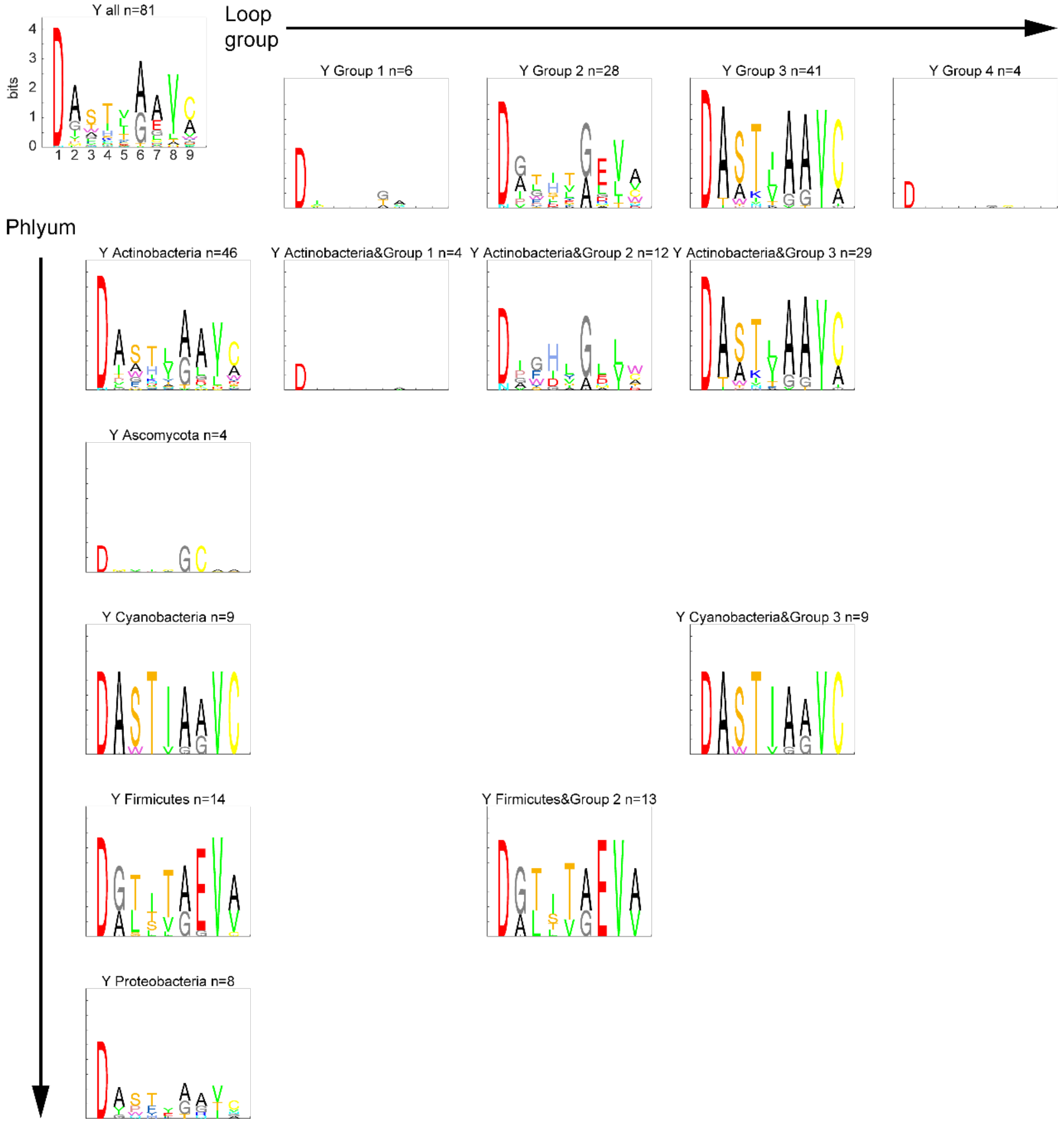


**Figure S38. The sequence logo of the specificity-conferring code for substrate tyrosine in the dimension of phylum and loop group**

Similar with Figure S34, but for substrate tyrosine (Y).

462 Figure S39

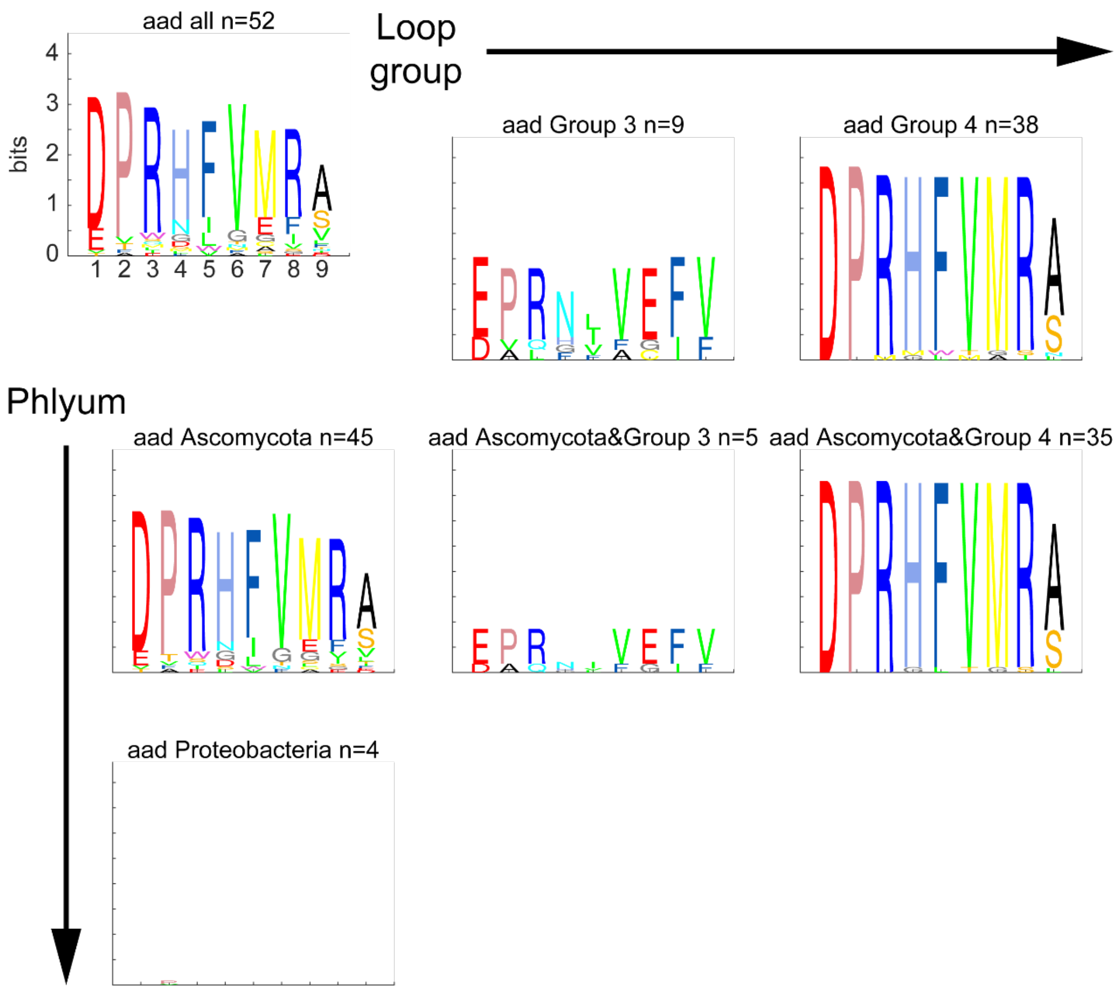


463

464 **Figure S39. The sequence logo of the specificity-conferring code for substrate 2-**
465 **amino-adipic-acidin the dimension of phylum and loop group**
466 Similar with Figure S34, but for substrate 2-amino-adipic-acid (aad).

#### Figure S40

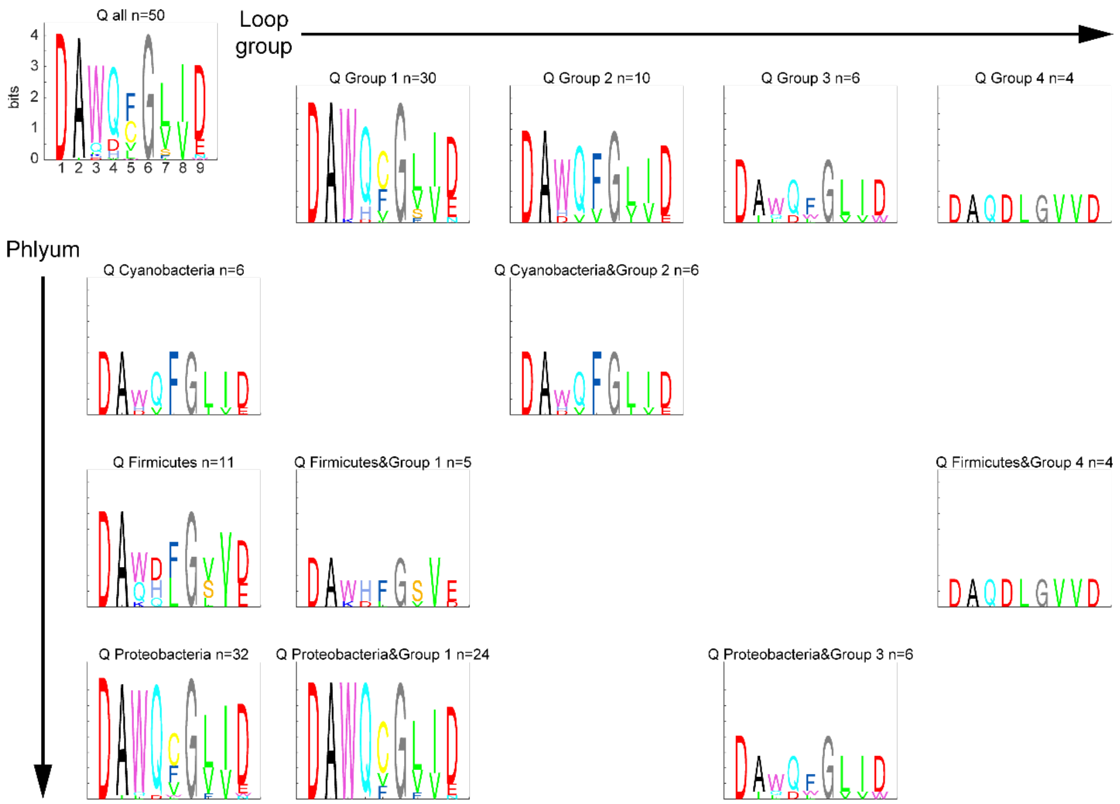


**Figure S40. The sequence logo of the specificity-conferring code for substrate glutamine in the dimension of phylum and loop group**

Similar with Figure S34, but for substrate glutamine (Q).

Figure S41

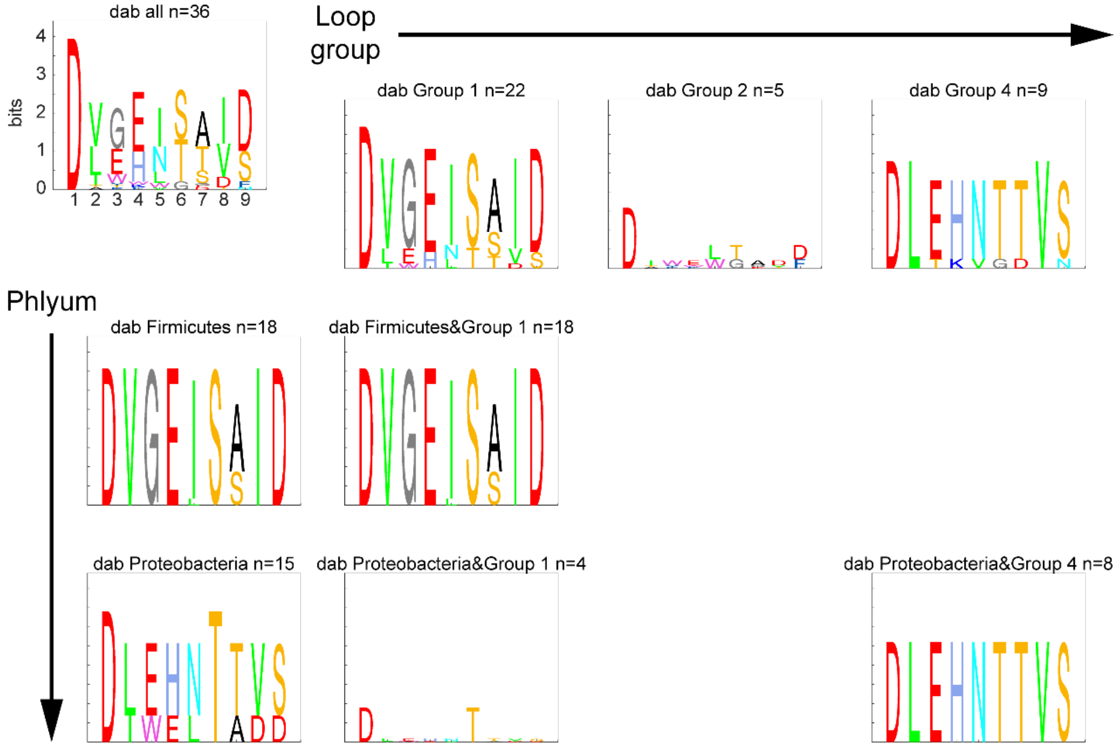


**Figure S41. The sequence logo of the specificity-conferring code for substrate diaminobutyric acid in the dimension of phylum and loop group**

Similar with Figure S34, but for substrate diaminobutyric acid (dab).

Figure S42

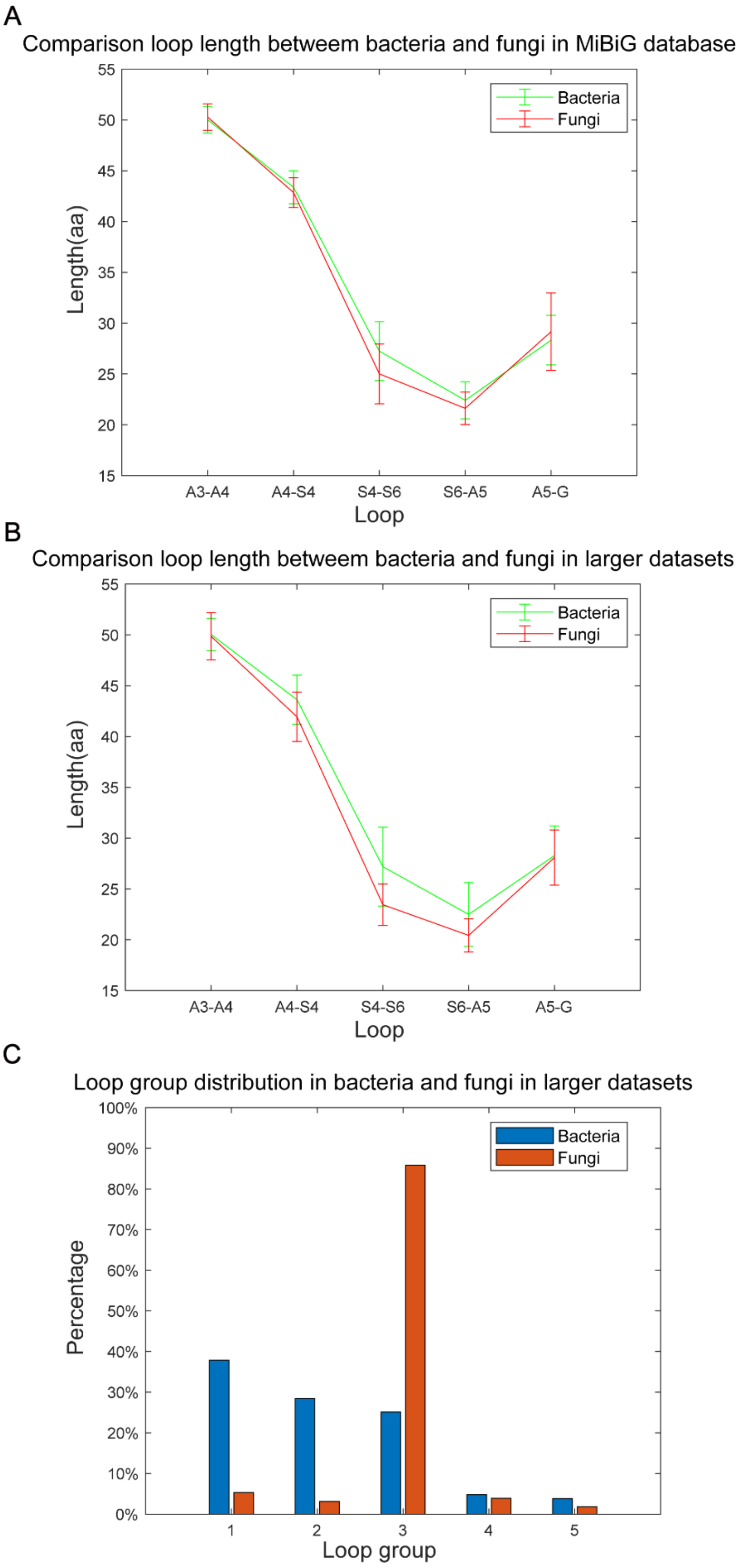


**Figure S42. Loop length and group distributions in bacteria and fungi**

**A.** Comparison loop length between bacteria and fungi in MiBiG database. Numbers of A domains are 2,370 and 215 for bacteria and fungi respectively.

**B.** Comparison loop length between bacteria and fungi in larger dataset. Numbers of A domains are 61,494 and 4,484 for bacteria and fungi respectively.
**C.** Loop group distribution in bacteria and fungi in larger datasets. Numbers of A domains are same with B. Their loop groups are predicted as the closest one loop group in MiBiG database by calculating Euclidean distance. A small amount of data (<5%) is not counted because they are the same distance from multiple loop groups.

#### Figure S43

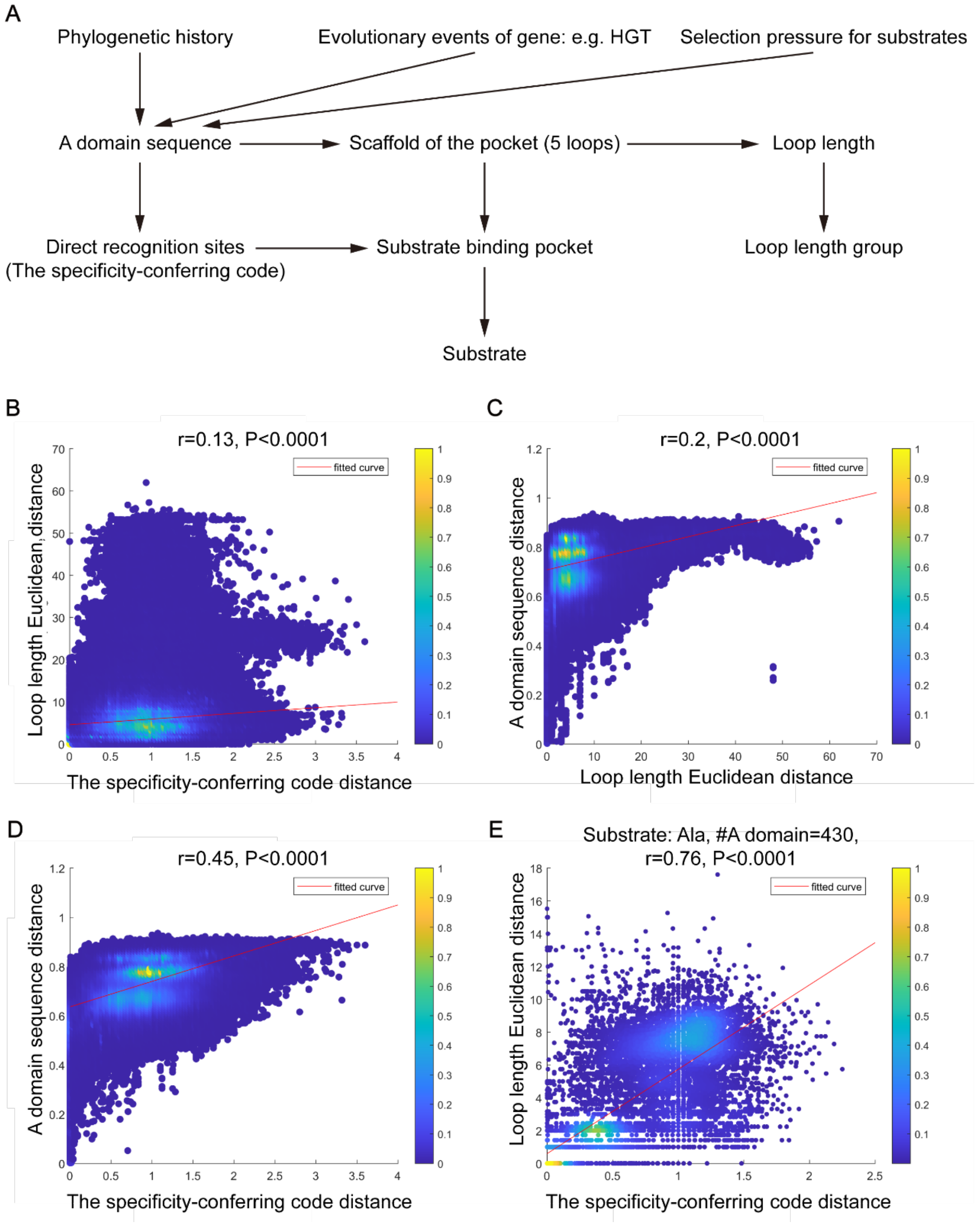


**Figure S43. Causal analysis of A domain substrate specificity**

The specificity-conferring code distance used is alignment-score distance. A domain sequence distance used is p-distance. Loop length distance used is Euclidean distance. r is Pearson correlation coefficient.

**A.** Causal diagram of A domain substrate specificity

**B.** Relationship between the specificity-conferring code distance and Loop length Euclidean distance.

**C.** Relationship between Loop length Euclidean distance and A domain sequence distance.

**D.** Relationship between the specificity-conferring code distance and A domain sequence distance.

**E.** Relationship between the specificity-conferring code distance and Loop length Euclidean distance for A domains with substrate Ala. Similar with B, but for 430 A domains activating Ala as substrate.

Figure S44

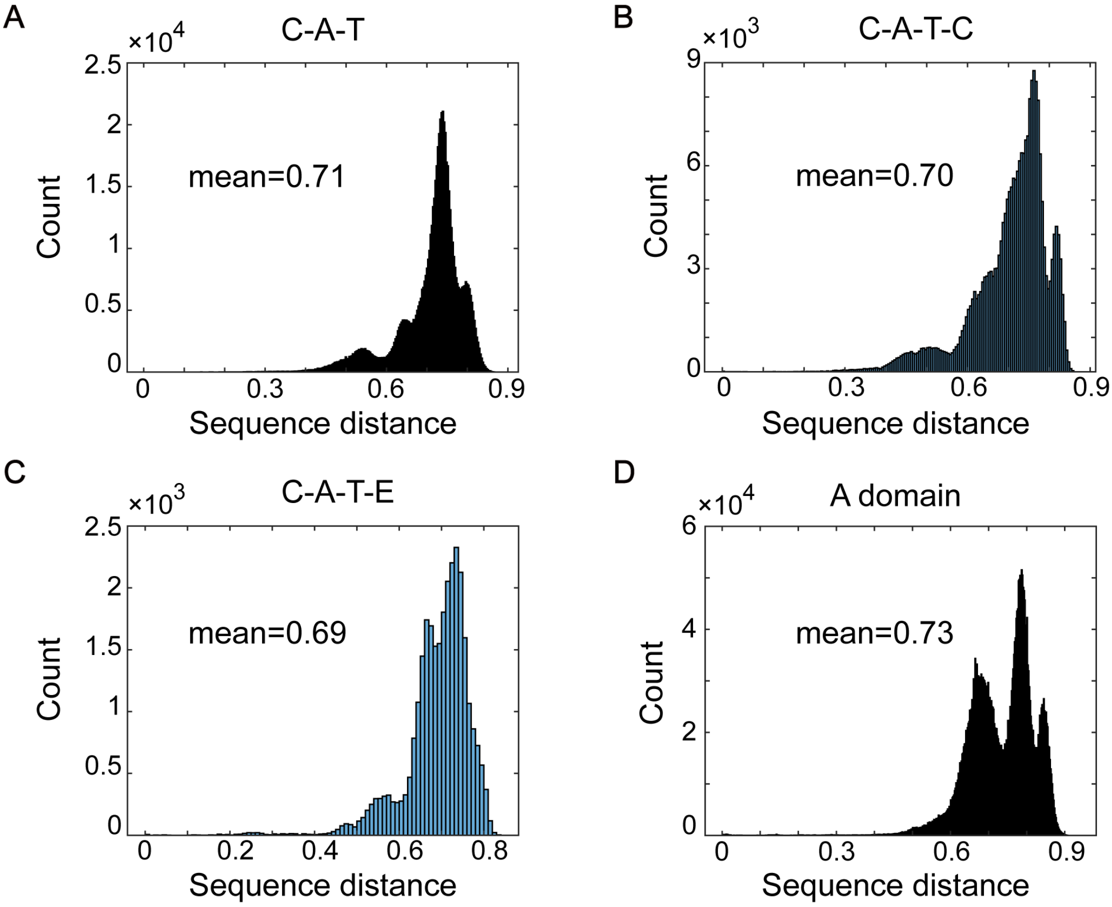


**Figure S44. Protein sequence pairwise distance distribution**

**A.** Pairwise distance distribution of 1,161 C+A+T NRPS sequences. Calculation was based on the p-distance, representing the fraction of amino acid being different after global alignment.

**B.** Same as that in **A**, but for 685 C+A+T+C NRPS sequences.

**C.** Same as that in **A**, but for 245 C+A+T+E NRPS sequences.

**D.** Same as that in **A**, but for 2,636 A domain sequences.

Figure S45

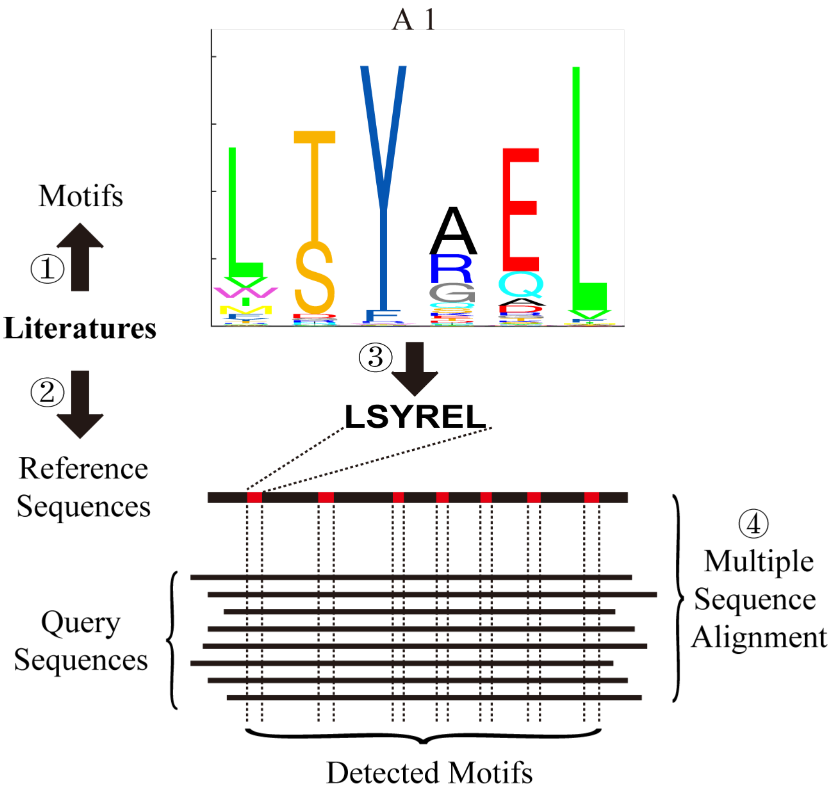


**Figure S45. Workflow of detecting conserved motifs in NRPS domain**

Illustration of the steps in locating known core motifs to query NRPS sequences. First, we curated known core motifs and reference sequences from literatures. Then known motifs on reference sequences were mapped according to previous research, with their locations recorded. Finally, multiple sequence alignment was performed between reference sequences and query sequences. Locations of core motifs in query sequences were inferred by the aligned reference sequences.
