## Supplementary material for "Knowledge-guided data mining on the standardized architecture of NRPS: subtypes, novel motifs, and sequence entanglements": NRPS Motif Finder Matlab version code: Find_NRPS_motif_module_pfam_HRL20221221.m

```
function [result] = Find_NRPS_motif_module_pfam_HRL20221221(seq,Pfam_database_path,reference_motif_path,new_motif,loop_S_code_judge)
%Find_NRPS_motif_module_pfam_HRL20221221 find CATETe domain in the NRPS sequences and mark motifs
%   Note: No blanks in your matltab script path, Pfam_database_path and reference_motif_path
%   Note: 'X' in your sequence will be removed.
%   depend local hmmer3 and a folder with reference sequence and motif information
%   work for C/A/T/E/Te
%   motif number: C:7 A:10(+ 0~2) T:1(+ 0~1) E:7 Te:1  details see reference_motif/reference_motif.xlsx
%   domain id: C:1 A:2 T:3 E:4 Te:5
%   input:seq is the reuslt of fastaread (note: Header of sequence can't
%   strat with blank, that is, ' ')
%       Pfam_database_path is path of Pfam database constructed by hmmpress
%       reference_motif_path is the path of reference motif file folder
%       new_motif: {'Aalpha','G','Talpha'}
%       loop_S_code_judge: 1=calculate loop length, loop group and return
%       loop sequence and S codes.
%   result: a struct
%       Header:the Header in seq
%       domain_list:show the domain compositions of input NRPS sequence
%       motifid_mat:display the motif type in seq_list
%       seq_list:sequence corresponding with motifid_mat
%       index_mat:sequence index for each row in motifid_mat
%       C_subtype_list: the most possible domain subtype for each C domain
%       in sequence
%       C_score: HMM profile alignment score for C subtype. If C_score >=
%       200, prediction is credible. If C_score < 200, it should be careful
%       with the results
%       Raw_C_score_list: HMM profile alignment score for each C subtype
%       C_subtype_str: string of C subtype in C_subtype_list
%       loop length: length of 5 loops:
%           [A3-A4),[A4,S4),[S4,S6),[S6,A5),[A5,G)
%       loop_seq: sequence of 5 loops
%       note: -/*/X in the sequence will be removed, and the index in
%       index_mat will be influenced (shorter than the original)
%       if domain is incomplete, but the length is more than 0.6* full
%       length, we would put NaN in index_mat and '' in seq_list.
%       if index is out of range, we would put '1' in index_mat
%%
if isempty(Pfam_database_path)
    Pfam_database_path='D:/cygwin64/home/74005/Pfam';
%     Pfam_database_path='/storage/disk2/SYZ/program/hmm';
end
if isempty(reference_motif_path)
    reference_motif_path='D:/Shared with me/MATLAB_toolbox/reference_motif';
%     reference_motif_path='/storage/disk2/SM_in_genome/analysis/tools_1025/reference_motif';
end

length_threshold=0.6;% a relaxed thresholds for domain detection by hmmer3
result=[];
for i = 1:length(seq)
    seq(i).Sequence=strrep(seq(i).Sequence,'-','');
    seq(i).Sequence=strrep(seq(i).Sequence,'*','');
    seq(i).Sequence=strrep(seq(i).Sequence,'X','');
end
%import reference sequence and motif information
A_motif_list = readmatrix([reference_motif_path,'/reference_motif.xlsx'],'Sheet',"2VSQ",'Range','L10:M21');
if ~ismember('G',new_motif)
    A_motif_list(7,:)=[];
end
if ~ismember('Aalpha',new_motif)
    A_motif_list(1,:)=[];
end
T_motif_list = readmatrix([reference_motif_path,'/reference_motif.xlsx'],'Sheet',"2VSQ",'Range','L23:M24');
if ~ismember('Talpha',new_motif)
    T_motif_list(1,:)=[];
end
TE_motif_list = readmatrix([reference_motif_path,'/reference_motif.xlsx'],'Sheet',"2VSQ",'Range','L26:M26');
A_seq_range = readmatrix([reference_motif_path,'/reference_motif.xlsx'],'Sheet',"2VSQ",'Range','G10:H10');
T_seq_range = readmatrix([reference_motif_path,'/reference_motif.xlsx'],'Sheet',"2VSQ",'Range','G23:H23');
TE_seq_range = readmatrix([reference_motif_path,'/reference_motif.xlsx'],'Sheet',"2VSQ",'Range','G26:H26');
Module_refer_seq = fastaread([reference_motif_path,'/rcsb_pdb_2VSQ.fasta']);

A_refer_seq = struct;
A_refer_seq.Header=[Module_refer_seq.Header,'|A domain'];
A_refer_seq.Sequence=Module_refer_seq.Sequence(A_seq_range(1):A_seq_range(2));
T_refer_seq = struct;
T_refer_seq.Header=[Module_refer_seq.Header,'|T domain'];
T_refer_seq.Sequence=Module_refer_seq.Sequence(T_seq_range(1):T_seq_range(2));
TE_refer_seq = struct;
TE_refer_seq.Header=[Module_refer_seq.Header,'|TE domain'];
TE_refer_seq.Sequence=Module_refer_seq.Sequence(TE_seq_range(1):TE_seq_range(2));

referOther_seq = [cell(1,1);A_refer_seq;T_refer_seq;cell(1,1);TE_refer_seq];
referOther_motif_list={cell(1,1);A_motif_list;T_motif_list;cell(1,1);TE_motif_list};

referCE_seq_raw=fastaread([reference_motif_path,'/reference.fasta']);
referCE_motif_list_raw=readmatrix([reference_motif_path,'/reference_motif.xlsx'],'Sheet','Sheet1','Range','C2:D134');
referCE_motif_list=cell(length(referCE_seq_raw),1);
referCE_seq=cell(length(referCE_seq_raw),1);
referCE_dtype_str=readcell([reference_motif_path,'/reference_motif.xlsx'],'Sheet','gap','Range','A5:A23');
for i = 1:length(referCE_seq_raw)
    referCE_seq{i}=referCE_seq_raw(i);
    referCE_motif_list{i}=referCE_motif_list_raw(1+(i-1)*7:i*7,:);
end
if ~exist('referCE_HMM_struct','var')
    load([reference_motif_path,'/HMM/hmm_struct.mat']);
end

loop_group_ref_seq=fastaread([reference_motif_path,'/5loop_seq.fasta']);
seq_1AMU=fastaread([reference_motif_path,'/rcsb_pdb_1AMU.fasta']);
S_code_site=readmatrix([reference_motif_path,'/reference_motif.xlsx'],'Sheet',"2VSQ",'Range','B35:B43');
loop_range_list = readmatrix([reference_motif_path,'/reference_motif.xlsx'],'Sheet',"2VSQ",'Range','L28:M33');
%% hmm scan by pfam
% loc_pwd=pwd;
tmp_folder='tmp_24601';
if exist('tmp_24601','dir')==7
    delete(['./',tmp_folder,'/*.*']);
else
    mkdir tmp_24601 % bulid a temporary folder
end

fastawrite('./tmp_24601/input_seq.fasta',seq); % prepare fasta file for hmmer3
command=['hmmscan -o ./tmp_24601/out.txt --domtblout ./tmp_24601/out1.txt --notextw --noali -E 1e-5 --domE 1e-5 ',Pfam_database_path,'/Pfam-A.hmm ./tmp_24601/input_seq.fasta'];
% command=strrep(command,'/','\');
system(command);
fid = fopen('./tmp_24601/out1.txt','rt');%extract domain detction result from hmmer3 output file
hmm_r=[];% hmmer3 result
while true
    thisline = fgetl(fid);
    if ischar(thisline)
        if ~startsWith(thisline,'#')
            newthisline = split(thisline)';
            if length(newthisline)>23
            newthisline(24:end)=[];
            end
            hmm_r=[hmm_r;newthisline];
        end
    else
      break; 
    end
end
fclose(fid);
%%
for ii = 1:length(seq) % for each sequence
    loc_header = seq(ii).Header;
    loc_header = split(loc_header,' ');
    loc_header = loc_header{1};
    input_seq = seq(ii).Sequence;
    region = find(strcmp(hmm_r(:,4),loc_header));
    % process pfam result
    start_list=[];
    end_list=[];
    domain_list=[];
    pesu_region=[];
    pesu_domain_list=[];
    for k = 1:length(region)
        i = region(k);
%         if ((str2double(hmm_r(i,21))-str2double(hmm_r(i,20))+1)>length_threshold*str2double(hmm_r(i,3)))||(strcmp(hmm_r{i,1},'Thioesterase')&&(str2double(hmm_r(i,21))-str2double(hmm_r(i,20))+1)>0.5*str2double(hmm_r(i,3))) % the length of matching shouldn't be too short (>length_threshold*domain_length)
        if ((str2double(hmm_r(i,17))-str2double(hmm_r(i,16))+1)>length_threshold*str2double(hmm_r(i,3)))||(strcmp(hmm_r{i,1},'Thioesterase')&&(str2double(hmm_r(i,17))-str2double(hmm_r(i,16))+1)>0.5*str2double(hmm_r(i,3))) % the length of matching shouldn't be too short (>length_threshold*domain_length)
            % a more relaxed threshold for Te domain, here is 0.5
            if strcmp(hmm_r{i,1},'Condensation')
                domain_list=[domain_list;1];
            elseif strcmp(hmm_r{i,1},'AMP-binding')
                domain_list=[domain_list;2];
            elseif strcmp(hmm_r{i,1},'PP-binding')
                domain_list=[domain_list;3];
            elseif strcmp(hmm_r{i,1},'Thioesterase')
                domain_list=[domain_list;5];
            else
                continue % if not CATTe, ignore and continue
            end
            start_list=[start_list;str2double(hmm_r(i,20))];
            end_list=[end_list;str2double(hmm_r(i,21))];
        else
            pesu_region=[pesu_region;region(k)];
            if strcmp(hmm_r{i,1},'Condensation')
                pesu_domain_list=[pesu_domain_list;1];
            elseif strcmp(hmm_r{i,1},'AMP-binding')
                pesu_domain_list=[pesu_domain_list;2];
            elseif strcmp(hmm_r{i,1},'PP-binding')
                pesu_domain_list=[pesu_domain_list;3];
            elseif strcmp(hmm_r{i,1},'Thioesterase')
                pesu_domain_list=[pesu_domain_list;5];
            else
                pesu_region(end)=[];
                continue
            end
        end
    end
    % process the problem because hmmer3 sometime will spilt one domain to
    % two same type domains.
    pesu_region_type=hmm_r(pesu_region,1);
    [uni_pesu_region_type,~,ic]=unique(pesu_region_type);
    for i = 1:length(uni_pesu_region_type)
        loc_index = find(ic==i);
        if length(loc_index)>1
            remove_list=[];
            domain_length=str2double(hmm_r{pesu_region(loc_index(1)),3});
            for j = 1:length(loc_index)
                if ~ismember(j,remove_list)
                   for k = 1:length(loc_index)
                       if ~ismember(k,[remove_list;j])%j~=k
                           start1=str2double(hmm_r(pesu_region(loc_index(j)),20));
                           start2=str2double(hmm_r(pesu_region(loc_index(k)),20));
                           end1=str2double(hmm_r(pesu_region(loc_index(j)),21));
                           end2=str2double(hmm_r(pesu_region(loc_index(k)),21));
                           if (start1<=start2&&start2<=end1)||(start1<=end2&&end2<=end1)||(end2<=start1&&((start1-end2)<0.5*domain_length))||(start2>=end1&&((start2-end1)<0.5*domain_length))
                               new_start=min([start1,start2]);
                               new_end=max([end1,end2]);
                           else
                               remove_list=[remove_list;k;j]; % no overlaping, also remove it
                               continue
                           end
                           if (~any(start_list>new_start&start_list<new_end))&&(~any(end_list>new_start&end_list<new_end))&&((new_end-new_start+1)>length_threshold*domain_length)
                               domain_list=[domain_list;pesu_domain_list(loc_index(j))];
                               start_list=[start_list;new_start];
                               end_list=[end_list;new_end];
                               remove_list=[remove_list;k;j];
                               break;
                           end
                       end
                   end
                end
            end
        end
    end
    [start_list_sorted,I]=sort(start_list);
    end_list_sorted=end_list(I);
    domain_list_sorted=domain_list(I);
    %% find motif by reference sequence
    motif_list=[];
    C_score=[];
    C_subtype_list=[];
    Raw_C_score_list=[];
    loop_length=[];
    loop_seq=[];
    S_code=[];
    for i = 1:length(domain_list_sorted)
        if domain_list_sorted(i)==1 %if the domain type detected is C, we must check whether it is a E domain
            if i == length(domain_list_sorted)
               loc_seq=input_seq(start_list_sorted(i):end);
            else
               loc_seq=input_seq(start_list_sorted(i):start_list_sorted(i+1));
            end
            loc_score_matrix=zeros(length(referCE_HMM_struct),1);
            for j = 1:length(referCE_HMM_struct)
                loc_score_matrix(j)=hmmprofalign(referCE_HMM_struct{j},loc_seq);
            end
            [loc_C_score,I]=max(loc_score_matrix);
            if I == length(referCE_dtype_str) % E domain is the last one in referCE
                domain_list_sorted(i)=4;
            else
                C_subtype_list=[C_subtype_list;I];
                C_score=[C_score;loc_C_score];
                Raw_C_score_list=[Raw_C_score_list;loc_score_matrix'];
            end
            loc_header={'query';referCE_seq{I}.Header};
            loc_sequence={loc_seq;referCE_seq{I}.Sequence};
            fastawrite(['./tmp_24601/input',num2str(ii),'_',num2str(i),'_seq.fasta'],loc_header,loc_sequence);
            command = ['clustalo -i ./tmp_24601/input',num2str(ii),'_',num2str(i),'_seq.fasta -o ./tmp_24601/input',num2str(ii),'_',num2str(i),'_seq_MSA.fasta --outfmt fasta --output-order input-order'];
            system(command);
            loc_seq_MSA=fastaread(['./tmp_24601/input',num2str(ii),'_',num2str(i),'_seq_MSA.fasta']);
            ref_start=strfind(referCE_seq{I}.Sequence,strrep(loc_seq_MSA(2).Sequence,'-',''));
            loc_motif_list=[];
            gap_list = strfind(loc_seq_MSA(1).Sequence,'-');
            for j = 1:size(referCE_motif_list{I},1)
                loc_start=Codetransform_HRL(loc_seq_MSA(2).Sequence,referCE_motif_list{I}(j,1)-ref_start+1);
                loc_end=Codetransform_HRL(loc_seq_MSA(2).Sequence,referCE_motif_list{I}(j,2)-ref_start+1);
                loc_start2=loc_start-sum(gap_list<loc_start)+start_list_sorted(i)-1;
                loc_end2=loc_end-sum(gap_list<loc_end)+start_list_sorted(i)-1;
                if i == length(domain_list_sorted)
                    compared_end = length(input_seq);
                else
                    compared_end = start_list_sorted(i+1);
                end
                if loc_start <=0 ||loc_start2==loc_end2 || loc_end2>= compared_end
                    loc_motif_list=[loc_motif_list;nan,nan];
                else
                    loc_motif_list=[loc_motif_list;loc_start2,loc_end2];
                end
            end
        else
            if i == 1
                loc_seq_start=1;
            else
                if domain_list_sorted(i)==2||domain_list_sorted(i)==3
                    loc_seq_start=end_list_sorted(i-1);
                elseif domain_list_sorted(i)==5
                    loc_seq_start=start_list_sorted(i);
                end
            end
            if i == length(domain_list_sorted)
                loc_seq_end = length(input_seq);
            else
                if domain_list_sorted(i)==2
                    loc_seq_end = start_list_sorted(i+1);
                elseif domain_list_sorted(i)==3||domain_list_sorted(i)==5
                    loc_seq_end = end_list_sorted(i);
                end
            end
            loc_seq=input_seq(loc_seq_start:loc_seq_end);
            loc_header={'query';referOther_seq{domain_list_sorted(i)}.Header};
            loc_sequence={loc_seq;referOther_seq{domain_list_sorted(i)}.Sequence};
            fastawrite(['./tmp_24601/input',num2str(ii),'_',num2str(i),'_seq.fasta'],loc_header,loc_sequence);
            command = ['clustalo -i ./tmp_24601/input',num2str(ii),'_',num2str(i),'_seq.fasta -o ./tmp_24601/input',num2str(ii),'_',num2str(i),'_seq_MSA.fasta --outfmt fasta --output-order input-order'];
            system(command);
            loc_seq_MSA=fastaread(['./tmp_24601/input',num2str(ii),'_',num2str(i),'_seq_MSA.fasta']);
            ref_start=strfind(referOther_seq{domain_list_sorted(i)}.Sequence,strrep(loc_seq_MSA(2).Sequence,'-',''));
            seq_start=strfind(loc_seq,strrep(loc_seq_MSA(1).Sequence,'-',''));
            loc_motif_list=[];
            gap_list = strfind(loc_seq_MSA(1).Sequence,'-');
            for j = 1:size(referOther_motif_list{domain_list_sorted(i)},1)
                loc_start=Codetransform_HRL(loc_seq_MSA(2).Sequence,referOther_motif_list{domain_list_sorted(i)}(j,1)-ref_start+1);
                loc_end=Codetransform_HRL(loc_seq_MSA(2).Sequence,referOther_motif_list{domain_list_sorted(i)}(j,2)-ref_start+1);
                loc_start2=loc_start-sum(gap_list<loc_start)+loc_seq_start+seq_start-2;
                loc_end2=loc_end-sum(gap_list<loc_end)+loc_seq_start+seq_start-2;
                if i == length(domain_list_sorted)
                    compared_end = length(input_seq);
                else
                    compared_end = start_list_sorted(i+1);
                end
                if loc_start <=0 || loc_start2==loc_end2 || loc_end2>= compared_end
                    loc_motif_list=[loc_motif_list;nan,nan];
                else
                    loc_motif_list=[loc_motif_list;loc_start2,loc_end2];
                end
            end
            % calculate loop length
            if loop_S_code_judge
                loc_loop_length=[];
                loc_loop_seq=[];
                loc_S_code=[];
                if domain_list_sorted(i)==2
                    loc_seq_struct.Sequence=loc_seq;
                    loc_seq_struct.Header='input';
                    loc_loop_seq_struct=[loc_seq_struct;A_refer_seq;seq_1AMU;loop_group_ref_seq];
                    fastawrite(['./tmp_24601/input',num2str(ii),'_',num2str(i),'_seq1.fasta'],loc_loop_seq_struct);
                    command = ['clustalo -i ./tmp_24601/input',num2str(ii),'_',num2str(i),'_seq1.fasta -o ./tmp_24601/input',num2str(ii),'_',num2str(i),'_seq1_MSA.fasta --outfmt fasta --output-order input-order'];
                    system(command);
                    loc_loop_seq_MSA=fastaread(['./tmp_24601/input',num2str(ii),'_',num2str(i),'_seq1_MSA.fasta']);
                    loc_loop_range=zeros(size(loop_range_list,1),1);
                    for j = 1:size(loop_range_list,1)
                        loc_loop_range(j) = Codetransform_HRL(loc_loop_seq_MSA(2).Sequence,loop_range_list(j,1));
                    end
                    loc_loop_length=zeros(1,size(loop_range_list,1)-1);
                    loc_loop_seq=cell(1,size(loop_range_list,1)-1);
                    for j = 1:size(loop_range_list,1)-1
                        loc_seq1=strrep(loc_loop_seq_MSA(1).Sequence(loc_loop_range(j):loc_loop_range(j+1)-1),'-','');
                        loc_loop_length(j)=length(loc_seq1);
                        loc_loop_seq{j}=loc_seq1;
                    end
                    loc_index1=zeros(1,length(S_code_site));
                    for j = 1:length(S_code_site)
                        loc_index1(j)=Codetransform_HRL(loc_loop_seq_MSA(3).Sequence,S_code_site(j));
                    end
                    loc_S_code={loc_loop_seq_MSA(1).Sequence(loc_index1)};
                end
                loop_length=[loop_length;loc_loop_length];
                loop_seq=[loop_seq;loc_loop_seq];
                S_code=[S_code;loc_S_code];
            end
        end
        motif_list=[motif_list;{loc_motif_list}];
    end
    %%
    % obtain sequence from index
    motifid_mat=[];
    seq_list=[];
    index_mat=[];
    for i = 1:length(domain_list_sorted)
        loc_motifid_mat=[domain_list_sorted(i)*ones(2*size(motif_list{i},1),1),linspace(0.5,size(motif_list{i},1),2*size(motif_list{i},1))'];
        if ~isnan(motif_list{i}(1,1))
            start_is_nan=0;
            if i == 1
                loc_seq={input_seq(1:motif_list{i}(1,1)-1)};
                loc_index_mat=[1,motif_list{i}(1,1)-1];
            else
                if ~isnan(motif_list{i-1}(end,2))
                    loc_seq={input_seq(motif_list{i-1}(end,2)+1:motif_list{i}(1,1)-1)};
                    loc_index_mat=[motif_list{i-1}(end,2)+1,motif_list{i}(1,1)-1];
                else % if last domain index is NaN, take the last is not NaN index
                    last_nan = motif_list{i-1}(end,2);
                    last_end = size(motif_list{i-1},1);
                    while isnan(last_nan)
                        last_end=last_end-1;
                        last_nan = motif_list{i-1}(last_end,2);
                    end
                    loc_seq={input_seq(motif_list{i-1}(last_end,2)+1:motif_list{i}(1,1)-1)};
                    loc_index_mat=[motif_list{i-1}(last_end,2)+1,motif_list{i}(1,1)-1];
                end
            end
            loc_seq=[loc_seq;{input_seq(motif_list{i}(1,1):motif_list{i}(1,2))}];
            loc_index_mat=[loc_index_mat;motif_list{i}(1,1),motif_list{i}(1,2)];
        else
            start_is_nan=1;
            loc_seq=[{''};{''}];
            loc_index_mat=[nan,nan;nan,nan];
        end
        if size(motif_list{i},1) > 1
            for j = 2:size(motif_list{i},1)
                if ~isnan(motif_list{i}(j,1))
                    before_all_nan=0;
                    if ~isnan(motif_list{i}(j-1,1))
                        inter_start=motif_list{i}(j-1,2)+1;
                    else
                        before_motif_list=motif_list{i}(1:j-1,2);
                        if ~all(isnan(before_motif_list))
                            inter_start=motif_list{i}(find(~isnan(before_motif_list),1,'last'),2)+1;% last non-nan end position
                        else
                            before_all_nan=1;
                        end
                    end
                    inter_end=motif_list{i}(j,1)-1;
                    motif_start=motif_list{i}(j,1);
                    motif_end=motif_list{i}(j,2);
                    if before_all_nan
                         loc_seq=[loc_seq;{''}];
                         loc_index_mat=[loc_index_mat;[nan,nan]];
                    else
                        loc_seq=[loc_seq;{input_seq(inter_start:inter_end)}];%inter
                        loc_index_mat=[loc_index_mat;inter_start,inter_end];
                    end
                    loc_seq=[loc_seq;{input_seq(motif_start:motif_end)}];%motif
                    loc_index_mat=[loc_index_mat;motif_start,motif_end];
                else% this is a nan
                    loc_seq=[loc_seq;{''};{''}];%inter
                    loc_index_mat=[loc_index_mat;nan,nan;nan,nan];
                end
            end
        else
            j = 1;
        end
        if i == length(domain_list_sorted)
            loc_motifid_mat=[loc_motifid_mat;loc_motifid_mat(end,1),loc_motifid_mat(end,2)+0.5];
            if ~isnan(motif_list{i}(j,1))
                last_start=motif_list{i}(j,2)+1;
            else
                before_motif_list=motif_list{i}(1:j,2);
                last_start=motif_list{i}(find(~isnan(before_motif_list),1,'last'),2)+1;% last non-nan end position
            end
            loc_seq=[loc_seq;{input_seq(last_start:end)}];
            last_end=length(input_seq);
            if last_start>=length(input_seq)
                last_start=length(input_seq);
                last_end=1;
            else
                if last_end>=length(input_seq)
                    last_end=length(input_seq);
                end
            end
            loc_index_mat=[loc_index_mat;last_start,last_end];
        end
        if start_is_nan&&~isempty(index_mat)
            last_non_nan=find(~isnan(index_mat(:,2)),1,'last');
            first_non_nan=find(~isnan(loc_index_mat(:,2)),1);
            loc_index_mat(1,:)=[index_mat(last_non_nan,2)+1,loc_index_mat(first_non_nan,1)-1];% put on the first one
            loc_seq{1}=input_seq(loc_index_mat(1,1):loc_index_mat(1,2));
        end
        motifid_mat=[motifid_mat;loc_motifid_mat];
        seq_list=[seq_list;loc_seq];
        index_mat=[index_mat;loc_index_mat];
    end
    loc_result.Header=seq(ii).Header;
    loc_result.domain_list=domain_list_sorted;
    loc_result.motifid_mat=motifid_mat;
    loc_result.seq_list=seq_list;
    loc_result.index_mat=index_mat;
    loc_result.C_subtype_list=C_subtype_list;
    loc_result.C_score=C_score;
    loc_result.Raw_C_score_list=Raw_C_score_list(:,1:end-1);
    loc_result.C_subtype_str=referCE_dtype_str(1:end-1);
    if loop_S_code_judge
        loc_result.loop_length=loop_length;
        loc_result.loop_seq=loop_seq;
        loc_result.S_code=S_code;
        loop_group = Loop_length2group_HRL(reference_motif_path,loop_length);
        loc_result.loop_group=loop_group;
    end
    result=[result;loc_result];
end
rmdir('tmp_24601', 's')
end
```
