## Supplementary material for "Knowledge-guided data mining on the standardized architecture of NRPS: subtypes, novel motifs, and sequence entanglements": NRPS Motif Finder online version code (Python): Find_motif_HRL_0207.py

```
from cProfile import run
from operator import itemgetter
import subprocess
import sys
import getopt
import json5
import pandas as pd
import numpy as np
import scipy.io as sio
import os
import re

from Bio import SeqIO
from Bio.Seq import Seq
from Bio.SeqRecord import SeqRecord
from Bio.SeqFeature import SeqFeature, FeatureLocation
from Bio.Align import substitution_matrices
from Bio import Align

# 用于写变量到字符串中
import sys

import warnings
warnings.filterwarnings("ignore")


class safesub(dict):
	"""防止key找不到"""

	def __missing__(self, key):
		return '{' + key + '}'


outputdic = {}
argv = sys.argv[1:]

inputfile = outputfile = processid = ''

try:
	opts, args = getopt.getopt(argv, "h i o p G A T", [
		"i=", "o=", "p=", "G=", "A=", "T="])
except getopt.GetoptError:
	outputdic["Result data"] = False
	outputdic["Reason data"] = "Get opt error"
	outputjson = json5.dumps(outputdic)
	print(outputjson)
	sys.exit()
for opt, arg in opts:
	if opt == '-h':
		print('Find_motif_HRL.py -i <inputfile> -o <outputfile> -p <processid> -G <1 or 0> -A <1 or 0> -T <1 or 0>')
		sys.exit()
	elif opt in ("--i"):
		inputfile = arg
	elif opt in ("--o"):
		outputfile = arg
	elif opt in ("--p"):
		processid = arg
	elif opt in ("--G"):
		G_judge = arg
	elif opt in ("--A"):
		A_judge = arg
	elif opt in ("--T"):
		T_judge = arg

# source_path = "/Users/chenhaoran/code/rawpool/biology/"

# source_path = "/Users/chenhaoran/code/rawpool/biology/python/NRPSMotif/"

run_default_path = "/data/services/biology/python/NRPSMotif/"
source_path = ""

os.chdir(run_default_path)

hmm_out_dom_path = "./Temp/hmm_out_" + \
	processid + ".dom"
hmm_out_txt_path = "./Temp/hmm_out_" + \
	processid + ".txt"
temp_aln_fasta_path = "./Temp/temp_aln_" + \
	processid + ".fasta"
temp_fasta_path = "./Temp/temp_" + \
	processid + ".fasta"
open(hmm_out_dom_path, "w")
open(hmm_out_txt_path, "w")
open(temp_aln_fasta_path, "w")
open(temp_fasta_path, "w")

seqrecords = [seq for seq in SeqIO.parse(source_path + inputfile, "fasta")]


def sub(text):
	return text.format_map(safesub(sys._getframe(1).f_locals))


def Codetransform_HRL(align_seq, code):
	gap_site = [i+1 for i, char in enumerate(align_seq) if char == '-']
	try:
		code_align = np.zeros((len(code), 1))
	except TypeError:
		code = np.array([code])
		code_align = np.zeros((len(code), 1))
	for i in range(len(code)):
		new = code[i] + len([a for a in gap_site if a <= code[i]])
		old = code[i]
		while len([a for a in gap_site if a <= old]) != len([a for a in gap_site if a <= new]):
			old = new
			new = code[i] + sum(gap_site <= new)
		code_align[i] = new
	return(code_align)


def find_motif(seq, ref_seq, motif_list, list_sorted, i_, seqstart):
	query = SeqRecord(seq, id='query')
	refrence = SeqRecord(Seq(ref_seq), id='targrt')
	SeqIO.write([query, refrence], temp_fasta_path, "fasta")
	subprocess.run(
		sub('./clustal -i '+temp_fasta_path+' -o '+temp_aln_fasta_path+' --force'), shell=True)
	loc_seq_MSA = [seq for seq in SeqIO.parse(
		sub(temp_aln_fasta_path), "fasta")]
	# ref_start = min([i for i, char in enumerate(loc_seq_MSA[1].seq) if char != '-'])
	loc_motif_list = []
	gap_list = [i+1 for i,
				char in enumerate(loc_seq_MSA[0].seq) if char == '-']

	for j in range(len(motif_list)):
		# 注意参考是matlab坐标
		loc_start = Codetransform_HRL(loc_seq_MSA[1].seq, motif_list[j, 0])
		loc_end = Codetransform_HRL(loc_seq_MSA[1].seq, motif_list[j, 1])
		loc_start2 = loc_start - \
			len([a for a in gap_list if a < loc_start]) + seqstart
		loc_end2 = loc_end - \
			len([a for a in gap_list if a < loc_end]) + seqstart
		if i_ == len(list_sorted)-1:
			compared_end = len(seq)
		else:
			compared_end = list_sorted.loc[i_+1]['start']
		if (int(loc_start) <= -1) or (int(loc_start2) == int(loc_end2)) or (int(loc_end2) >= compared_end):
			loc_motif_list.append([np.array([np.nan]), np.array([np.nan])])
		else:
			loc_motif_list.append([loc_start2, loc_end2])
	return(loc_motif_list)

# loop_length 2-D arrary


def Loop_length2group_HRL(loop_length, mat_file):
	if np.size(loop_length) == 0:
		return([])
	loop_length_ref = mat_file['loop_length_ref']
	loop_group_ref = mat_file['loop_group_ref']
	loop_group = np.zeros(len(loop_length)) + 50

	for i in range(len(loop_length)):
		score = np.zeros(len(loop_length_ref))
		for j in range(len(loop_length_ref)):
			score[j] = np.linalg.norm(loop_length[i]-loop_length_ref[j])
			loop_group[i] = loop_group_ref[score == min(score)][0]
	return(loop_group)


NRPS_domains = ['Condensation', 'AMP-binding', 'PP-binding', 'Thioesterase']
AA_leter = 'ACDEFGHIKLMNPQRSTVWYX'

mat_file = sio.loadmat(run_default_path + 'matlab230206.mat')
A_refer_seq = mat_file['A_refer_seq']['Sequence'][0][0][0]
A_motif_list = mat_file['A_motif_list']
A_motif_name = ['A_alpha', 1, 2, 3, 4, 5, 6, 'G', 7, 8, 9, 10]
T_refer_seq = mat_file['T_refer_seq']['Sequence'][0][0][0]
T_motif_list = mat_file['T_motif_list']
T_motif_name = ['T_alpha', 1]

TE_refer_seq = mat_file['TE_refer_seq']['Sequence'][0][0][0]
TE_motif_list = mat_file['TE_motif_list']
TE_motif_name = list(range(1, len(TE_motif_list)+1))

# 2023/2/4 更新了C的subtype
referCE_dtype_str = ['LCL','SgcC5','DCL','Starter','Cyc','Dual','Cglyc','Hybrid','modAA','CT','CT-DCL','CT-LCL','FUM14','It','I','bL','PS','X','E']
referCE_motif_list = [mat_file['referCE_motif_list'][i][0] for i in range(len(mat_file['referCE_motif_list']))]
referCE_seq = [mat_file['referCE_seq'][i][0][0][0][1][0] for i in range(len(mat_file['referCE_seq']))]

loop_group_ref_seq = [mat_file['loop_group_ref_seq']['Sequence']
					  [i][0][0] for i in range(len(mat_file['loop_group_ref_seq']))]
seq_1AMU = mat_file['seq_1AMU']['Sequence'][0][0][0]
S_code_site = mat_file['S_code_site']
loop_range_list = mat_file['loop_range_list']

C_motif_dict = dict(zip(referCE_dtype_str,referCE_motif_list))
C_ref_seq_dict = dict(zip(referCE_dtype_str,referCE_seq))
C_motif_name_dict = dict(zip(referCE_dtype_str,[list(range(1,len(v)+1)) for v in C_motif_dict.values()]))

all_motif_list = {
	'AMP-binding':A_motif_list,
	'PP-binding':T_motif_list,
	'Thioesterase':TE_motif_list,
}
all_motif_list.update(C_motif_dict)

all_ref_seq = {
	'AMP-binding':A_refer_seq,
	'PP-binding':T_refer_seq,
	'Thioesterase':TE_refer_seq,
}
all_ref_seq.update(C_ref_seq_dict)

all_motif_name = {
	'AMP-binding':A_motif_name,
	'PP-binding':T_motif_name,
	'Thioesterase':TE_motif_name,
}
all_motif_name.update(C_motif_name_dict)

pro_matrix = substitution_matrices.load("BLOSUM62")
nuc_matrix = substitution_matrices.load("NUC.4.4")
# 使用EMBOSS里面Needle 和Water 参数
global_aligner = Align.PairwiseAligner()
global_aligner.mode = 'global'
# global_aligner.substitution_matrix = nuc_matrix
global_aligner.open_gap_score = -10
global_aligner.extend_gap_score = -0.5

local_aligner = Align.PairwiseAligner()
local_aligner.mode = 'local'
local_aligner.substitution_matrix = pro_matrix
local_aligner.open_gap_score = -10
local_aligner.extend_gap_score = -0.5

hmmheader = ['target name', 't accession', 'tlen', 'query name', 'q accession', 'qlen', 'E-value', 'full score', 'full bias', '#', 'of', 'c-Evalue', 'i-Evalue', 'domain score', 'domain bias', 'hmm from', 'hmm to', 'ali from', 'ali to', 'env from', 'env to', 'acc', 'description of target']

# 将氨基酸非法字符替换为'X'
for seqrecord in seqrecords:
	if set(seqrecord) - set(AA_leter):
		seq = str(seqrecord.seq)
		for letter in (set(seqrecord) - set(AA_leter)):
			seq = seq.replace(letter, 'X')
		seqrecord.seq = Seq(seq)

# hmm scan by pfam
SeqIO.write(seqrecords, temp_fasta_path, "fasta")

FNULL = open(os.devnull, 'w')
subprocess.run(sub(
	'hmmscan --domtblout '+hmm_out_dom_path+' --noali -E 1e-5 --domE 1e-5 Pfam-A.hmm  ' + temp_fasta_path), shell=True, stdout=FNULL)

# remove head and tail
lines = []
with open(hmm_out_dom_path, 'r') as fp:
	lines = fp.readlines()

with open(hmm_out_txt_path, 'w') as fp:
	for line in lines[3:-10]:
		fp.write(' '.join(re.split("\s+", line)[0:23]) + '\n')

hmmer_raw = pd.read_csv(hmm_out_txt_path, sep='\s+', header=None, names=hmmheader, index_col=False)
hmmer_raw = hmmer_raw.astype({"query name": str})

if len(hmmer_raw) == 0:
	print('No domain found')
	sys.exit()

##################
##################
# Final output file
##################
Result = {}

for seqrecord in seqrecords:
	region = hmmer_raw[hmmer_raw['query name'] == seqrecord.id]

	hmmer_Other_result = region[region['target name'].isin(
		[name for name in NRPS_domains if name != 'Thioesterase'])]
	hmmer_TE_result = region[region['target name'] == 'Thioesterase']
	# print(hmmer_raw)
	TE_over = hmmer_TE_result[hmmer_TE_result['hmm to'] -
							  hmmer_TE_result['hmm from'] > hmmer_TE_result['tlen']*0.5]
	TE_under = hmmer_TE_result[hmmer_TE_result['hmm to'] -
							   hmmer_TE_result['hmm from'] <= hmmer_TE_result['tlen']*0.5]
	Other_over = hmmer_Other_result[hmmer_Other_result['hmm to'] -
									hmmer_Other_result['hmm from'] > hmmer_Other_result['tlen']*0.6]
	Other_under = hmmer_Other_result[hmmer_Other_result['hmm to'] -
									 hmmer_Other_result['hmm from'] <= hmmer_Other_result['tlen']*0.6]

	true_region = pd.concat([Other_over, TE_over])
	pesu_region = pd.concat([Other_under, TE_under])

	start_list = np.array(true_region['env from'])
	end_list = np.array(true_region['env to'])
	domain_list = np.array(true_region['target name'])
	pesu_domain_list = pesu_region['target name']

	# process the problem because hmmer3 sometime will spilt one domain to two same type domains.
	for domainname in pesu_domain_list.unique():
		hmm_domain = pesu_region[pesu_region['target name'] == domainname]
		if len(hmm_domain) > 0:
			remove_list = []
			domain_length = hmm_domain['tlen'].iloc[0]
			for index1, rows1 in hmm_domain.iterrows():
				if index1 not in remove_list:
					for index2, rows2 in hmm_domain.iterrows():
						if (index2 not in remove_list) and (index1 != index2):
							start1 = hmm_domain.loc[index1, 'env from']
							start2 = hmm_domain.loc[index2, 'env from']
							end1 = hmm_domain.loc[index1, 'env to']
							end2 = hmm_domain.loc[index2, 'env to']
							# if overlap or gap <50% domain length
							if (start1 <= start2 and start2 <= end1) or (start1 <= end2 and end2 <= end1) or (end2 <= start1 and ((start1-end2) < 0.5*domain_length)) or (start2 >= end1 and ((start2-end1) < 0.5*domain_length)):
								new_start = min([start1, start2])
								new_end = max([end1, end2])
							else:
								remove_list.append(index1)
								remove_list.append(index2)
								continue
							# merge two picies, looking for more
							if (not (np.logical_and(start_list > new_start, start_list < new_end).any())) and (not (np.logical_and(end_list > new_start, end_list < new_end).any())) and ((new_end-new_start+1) > 0.6*domain_length):
								domain_list = np.append(
									domain_list, pesu_domain_list.loc[index1])
								start_list = np.append(start_list, new_start)
								end_list = np.append(end_list, new_end)
								remove_list.append(index1)
								remove_list.append(index2)
								break

	# sort by order
	list_sorted = pd.DataFrame(np.array(
		[start_list, end_list, domain_list]).T, columns=['start', 'end', 'domain'])
	list_sorted = list_sorted.sort_values(by=['start'])

	list_sorted = list_sorted.reset_index(drop=True)
	list_sorted.loc[-1] = [np.nan, 1, "START"]  # adding a row
	list_sorted.index = list_sorted.index + 1  # shifting index
	list_sorted.loc[len(list_sorted)] = [len(seqrecord), np.nan, "END"]
	list_sorted = list_sorted.sort_index()

	# find motif by reference sequence
	result_table = pd.DataFrame(columns=['start', 'end', 'domain', 'motif_seq',
								'motif_name', 'inter_motif_seq', 'C_subtype', 'C_score','loop_length', 'loop_seq', 'loop_group', 'S_code'])
	last_domain_end = 1
	for i in range(len(list_sorted)):
		if i == 0 or i == len(list_sorted)-1:
			continue
		else:
			domain_name = list_sorted.loc[i]['domain']
			# C this start - next start
			if domain_name == 'Condensation':
				start = list_sorted.loc[i]['start']
				end = list_sorted.loc[i+1]['start']
			# A last end - next start
			elif domain_name == 'AMP-binding':
				start = list_sorted.loc[i-1]['end']
				end = list_sorted.loc[i+1]['start']
			# #T last end - this end
			# elif domain_name == 'PP-binding':
			# 	start = list_sorted.loc[i]['end']
			# 	end = list_sorted.loc[i]['end']
			# TE T this
			else:
				start = list_sorted.loc[i]['start']
				end = list_sorted.loc[i]['end']

		# given seq and domain type, identity motif
		seq = seqrecord[start-1:end].seq

		if domain_name == 'Condensation':
			SeqIO.write(SeqRecord(seq,id = 'temp_C'), temp_fasta_path, "fasta")
			subprocess.run(sub(
				'hmmscan --domtblout '+hmm_out_dom_path+' --noali -E 1e-5 --domE 1e-5 C_model.hmm  ' + temp_fasta_path), shell=True, stdout=FNULL)
			# remove head and tail
			lines = []
			with open(hmm_out_dom_path, 'r') as fp:
				lines = fp.readlines()
			with open(hmm_out_txt_path, 'w') as fp:
				for line in lines[3:-10]:
					fp.write(' '.join(re.split("\s+", line)[0:23]) + '\n')
			hmmer_raw = pd.read_csv(hmm_out_txt_path, sep='\s+', header=None, names=hmmheader, index_col=False)
			hmmer_raw = hmmer_raw.astype({"query name": str})

			domain_name = hmmer_raw.loc[0]['target name']
			list_sorted.loc[i]['domain'] = domain_name

		# look for loop
		loc_loop_length = []
		loc_loop_seq = []
		loc_S_code = []

		if domain_name == 'AMP-binding':
			loc_seq_struct = SeqRecord(seq, id='input')
			A_refer_seq_ = SeqRecord(Seq(A_refer_seq), id='A_refer_seq')
			seq_1AMU_ = SeqRecord(Seq(seq_1AMU), id='seq_1AMU')
			SeqIO.write([loc_seq_struct, A_refer_seq_, seq_1AMU_] + [SeqRecord(Seq(seq), id='loop_group_ref_seq' + str(i))
						for i, seq in enumerate(loop_group_ref_seq)], temp_fasta_path, "fasta")
			subprocess.run(
				sub('./clustal -i '+temp_fasta_path+' -o '+temp_aln_fasta_path+' --force'), shell=True)
			loc_loop_seq_MSA = [seq for seq in SeqIO.parse(
				sub(temp_aln_fasta_path), "fasta")]
			loc_loop_range = np.zeros((len(loop_range_list), 1))

			for j in range(len(loop_range_list)):
				loc_loop_range[j] = Codetransform_HRL(
					loc_loop_seq_MSA[1].seq, loop_range_list[j, 1])

			loc_loop_length = np.zeros(len(loop_range_list)-1)

			for j in range(len(loop_range_list)-1):
				loc_seq1 = str(loc_loop_seq_MSA[0][int(
					loc_loop_range[j]-1):int(loc_loop_range[j+1]-1)].seq).replace('-', '')
				loc_loop_length[j] = len(loc_seq1)
				loc_loop_seq.append(loc_seq1)

			loc_index1 = np.zeros(len(S_code_site))
			for j in range(len(S_code_site)):
				loc_index1[j] = Codetransform_HRL(
					loc_loop_seq_MSA[2].seq, S_code_site[j])
			loc_S_code = [str(loc_loop_seq_MSA[0].seq)[int(idx)-1]
						  for idx in loc_index1]
		
		# print(domain_name)
		# print(all_motif_name[domain_name])
		# print(all_ref_seq[domain_name])
		# print(all_motif_list[domain_name])
		motifs = find_motif(
			seq, all_ref_seq[domain_name], all_motif_list[domain_name], list_sorted, i, start)
		motifs = np.array(motifs).reshape(-1, 2)
		motifs_name = all_motif_name[domain_name]
		loop_group = Loop_length2group_HRL([loc_loop_length], mat_file)

		if (domain_name) == 'E':
			domain_name = 'Epimerization'

		if (domain_name) in C_motif_dict:
			domain_name = 'Condensation'
			C_subtype = hmmer_raw.loc[0]['target name']
			C_score = hmmer_raw.loc[0]['full score']
		else:
			C_subtype = '-'
			C_score = '-'

		for i, motif_pos in enumerate(motifs):
			if i == 0:
				if np.isnan(motifs.flatten()[0]):
					temp_result = [last_domain_end, motifs.flatten(
					)[0]-1, domain_name, '', 'inter', '', C_subtype, C_score, loc_loop_length, loc_loop_seq, loop_group, loc_S_code]
				else:
					temp_result = [last_domain_end, motifs.flatten()[0]-1, domain_name, '', 'inter', str(
						seqrecord[:int(motifs.flatten()[0])-1].seq), C_subtype, C_score, loc_loop_length, loc_loop_seq, loop_group, loc_S_code]
				result_table.loc[len(result_table.index)] = temp_result

			# inter 不包前后
			if i in list(range(1, len(motifs))):
				j = i*2
				if np.isnan(motifs.flatten()[j-1]) or np.isnan(motifs.flatten()[j]):
					temp_result = [motifs.flatten()[j-1]+1, motifs.flatten()[j]-1, domain_name,
								   '', 'inter', '', C_subtype, C_score, loc_loop_length, loc_loop_seq, loop_group, loc_S_code]
				else:
					temp_result = [motifs.flatten()[j-1]+1, motifs.flatten()[j]-1, domain_name, '', 'inter', str(seqrecord[int(
						motifs.flatten()[j-1]):int(motifs.flatten()[j])-1].seq), C_subtype, C_score, loc_loop_length, loc_loop_seq, loop_group, loc_S_code]

				result_table.loc[len(result_table.index)] = temp_result

			if np.isnan(motif_pos[0]):
				temp_result = [motif_pos[0], motif_pos[1], domain_name, '', motifs_name[i],
							   '', C_subtype, C_score, loc_loop_length, loc_loop_seq, loop_group, loc_S_code]
			else:
				temp_result = [motif_pos[0], motif_pos[1], domain_name, str(seqrecord[int(motif_pos[0])-1:int(
					motif_pos[1])].seq), motifs_name[i], '', C_subtype, C_score, loc_loop_length, loc_loop_seq, loop_group, loc_S_code]

				result_table.loc[len(result_table.index)] = temp_result

			if i == len(motifs)-1:
				last_domain_end = motifs.flatten()[i*2+1]+1

	test_df = result_table.copy(deep=True)
	index_ = -1
	flag = 0
	for index, row in test_df.iterrows():
		# 处理识别类型
		if row['motif_name'] == 'G':
			if not G_judge:
				row['start'] = np.nan
				row['end'] = np.nan
		elif row['motif_name'] == 'alpha' and row['domain'] == 'AMP-binding':
			if not A_judge:
				row['start'] = np.nan
				row['end'] = np.nan
		elif row['motif_name'] == 'alpha' and row['domain'] == 'PP-binding':
			if not T_judge:
				row['start'] = np.nan
				row['end'] = np.nan

		# 处理nan
		if np.isnan(row['start']) and np.isnan(row['end']):
			# 就单独一个
			if flag == 0:
				test_df.loc[index-1, 'end'] = test_df.loc[index+1, 'end']
				try:
					test_df.loc[index-1, 'inter_motif_seq'] = str(seqrecord[int(
						test_df.loc[index-1, 'start'])-1:int(test_df.loc[index+1, 'end'])].seq)
					test_df.drop([index], inplace=True)
					test_df.drop([index+1], inplace=True)
				# 后有NAN（前有的话flag = 1）
				except ValueError:
					flag = 1
					index_ = index-1
					test_df.drop([index], inplace=True)
			elif flag == 1:
				test_df.drop([index], inplace=True)
		elif np.isnan(row['end']):
			if flag == 0:
				flag = 1
				index_ = index
			elif flag == 1:
				test_df.drop([index], inplace=True)
		elif np.isnan(row['start']):
			test_df.loc[index_, 'end'] = row['end']
			test_df.loc[index_, 'inter_motif_seq'] = str(seqrecord[int(
				test_df.loc[index_, 'start'])-1:int(test_df.loc[index_, 'end'])].seq)
			test_df.drop([index], inplace=True)
			flag = 0

	Result["Seq " + seqrecord.id] = test_df.T.to_json()

outputdic["Result data"] = True
outputdic["Data json"] = Result
outputdic["Process id"] = processid
outputjson = json5.dumps(outputdic)
print(outputjson)
```
