## Supplementary material for "Knowledge-guided data mining on the standardized architecture of NRPS: subtypes, novel motifs, and sequence entanglements": NRPS Motif Finder online version code (Python): Readme.txt

Usage: python Find_motif_HRL_0207.py -h
python Find_motif_HRL.py -i <inputfile> -o <outputfile> -p <processid> -G <1 or 0> -A <1 or 0> -T <1 or 0>
-i <inputfile> is sequence file in FASTA format
-o <outputfile> is the prefix of result file
-p <processid> is a number to distingish the process from different users
-G <1 or 0> is to activate detecting new motif G-motif in A domain during analysis, 1 = activate, 0 = inactivate
-A <1 or 0> is to activate detecting new motif Aalpha1 motif in A domain during analysis, 1 = activate, 0 = inactivate
-T <1 or 0> is to activate detecting new motif Talpha1 motif in T domain during analysis, 1 = activate, 0 = inactivate
