## Supplementary figures and images for "Knowledge-guided data mining on the standardized architecture of NRPS: subtypes, novel motifs, and sequence entanglements"

### C_all_trim_out6_MSA_trim.fasta.treefile.pdf

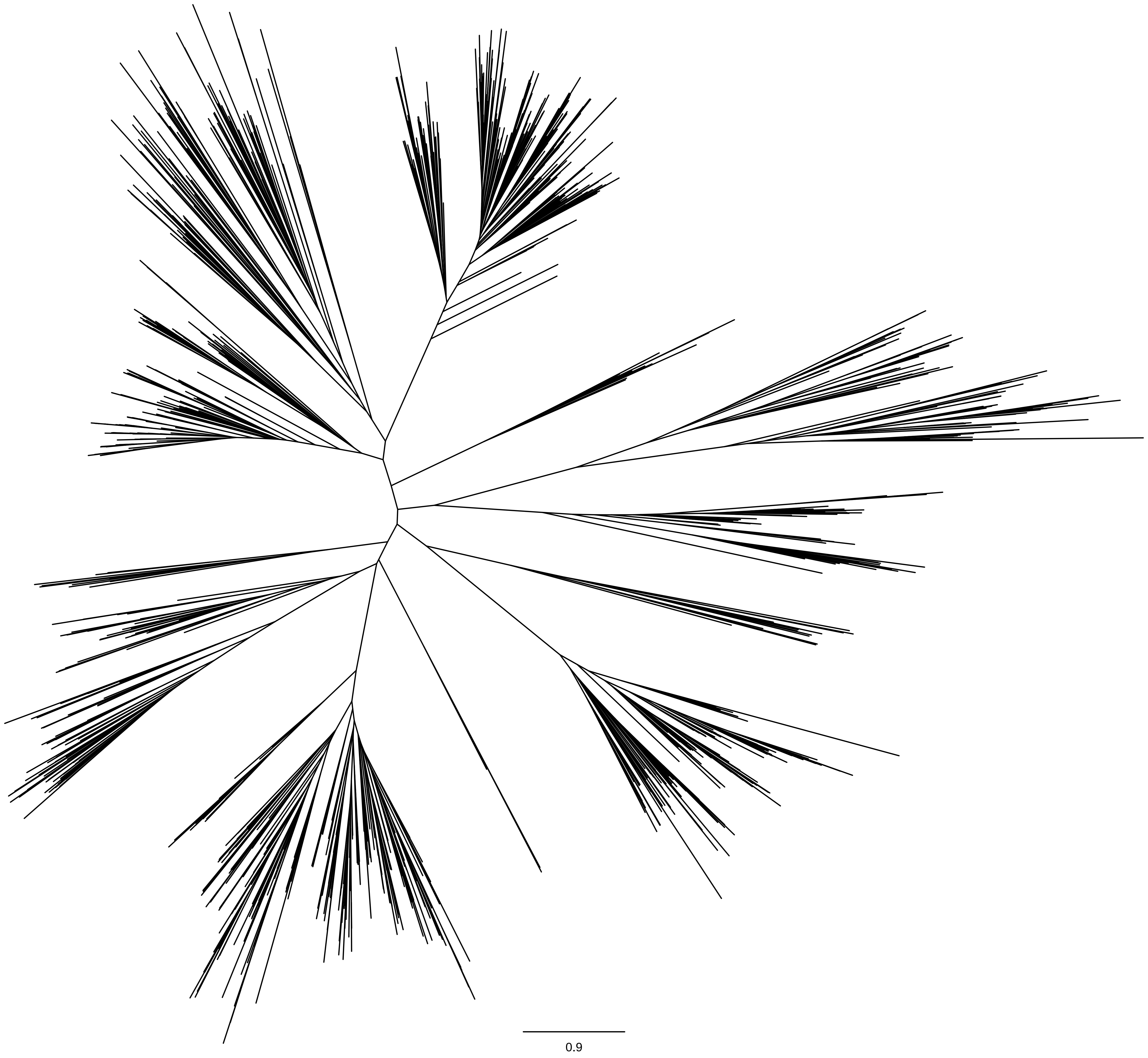
